# A unifying theory of receptive field heterogeneity predicts hippocampal spatial tuning

**DOI:** 10.1101/2025.07.26.666958

**Authors:** Zach Cohen, Jan Drugowitsch

## Abstract

Neural populations exhibit receptive fields that vary in their sizes and shapes. Despite the prevalence of such tuning heterogeneity, we lack a unified theory of its computational benefits. Here, we present a framework that unifies and extends previous theories, finding that receptive field heterogeneity generally increases the information encoded in population activity. The information gain depends on heterogeneity in receptive field size, shape, and on the dimensionality of the encoded quantity. For populations encoding two-dimensional quantities, such as place cells encoding allocentric spatial position, our theory predicts that both size and shape receptive field heterogeneity are necessary to induce information gain, whereas size heterogeneity alone is insufficient. We thus turned to CA1 hippocampal activity to test our theoretical predictions—in particular, to measure shape heterogeneity, which has previously received little attention. To overcome limitations of traditional methods for estimating place cell tuning, we developed a fully probabilistic approach for measuring size and shape heterogeneity, in which tuning estimates were strategically weighted by explicitly measured uncertainty arising from biased or incomplete traversals of the environment. Our method furnished evidence that hippocampal receptive fields indeed exhibit strong degrees of size and shape heterogeneity, abiding by the normative predictions of our theory. Overall, our work makes novel predictions about the relative benefits of receptive field heterogeneities beyond our application to place cells, and provides a principled technique for testing them.

## Introduction

Neural populations exhibit heterogeneity in their selectivity to input stimuli [1–4] and to more latent quantities [5], such as allocentric spatial position [6–10]. This heterogeneity is not only reflected in the stimulus values to which neurons in these populations are most selective, but also in the variety of sizes and shapes of their receptive fields. For example, some CA1 hippocampal place cells, which exhibit spatially localized receptive fields [11], respond only over narrow areas a few centimeters in size, while others are active over much larger areas spanning several meters [6, 8–10]. Heterogeneity in receptive field size has also been observed in heading direction neurons in mice [5] and in orientation-selective visual neurons in macaques [1]. Significant work has aimed to provide a theoretical account for the purpose of heterogeneity in neural tuning.

Previous work has investigated tuning heterogeneity from the perspective of efficient coding [12–17] — how neural tuning needs to be shaped to best encode important stimuli. This framework captures stimulus-driven receptive field heterogeneity, positing that it emerges as a consequence of shifting neural resources towards encoding more likely stimuli [18–20]. However, heterogeneously sized receptive fields also occur when stimulus patterns may have less of an impact on neural tuning, such as when all stimuli occur equally often [6, 8, 9]. Previous work has demonstrated the coding benefit of non stimulus-driven receptive field size heterogeneity only in specific instances, such as for orientation selective neurons in macaque visual cortex [1, 21], or place fields in rodent [8] or bat [10] hippocampus. Whether these insights generalize to populations encoding other quantities remains unknown.

In this study, we introduce a unified framework of the computational benefits of heterogeneity of receptive field size and shape. Specifically, we pursued the hypothesis that receptive field heterogeneity is a general coding principle that increases information about quantities encoded in population activity. Previous work has asked whether neurons should exhibit narrow or broad selectivity in order to maximize information encoded in population activity [22, 23], but primarily studied this question in model populations where all neurons shared the same receptive field sizes and shapes. We instead developed a framework that relates the amount of information encoded in population activity to the degree of receptive field heterogeneity. Our framework develops this relationship for encoded quantities of arbitrary dimensionality, such as orientation tuning (one-dimensional, e.g., [1, 21]), grounded allocentric spatial position (two-dimensional, e.g., [6, 8]), flying allocentric spatial position (three-dimensional, e.g., [10]), and so on. For simplicity, we will refer to all of these quantities as *stimuli*, even if they are latent or abstract quantities, and their dimensionality as the *stimulus dimensionality*. We found that heterogeneity in receptive field sizes across a population broadly increases encoded information as compared to populations without such heterogeneity, and that the magnitude of the information gain is strongly dependent upon the stimulus dimensionality. Intriguingly, we found that for populations encoding two- and three-dimensional stimuli, heterogeneous tuning only increases information if there is also heterogeneity in the shape of each neuron’s receptive field. For populations encoding quantities of higher dimensionality, size heterogeneity across the population remains advantageous, but shape heterogeneity becomes detrimental.

We then asked whether the activity of rat hippocampal CA1 neurons, which putatively encode two-dimensional allocentric spatial position, follows the predictions of our framework. While it is known that their receptive fields vary in size [6–10], heterogeneity of their shapes has been relatively understudied. In analyzing these populations, we found that shortcomings in accounting for noise and bias inherent in measuring receptive fields entailed in traditional approaches made them unsuitable for robustly estimating the degree of place field size and shape heterogeneity. To overcome these issues, we developed a fully probabilistic method for neural tuning estimation, which permits a principled and strategic use of uncertainty to account for noisy and incomplete data. This new approach furnished evidence that place fields indeed exhibit strong degrees of both tuning size and shape heterogeneity, suggesting that rat CA1 neurons exhibit a code predicted by our framework to maximize encoded information about spatial position.

## Results

### The information gain from size heterogeneity depends on stimulus dimensionality

How does heterogeneous tuning across neurons in a population influence how much information is encoded in population activity? We studied encoded information across populations that shared the same tuning statistics but whose individual neurons varied in their tuning properties. This allowed us to understand how tuning statistics on average affect encoded information, rather than focusing on a single model population with fixed tuning parameters. We studied this question in idealized populations of independent, Poisson spiking neurons with bell-shaped, localized tuning curves that uniformly tile the space of possible stimuli (Fig. 1a, Methods M1.1; see Supplementary Information [SI] for an extension to multimodal tuning curves, Extended Data Fig. 1). Their spiking activity encodes a *D*-dimensional quantity which can be an input stimulus, such as the orientation of a visual grating [1], or a latent variable, such as an animal’s allocentric spatial position [6, 10]. We quantified the encoded information,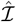, as the population Fisher information averaged across model populations, normalized by population size and stimulus dimensionality (Methods M1.2). The encoded information measures how well population activity supports discriminating close-by stimulus values (Fig. 1c). It scales inversely with the smallest possible decoding error of any unbiased decoder (Fig. 1c), which is achievable by a biologically plausible locally linear decoder (SI 2.1).

**Figure 1:**
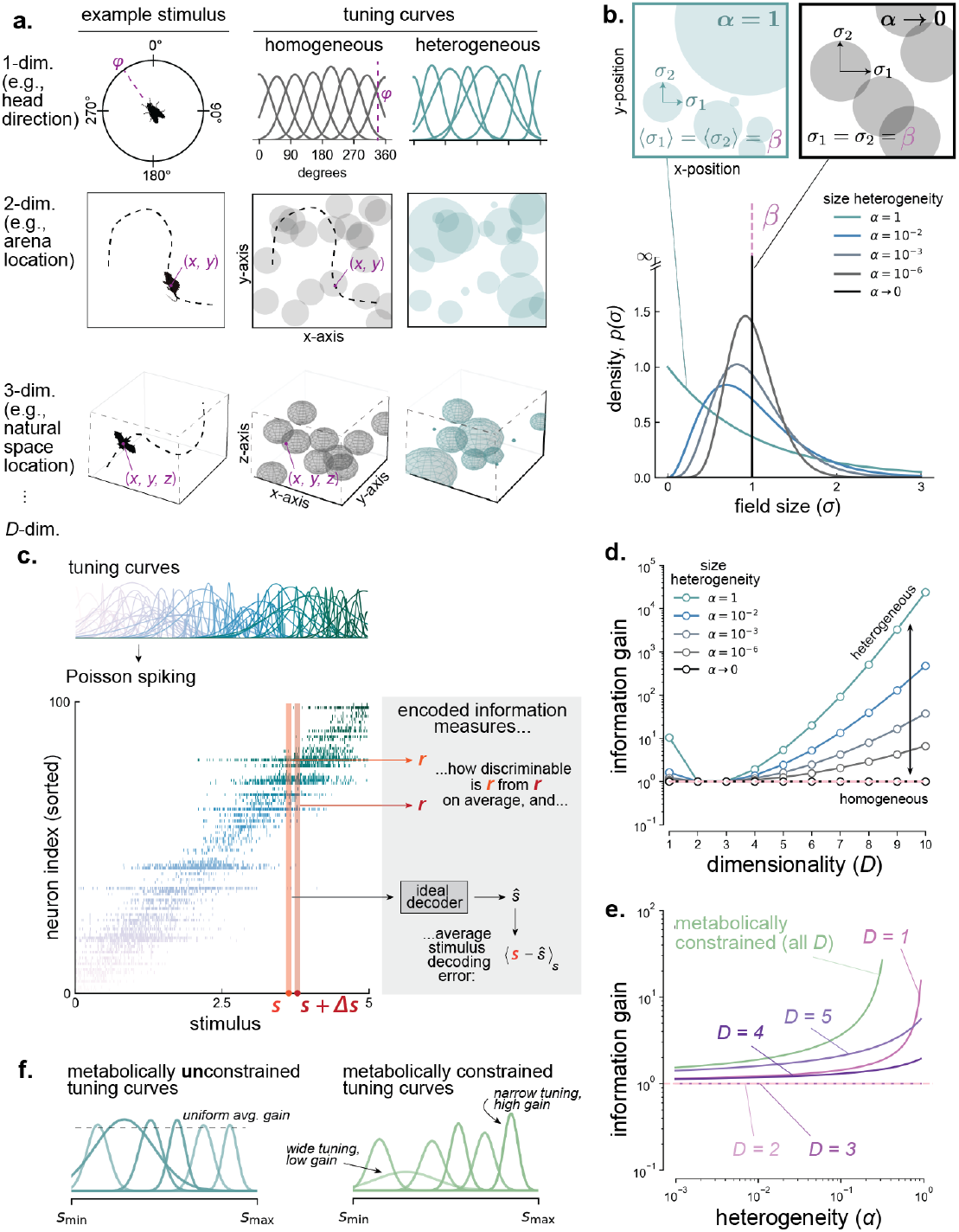
Receptive field size heterogeneity confers a dimensionality-dependent information gain. **a**. Example stimuli of different dimensionalities, and schematics of tuning curves over these stimuli with either homogeneously sized (left) or heterogeneously sized (right) receptive fields. The tuning curves completely covered the stimulus space, but we here only illustrated a subset of neurons to avoid clutter. **b**. The amount of field size heterogeneity in the population is determined by the heterogeneity parameter, *α* (*α* → 0: homogeneous; *α* = 1: maximally heterogeneous), which only impacts field size heterogeneity, but not its mean, *β*. To isolate the effect of size heterogeneity, we assume here that widths along each dimension for any given neuron (*σ*_1_, *σ*_2_) are equal, but allow these widths to vary between neurons in the population. **c**. The encoded information jointly measures how well population activity supports discrimination of similar inputs, and also inversely lower bounds the decoding error of an ideal decoder (which we show is a simple linear decoder). **d**. The information gain (ratio of encoded information in a population with heterogeneously sized fields, *α >* 0, and in a population with homogeneously sized fields, *α* → 0) as a function of the stimulus dimensionality *D* (color = heterogeneity *α*). **e**. The information gain as a function of the degree of field size heterogeneity (color = stimulus dimensionality *D*; green = metabolically constraint population, independent of *D*). **f**. Schematics depicting different assumptions about metabolic expenditure; either there are no constraints placed on the amplitudes of tuning curves (left panel, all lines in **d**. and lines with dimensionality labels in **e**.) or differences in metabolic expenditures as a result of varying field sizes are accounted for through amplitude constraints (right panel, green line in **e**.).

To determine how the heterogeneity of receptive field sizes across neurons in a population modulates encoded information, we derived closed-form expressions (verified through simulation, Extended Data Fig. 2, Methods M2) for the encoded information as a function of receptive field size heterogeneity, *α* (Fig. 1b). At lowest heterogeneity (*α* → 0), all receptive fields have the same size. At highest heterogeneity (*α* = 1), receptive field sizes vary maximally across neurons. To isolate the impact of field size heterogeneity on information, we focused primarily on the information gain 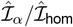 that a population with heterogeneous field sizes (heterogeneity *α*) and encoded information 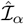 provides over a population with equally sized receptive fields (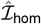; heterogeneity *α* → 0), while keeping the average field size in the heterogeneous population, *β*, the same as the fixed size in the homogeneous population (Fig. 1b).

We found that the information gain due to heterogeneity in receptive field sizes depends strongly on the dimensionality of the encoded stimulus (Fig. 1d). Any amount of heterogeneity confers an information gain for populations encoding either one-dimensional stimuli or stimuli of dimensionality greater than three, with greater heterogeneity imparting a commensurately higher information gain (Fig. 1e). In contrast, for populations encoding two- or three-dimensional stimuli, heterogeneous receptive field sizes confer no information gain over a comparably tuned population with equally-sized receptive fields (Fig. 1d). While these results were derived under the assumption of very large populations, we confirmed in simulations that they also apply to neural populations of moderate sizes (Extended Data Fig. 2).

Our analytic results lend intuition to these findings. The total encoded information equals the geometric mean of the information conveyed about each individual encoded dimension. Thus, we can for illustration focus on just a single dimension *i*. For an individual neuron, *n*, we showed that in general, encoded information along this dimension depends on the receptive field widths by

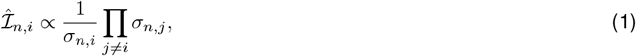

where *σ*_*n,i*_ is the field width for neuron *n* along the *i*-th encoded dimension, and the product is over the field widths, *σ*_*n,j*_, in the remaining encoded dimensions. This expression suggests that information along the *i*-th dimension is maximized when the width of a neuron’s receptive field along this dimension is narrowed, and the widths along remaining dimensions are broadened. A consequence is that the neuron’s activity is maximally sensitive to changes in the stimulus along the *i*-th dimension, and minimally sensitive to changes along remaining dimensions.

The normalized information that the population encodes about stimulus dimension *i* is the information averaged across neurons,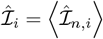. To isolate the effect of size heterogeneity, we first assumed that all neurons shared the same circular shape (concretely, the widths along each dimension were equal: *σ*_*n,i*_ = *σ*_*n,j*_ = *σ*_*n*_), but allowed these widths to vary between neurons in the population. Then, averaged across neurons, Eq. (1) becomes

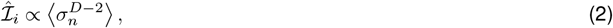

for general dimensionality *D*. Equation (2) tells us that a consequence of assuming that widths along all dimensions are identical is that narrowing (broadening) along dimension *i* leads to commensurate narrowing (broadening) along the alternative dimensions. For two-dimensional stimuli (*D* = 2), this means that any change in information due to narrowing (or broadening) along dimension *i* is canceled out by equivalent narrowing (or broadening) along the alternative dimension *j*. Therefore, the encoded information for any neuron in the population does not depend on the size of its receptive field. Hence, variations in field sizes across the population have no impact on encoded information.

For three-dimensional stimuli (*D* = 3), narrowing (broadening) along dimension *i* again cancels with narrowing (broadening) along one of the alternative dimensions, *j*, such that the encoded information for neuron *n* scales in proportion to the field width *σ*_*n,k*_ along the remaining dimension 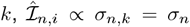. The population’s encoded information is thus 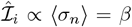. Accordingly, encoded information grows with the average size of the receptive fields, but heterogeneity in field size has no additional impact on information. Therefore, a population with heterogeneously sized fields achieves no information gain over a homogeneously tuned population that matches the average field size.

For higher-dimensional stimuli (*D >* 3), Eq. (2) tells us that the encoded information scales with higher powers of the field size. Populations with homogeneous field sizes thus encode information proportional to powers of this size. When this field size varies across neurons in the population, the higher-order moments of the distribution of field sizes additionally contribute to the encoded information (Fig. 1b). The associated information gain is due to stochastic variation in the field sizes causing some of these fields to be, by chance, much larger—an effect that grows with the magnitude of size heterogeneity in the population (*α* → 1). Therefore, when encoding higher-dimensional stimuli, populations with field size heterogeneity always encode more information than populations with equally-sized receptive fields.

Interestingly, the information gain ceases to depend on stimulus dimensionality once we assume that neurons modulate their peak activity to equalize their metabolic resource use across neurons (Fig. 1e). This causes neurons with larger receptive field sizes to have a lower peak firing rate (Fig. 1f, left vs. right). As a consequence, information gain becomes independent of stimulus dimensionality, while it continues to grow with field size heterogeneity, as before. Subsequent data analysis (see below) did not support signatures of these metabolic constraints in rodent place cells (Extended Data Fig. 3). Thus, for the remainder of the text we focus on the case without metabolic constraints.

### Receptive field shape heterogeneity increases encoded information for two- and three-dimensional stimuli

Our results show that receptive field size heterogeneity increases information in populations encoding one-dimensional and greater-than-three-dimensional stimuli. This provides a normative explanation for the observed size heterogeneity in neural populations encoding one-dimensional stimuli such as head direction [5]. However, our results also show that heterogeneity in receptive field size has no benefit for populations encoding two- or three-dimensional stimuli, which is curious, given that receptive field heterogeneity has been observed in these populations [9, 10]. Thus, we next asked how another receptive field characteristic—namely the “shape” of the receptive fields—may additionally control the population information.

Recall that we have thus far assumed that all receptive fields shared the same, circular shape. We next derived the encoded information for populations in which receptive fields varied in shape. We modeled shape heterogeneity by assuming that field widths are drawn independently in each dimension from the size heterogeneity distribution (Fig. 2a). This causes receptive fields to become on average more ellipsoidal, rather than perfectly spherical (Fig. 2a,b). This seemingly minor change had a profound impact on encoded information. In particular, we found that adding shape heterogeneity to size heterogeneity induces information gain in populations encoding two- and three-dimensional stimuli (Fig. 2c), unlike in the above case where shape heterogeneity is absent. Notably, however, shape heterogeneity is not a panacea. While heterogeneity (of both size and shape) is always beneficial, the information gain from shape heterogeneity in addition to size heterogeneity does not increase with stimulus dimensionality (Fig. 2d, left). In contrast, the information gain from size heterogeneity alone *does* (Fig. 1d). Consequently, size heterogeneity *without* shape heterogeneity emerges as the preferable coding strategy in higher dimensions (Fig. 2c).

**Figure 2:**
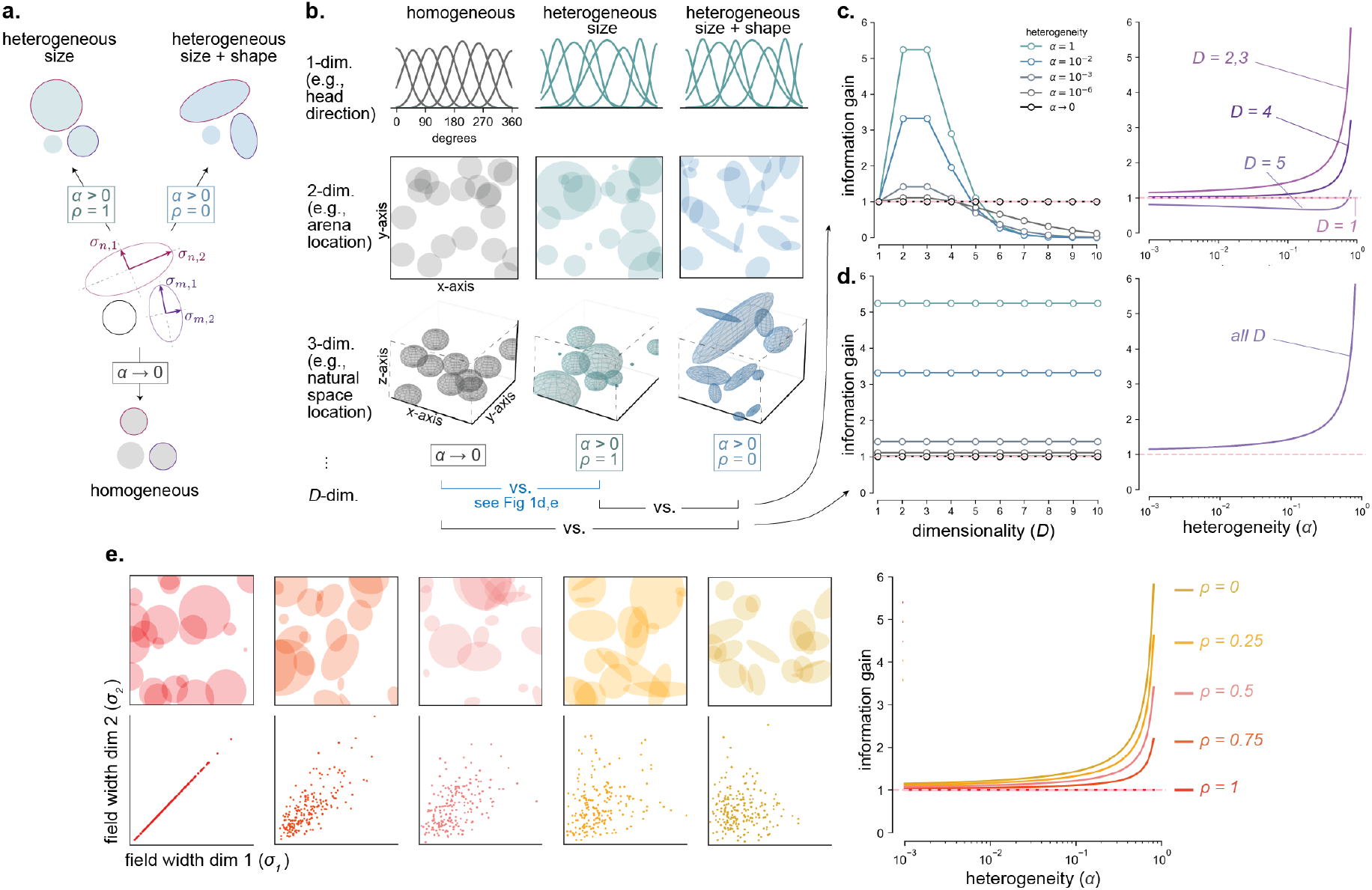
Introducing shape heterogeneity leads to an information gain for lower dimensional stimuli that vanishes for higher dimensional stimuli. **a**. Schematic demonstrating how the parameters governing size heterogeneity (*α* ∈ (0, 1]) and shape heterogeneity (for general dimensionality, *ρ* ∈ {0, 1}), yield populations with different properties. When *ρ* = 0, the principal widths (*σ*_*n*,1_, *σ*_*n*,2_) of a neuron *n*’s receptive field are not necessarily the same, leading to shape heterogeneity. **b**. Example stimuli and example tuning curves that exhibit homogeneously sized receptive fields (left), heterogeneously sized fields (middle) and heterogeneously sized and shaped fields (right). As in **Fig. 1a** we only show a subset of neurons to avoid clutter. **c**. The information gain of size and shape heterogeneity over size heterogeneity alone for different stimulus dimensionalities (left; color = size heterogeneity, *α*) and as a function of the degree of size heterogeneity (right; color = stimulus dimensionality, *D*). **d**. Same as **c**., but information gain of both size and shape heterogeneity over a completely homogeneous population. **e**. The joint impact of size and shape heterogeneity for populations encoding two-dimensional stimuli. For such populations, we derived the information gain as a smooth function of the degree of shape heterogeneity, *ρ* ∈ [0, 1]; resulting degrees of shape heterogeneity depicted in the left 5 panels, colors correspond to the line labels in the right-most plot depicting information gain. The *ρ* = 0 case equals the *D* = 2, 3 case in panel **c** (right). The *ρ* = 1 case equals the *D* = 2 case in **Fig. 1e**.

To better understand these results, recall that Eq. (1) shows that information about the *i*-th encoded dimension is larger when the receptive field width is narrow along that dimension and broad along the remaining dimensions *j* ≠ *i*. When field shapes are heterogeneous, the field widths of neuron *n, σ*_*n,i*_ *& σ*_*n,j*_’s vary independently, so that Eq. (1) becomes on average,

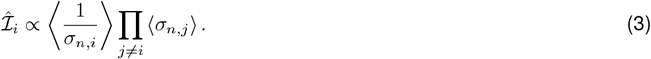

This decouples broadening and narrowing across alternative dimensions, such that the associated consequences no longer cancel each other out. For two- and three-dimensional stimuli this explains the additional information gain due to shape heterogeneity in populations with heterogeneous receptive field sizes (Fig. 2c, *D* = 2, 3).

However, for higher dimensions, the impact of shape heterogeneity on information gain reverses. In this case, only variability in the field width *σ*_*n,i*_ along the *i*-th dimension in Eq. (3) contributes to information gain. All other dimensions only contribute information in proportion to the average field width ⟨*σ*_*n,j*_ ⟩ (i.e., its first moment), just like in populations with homogeneous field widths. This has two consequences. First, the information gain due to size and shape heterogeneity does not depend on stimulus dimensionality, as only a single dimension contributes to this gain (Fig. 2d, left). Second, shape heterogeneity poses a disadvantage when compared to size heterogeneity alone, as in the latter case, higher-order moments of the field size distribution can provide an additional information gain (Fig. 2c, larger *D*).

Given the special relationship between width and shape heterogeneity for populations encoding two-dimensional stimuli, we further characterized how receptive field shape impacts encoded information in this case. To do so, we introduced a new parameter, *ρ* ∈ [0, 1], that quantifies how strongly the two field widths are correlated, and that allowed us to continuously move between homogeneous (*ρ* = 1) and completely heterogeneous (*ρ* = 0) receptive field shapes (Fig. 2e). We found that stimulus information increases inversely linearly as field shapes become increasingly heterogeneous (*ρ* → 0). Further, for any degree of field shape heterogeneity, the stimulus information increases as field sizes become more heterogeneous (*α* → 1) (Fig. 2e). Importantly, given empirical measurements of 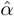 and 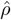 for a population of neurons encoding two-dimensional stimuli, this result allows us to precisely quantify the information gain of the population over comparably tuned homogeneous populations.

Crucially, our results reveal the yet un-characterized role of receptive field shape in population coding: For two- and three-dimensional stimuli, mere field size heterogeneity across the population does not confer any representational advantage. Indeed, field size heterogeneity is only beneficial in these regimes if receptive field shapes are also heterogeneous.

### Tuning curve estimates based on the spike-triggered average are sensitive to bias

Our theoretical findings predict that populations encoding two-dimensional variables can increase encoded information if their receptive fields vary in both sizes and shapes. This motivated us to next ask if there is evidence for such heterogeneity in data. Specifically, we studied the population activity of rat hippocampal place cells, which putatively encode two-dimensional allocentric position. While the distribution of place field sizes in rat CA1 neurons has been characterized before [6–9], their field shape heterogeneity has not yet been quantified. We thus sought to measure the degree of field size 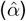 and shape 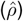 heterogeneity from rat CA1 neural data recorded as rats engaged in a free foraging task [24].

Most existing methods for measuring place cell tuning in two-dimensional arenas rely on computing the spike triggered average (STA) over uniform spatial bins covering the arena (Fig. 3a) [8, 9]. We found that this approach easily succumbs to issues arising from limited or incomplete sampling, leading to biased estimates of 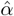 and 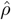.

**Figure 3:**
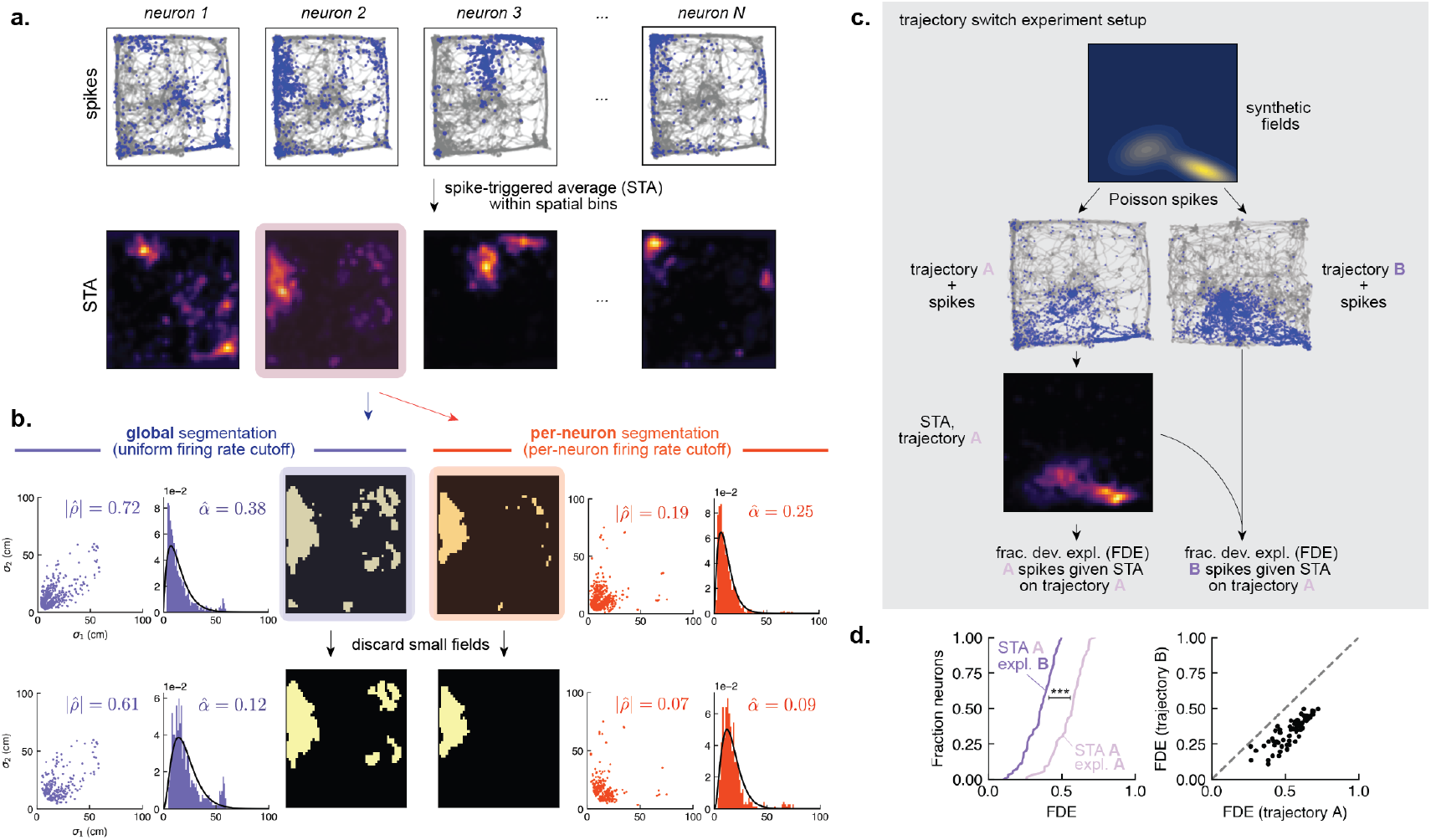
Place field tuning estimation with the STA is biased by heuristic noise filtering algorithms and by trajectories taken by the animal. **a**. Traditional place field estimation starts by computing the binned STA (top: gray lines = behavioral trajectories, blue dots = spikes; bottom: associated STAs). **b**. STAs are turned into rate maps, that are then segmented into subfields to estimate their widths. Varying hyperparameter choices for each step can yield different field width estimates that bias estimates of size heterogeneity (*α*, right column on either side) and shape heterogeneity (*ρ*, left column on either side). Common choices are either uniform rate and size cutoffs [7, 8] (left), or per-neuron rate and size cutoffs [9] (right). **c**. A schematic of the trajectory switch experiment. The STA obtained from one real trajectory (trajectory A) on synthetically generated fields and spikes achieves a lower fraction of deviance explained (FDE) on spike sequences generated from a different trajectory (trajectory B); see Methods M4.2 for details. **d**. The FDE for trajectory B is on average 35.89% lower than that for trajectory A (paired t-test, *p* = 1.95 *×* 10^−37^), as illustrated by the CDF across neurons (left) and by a neuron-by-neuron comparison (right; dots = neurons), indicating that the STA provides biased receptive field estimates.

First, limited sampling makes it challenging to distinguish between signal and noise. The STA furnishes a single point estimate of the average firing rate at each (binned) location in the environment. These estimates can be noisy in locations where data is limited, such as in locations that the rodent rarely visits. Segmentation algorithms that extract field shape and size from the STA must filter out this noise. However, this is often done heuristically. For example, segmentation algorithms that measure field size by counting contiguous bins in which a neuron is active often discard spatial bins in which spiking activity falls below a chosen firing rate threshold (Fig. 3b). More generally, any form of heuristic noise filtration can introduce measurement biases. Indeed, when applied to CA1 spiking data, we found that different, commonly used hyperparameter choices in such noise filtering algorithms (e.g., [6, 8, 9]) can produce vastly different measurements of both field size heterogeneity and field shape heterogeneity in the population (Fig. 3b). Importantly, it is difficult, if not impossible, to ascertain the “ideal” choice of hyperparameters in lieu of a ground truth.

Estimates based on the STA are also sensitive to incomplete sampling—in this case, non-uniform traversals of the environment by the rat, which leave some areas of the environment unvisited. To demonstrate this, we quantified how well the STA computed from one traversal pattern can explain spikes observed during another traversal pattern (Methods M4.2). We did so by generating spike sequences from simulated neurons with synthetic place fields while using two different real trajectories of a rat (trajectories A & B) over the simulated fields (Fig. 3c). We in turn used the spikes for trajectory A only to compute the STA, and quantified how well the STA explains the spike sequences for both trajectories by the fraction deviance explained (FDE, Methods M4.1). If the STA were an unbiased estimate of the true underlying simulated fields, we would only expect insignificant differences in the FDE between the two trajectories. However, we indeed find a significant and substantial drop in the FDE between explaining spikes from trajectory A and trajectory B (Fig. 3d; 35.89% drop in deviance explained, paired t-test test, *p* = 1.95 *×* 10^−37^). This suggests that the STA reflects the animal’s biased occupancy of the environment, and thus may fail to accurately measure the true tuning properties of a neuron.

In short, the standard method for measuring place cell tuning is problematic. It is impacted by the animal’s non-uniform occupancy of the environment. As a result, it produces biased estimates of place cell tuning. To better estimate place fields—and to test our theory—a different approach is needed.

### A fully probabilistic approach to tuning curve estimation

The biases inherent in STA-based approaches largely stem from failing to account for uncertainty in the firing rate estimates arising from limited and biased sampling of the environment. We hypothesized that explicitly tracking and making strategic use of this uncertainty could mitigate these biases. This, in turn, would permit more reliable measurements of the quantities of interest (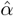 and 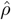). Specifically, explicitly accounting for uncertainty unlocks a principled approach to handling noise, where observations (in this case, field widths) are weighted by their associated uncertainties.

To accomplish this, we took a Bayesian approach to characterize the full probability distribution over tuning curve parameters given the observed spikes of hippocampal place cells. To support place cells with multiple place fields [7], we designed a generative model in which each neuron in a population can exhibit multimodal tuning curves with an arbitrary number of subfields (Fig. 4a), which jointly describe the firing rate at any location in the arena. This firing rate in turn determines a neuron’s spiking activity through a Poisson process. Finally, we used Markov chain Monte Carlo (MCMC) to invert the generative model and estimate a posterior distribution over the number of subfields and the parameters of each subfield (width, gain, center, etc.) for each recorded neuron given its spiking information and the rat’s trajectory (Fig. 4b; Methods M3).

**Figure 4:**
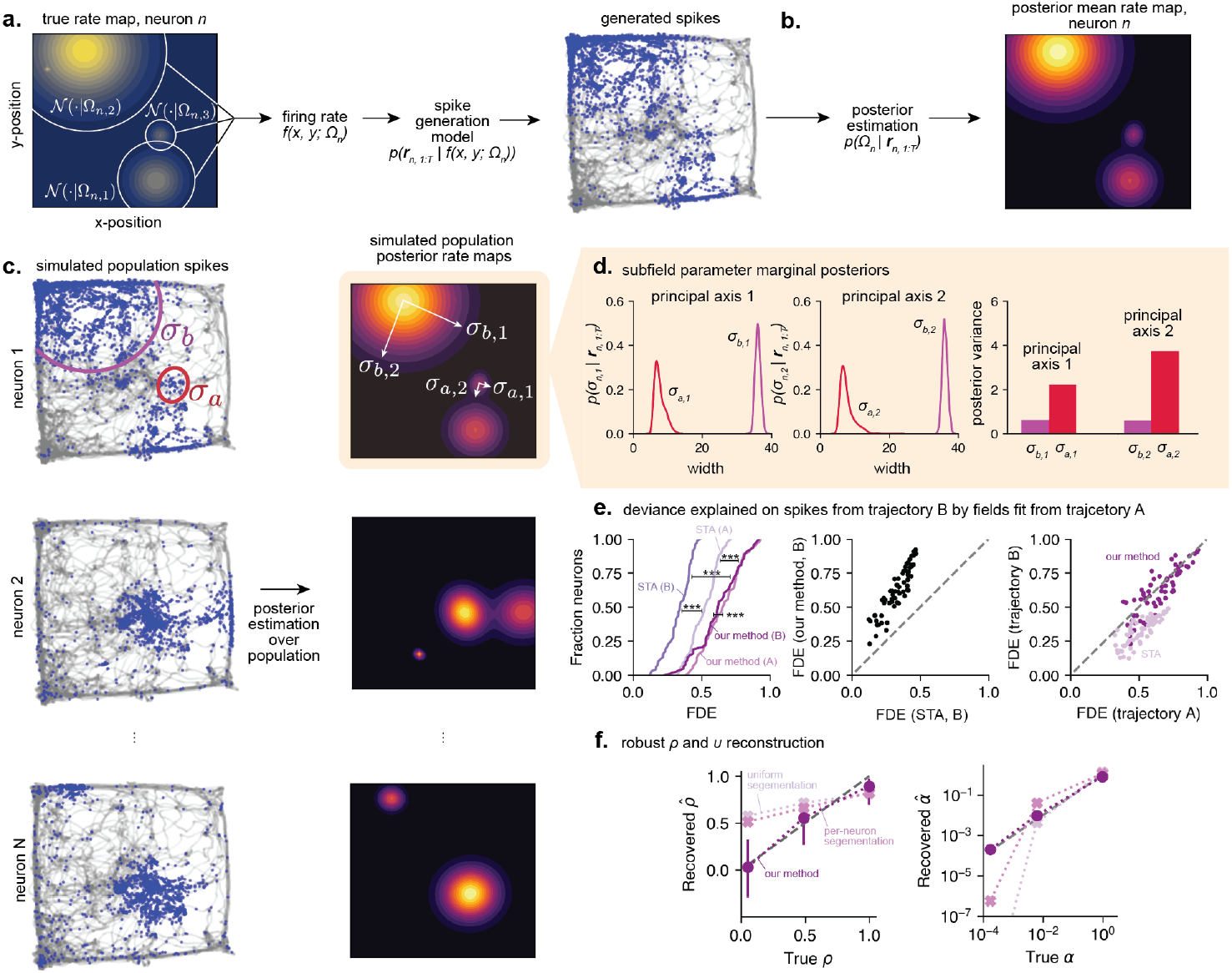
Bayesian place field parameter estimation permits strategic use of uncertainty for robust estimation of size and shape heterogeneity in a population. **a**. In our generative model, the firing rate of a neuron *n* at location (*x, y*) is given by the collective activations of bell-shaped subfields, each with its own set of parameters, which together comprise the total set of parameters for neuron *n* (here, {Ω_*n*_ = Ω_*n*,1_, Ω_*n*,2_, Ω_*n*,3_}). Spikes are generated by a Poisson process. **b**. Inverting this generative model upon observing spikes and location sequences yields a posterior distribution over a neuron’s rate map and subfield parameters. **c**. Example posterior rate maps (right) obtained using synthetically generated fields and spike sequences over these fields obtained from real rat trajectories (left). **d**. The marginal posterior variance of each subfield’s parameters (left and middle plots) reflects the relative information available to explain each subfield. The trajectory and spike information about *σ*_*a*_ is smaller compared to *σ*_*b*_ (**c**.), leading to higher uncertainty about estimating *σ*_*a*_ versus *σ*_*b*_, which is reflected in the larger posterior variance for *σ*_*a*_ versus *σ*_*b*_ (right). **e**. Revisiting the trajectory switch experiment (**Fig. 3b**) with our method. This method incurs only a minor, 4.52% drop in FDE for spikes from trajectory B vs. A (left: CDF; right: per-neuron comparison in dark magenta; paired t-test, Bonferroni corrected *p* = 0.015), whereas using the STA results in a 35.83% drop (left: CDF; right: per-neuron comparison in pale pink; paired t-test, Bonferroni corrected *p* = 3.98 *×* 10^−38^). The STA consistently resulted in lower FDEs than our method (left: CDFs; middle: per-neuron comparison for trajectory B; trajectory A: 21.91% drop in deviance explained, paired t-test, Bonferroni corrected *p* = 7.18 *×* 10^−33^; trajectory B: 47.51% drop in deviance explained, paired t-test, Bonferroni corrected *p* = 1.86 *×* 10^−35^). **f**. Our method correctly estimates shape (*ρ*) and size (*α*) heterogeneity in synthetic populations with known parameters (dark magenta; error bars = 95% confidence interval) whereas STA with different segmentation hyperparameters yields biased estimates (pale pink: uniform segmentation, pink: per-neuron segmentation).

To ensure that our approach led to accurate characterization of receptive field properties, we initially tested it on a simulated dataset where the ground-truth subfield parameters are known (Fig. 4c). We generated simulated spiking sequences over the synthetic fields using real trajectories taken by a rat [24]. To study the method’s effectiveness under conditions that mimic its later application to real data, we chose place field parameters that statistically matched those of empirical data (Methods M3.5, Extended Data Fig. 4). This revealed several advantages of our method over the STA-based approach.

First, our method explicitly quantifies uncertainty in the estimated field parameters. We can see this by examining the marginal posterior distributions over subfield widths estimated by the Bayesian approach on simulated data (Fig. 4d). In areas of the environment with dense traversal patterns and many spikes, (over subfield *σ*_*b*_ in Fig. 4c-d) the width of the marginal posterior (i.e., its variance) is much smaller than in areas with sparser traversal patterns and fewer spikes (over, for example, subfield *σ*_*a*_ in Fig. 4c-d). In other words, the greater amount of spiking (and traversal) data available for subfield *σ*_*b*_ permits greater certainty about our estimate of its properties than about those of subfield *σ*_*a*_, for which there is less data available. This means that for each subfield of each neuron, we can measure not only the subfield width, but also quantify how certain we are of the measured width.

Second, our method was less impacted by incomplete sampling of the environment as a result of biased rat trajectories. To show this, we applied the same trajectory switch experiment from the previous section (Fig. 3c), asking how well the subfield parameters inferred from spiking data on trajectory A can explain spikes observed on trajectory B. We found that, unlike the STA (Fig. 4e; 35.83% drop in deviance explained, Bonferroni corrected *p* = 3.98 *×* 10^−38^), the inferred subfield parameters using our Bayesian method can explain spikes from trajectory A and B nearly equally well (Fig. 4e; 4.52% drop in deviance explained, paired t-test, Bonferroni corrected *p* = 0.015). In fact, our method appears to better explain spiking data than the STA regardless of the trajectory (Fig. 4e; STA A vs. our method A: 21.91% drop in deviance explained, paired t-test, Bonferroni corrected *p* = 7.18 *×* 10^−33^, STA B vs. our method B: 47.51% drop in deviance explained, paired t-test, Bonferroni corrected *p* = 1.86 *×* 10^−35^), though this result is expected for simulated data that exactly matches the assumed generative model of the Bayesian approach.

Lastly, our method more robustly measures the quantities of interest to our theory, 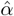 and 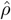. To show this, we generated simulated spiking sequences over real rodent trajectories from synthetic populations with varying degrees of shape (*ρ*) and size (*α*) heterogeneity while keeping all other parameters fixed. We then applied our method to these spike sequences to infer the tuning parameters of each of the stimulated neurons. Finally, we estimated 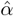 and 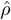 from the maximum a-posteriori (MAP) estimates of the field widths while weighting the individual estimates in proportion to their certainty (see Methods M3.5). We found that our method accurately recovered the desired ground-truth parameters (Fig. 4f), and outperformed reconstruction of these parameters based on the STA (Fig. 4f). While noise manifesting from spike variability and incomplete sampling appeared to bias estimates of the degrees of shape and size heterogeneity based on the STA, this was not the case for our method. Indeed, our method instead took into account uncertainty induced by such noise, and weighted estimates appropriately, mitigating bias in turn.

Taken together, our results show that our Bayesian approach to tuning curve parameter measurement permits robust measurement of the key quantities that our theory predicts control encoded information. The robustness conferred by the Bayesian approach follows directly from the explicit quantification and strategic use of uncertainty.

### Hippocampal CA1 receptive fields exhibit near optimal width and shape heterogeneity

Finally, we used our method to measure the empirical 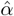 and 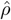 in CA1 spiking data ([24]) as rats freely foraged around a 2×2m arena. We restricted our analysis to recording periods in which the animal was moving, to remove hippocampal activity associated with non-local activations that arise during periods of stillness, such as sharp-wave ripples [24, 25]. We applied our measurement technique to 8 individual sessions (2 sessions per 4 rats), and recovered place fields that qualitatively matched the STA (Fig. 5a). In fact, our method explained roughly 90-95% of the fraction of deviance of the spiking data as place fields recovered by STA (Fig. 5b; percent change in deviance explained inset, paired t-test *p*-values reported in figure legend; but see also Extended Data Fig. 5, which details how modest hyperparameter choices in computing the STA can lead to our method better explaining data across all sessions), indicating that our generative model is a valid characterization of place field activity and provided a good fit to the observed spiking data.

**Figure 5:**
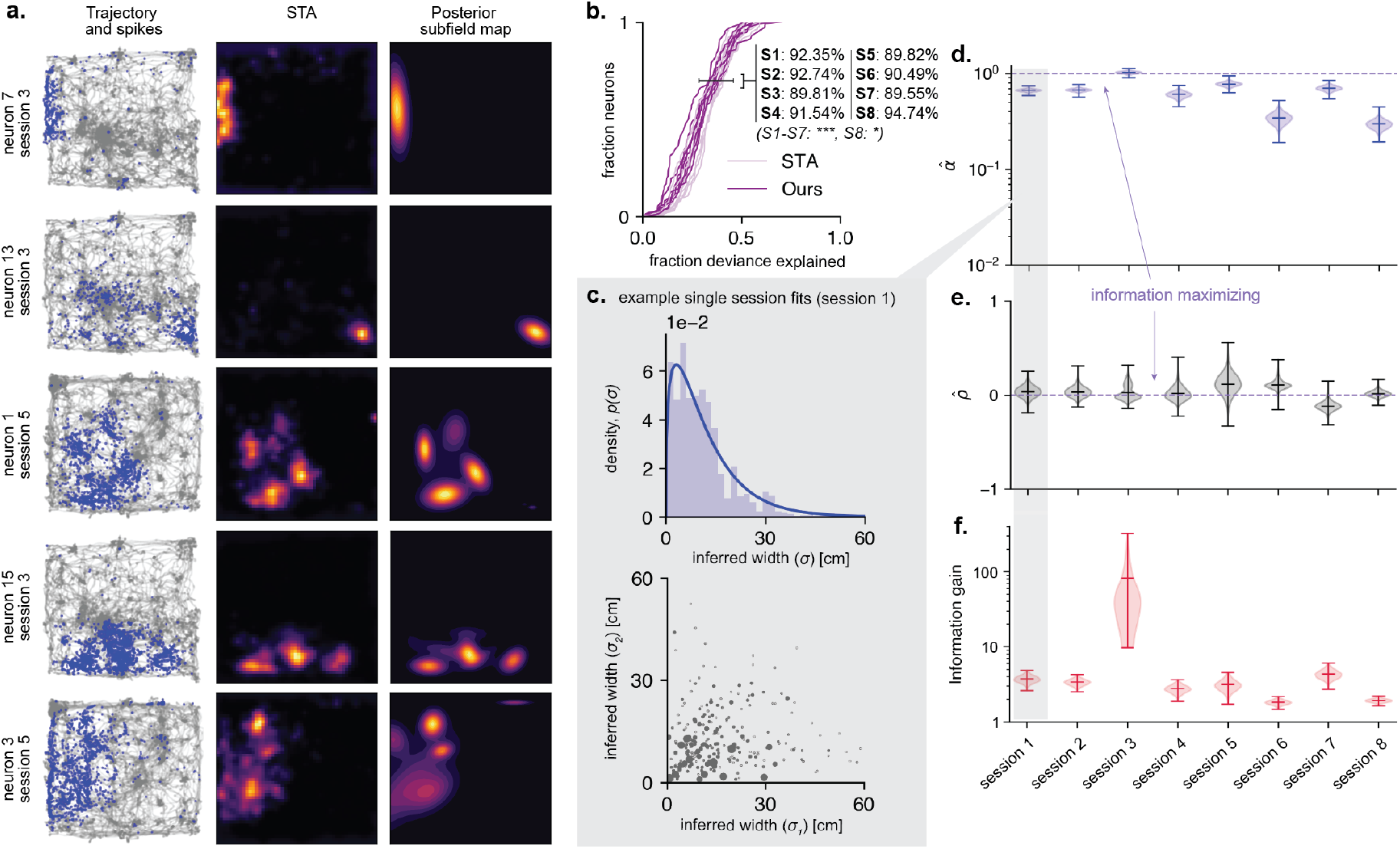
Application of our method to CA1 hippocampal spike sequences reveals information maximizing degree of shape and size heterogeneity. **a**. Example spike and trajectory sequences (left), and rate maps recovered by the STA (middle) and by our inference method (right); five example neurons from across 8 possible sessions. **b**. Comparison of FDE CDFs between the STA and our method. Our inference method explains 89-95% of the deviance explained by the STA (percentages reported in figure inset, paired t-tests: S1: *p* = 4.75 *×* 10^−24^, S2: *p* = 6.52 *×* 10^−54^, S3: *p* = 9.42 *×* 10^−27^, S4: *p* = 3.5 *×* 10^−12^, S5: *p* = 2.24 *×* 10^−21^, S6: *p* = 3.32 *×* 10^−16^, S7: *p* = 4.25 *×* 10^−19^, S8: *p* = 0.04), suggesting our generative model is valid to describe place field activity. **c**. Example recovery of the degree of size (top) and shape (bottom) heterogeneity in a single run session. The sizing of dots in the bottom plot indicates the degree of certainty about the estimated principal width, as reflected in the variance of its corresponding marginal posterior (**Fig. 4d**.). **d**. Bootstrapped distributions of the degree of size heterogeneity in a population across all 8 sessions, which clustered near the value predicted to maximize information (*α* = 1). **e**. Bootstrapped distributions of the degree of shape heterogeneity in a population across all 8 sessions, which clustered near the value predicted to maximize information (*ρ* = 0). **f**. The distribution over information gain over a homogeneously tuned population expected given the distributions over empirical size and shape heterogeneity (**d**,**e**, respectively). Center lines and error bars in **d**.-**f**. show means and maximum/minimum values, respectively.

We then used the fitted tuning curve parameters to measure the empirical 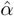 and 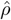 across all 8 sessions. As above, we weighted estimates of neurons’ subfield parameters by the associated certainty reported in the marginal posterior variance for each parameter. This results in measurements of 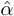 and 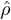 in Fig. 5d-e, with a single example session inset in Fig. 5c (see Extended Data Fig. 4 for remaining sessions).

We found that the estimated distributions of the degree of shape 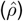 and size 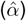 heterogeneity cluster near the values that are predicted by our theory to maximize information gain in populations encoding two-dimensional quantities. For this we relied on an extension of our theory that supports multiple subfields (see SI 2.2.1, 3.2). As for single subfields, the theory predicts that populations encoding two-dimensional stimuli can achieve maximum information if their receptive field shapes are broadly heterogeneous (with 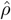 near 0) and with receptive field heterogeneity being maximal (with 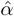 close to 1), in line with the parameters we recovered (Fig. 5d,e). Notably, in our theory we only considered positive shape heterogeneity parameters, *ρ* ∈ [0, 1]. However, our measurement procedure also supports identifying that subfield principal widths are anti-correlated 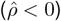. Nonetheless, we find that even when allowing for 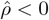, across sessions, 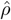 is near 0.

When we applied our theory to these recovered parameters, we found that the combined size and shape heterogeneity at least doubled the encoded information, and for some populations increased it 4- or even 10-fold (Fig. 5f). This suggests that place cells appear to adopt the normative coding approach predicted by our model to increase encoded spatial information available for downstream populations.

Previous work has documented receptive field distortions near the boundaries of explored environments [26], wherein place fields close to boundaries appear to be more ellipsoidal. To assess whether the shape heterogeneity that we measured is primarily due to these effects, we asked if subfields became more circular with increasing distance from environmental boundaries. While we observed a mild dependence between boundary distance and subfield circularity, this dependence was only statistically significant for one of the 8 sessions we analyzed (Extended Data Fig. 6), suggesting that shape heterogeneity is persistent in subfields across the arena. Furthermore, previous work found that task demands (such as reward location) can lead to place fields non-uniformly covering the environment [7, 27, 28]. In the dataset we analyzed, we found that subfield centers were roughly uniformly distributed across both spatial dimensions of the arena (Extended Data Fig. 4c & d), in line with the assumptions underlying our theory. This result is expected, given that the rodents were engaged in a free foraging task in which rewards were randomly distributed across the environment.

## Discussion

Neural populations across the brain exhibit heterogeneously sized and shaped receptive fields [1–10]. Using an analytically tractable model, we studied under which conditions the heterogeneity in receptive field sizes and shapes leads to increased information encoded in neural populations. Interestingly, for encoding stimuli of one or greater-than-three dimensions, we found that size heterogeneity alone is sufficient to increase information. For two- and three-dimensional stimuli, in contrast, information only increases if receptive fields vary in both size *and* shape. Furthermore, for encoding stimuli of dimensionality higher than three, shape heterogeneity becomes harmful rather than helpful.

Having identified two-dimensional encoding as a special case of our theory, we turned to test it in spiking data of rat CA1 hippocampal neurons that putatively encode two-dimensional spatial location. When doing so, we found that existing STA-based methods for measuring receptive fields were unsuitable to test our theory as they were sensitive to noise and biases in the collected data. We overcame these issues by developing a fully probabilistic approach for estimating neural tuning, which permits a principled and strategic use of uncertainty to account for noisy and incomplete data. Our new approach furnished evidence that place fields indeed exhibit strong degrees of both tuning size and shape heterogeneity—in line with our theory—to possibly increase available information about spatial position available for downstream populations.

In order to isolate the impact of receptive field size and shape heterogeneity, we primarily focused the information gain arising from this heterogeneity, rather than on the total, normalized encoded information. The latter depends on other tuning characteristics, such as the average field size, the average gain in the population, as well as other tuning properties (see Methods M1.3 and SI 3,4) that have in part been quantified elsewhere [22]. The total, unnormalized Fisher information furthermore scales with population size and the stimulus dimensionality. We applied normalizing transformations to account for such dependence, which allowed us to compare the information gain across stimulus dimensionalities and population sizes.

Our work builds on, and significantly generalizes previous work demonstrating that a specific type of field size heterogeneity can maximize information in populations encoding two-dimensional stimuli [8]. The authors in [8] focused mostly on size heterogeneity, but in simulations with two-dimensional tuning curves varied both size and shape to observe an information gain. As we have shown here, both types of heterogeneity are required in two dimensions, but might be harmful for other stimulus dimensionalities. Indeed, our analytic approach allowed us to disentangle the impact of size and shape heterogeneity on populations encoding two-dimensional quantities, which has not been directly characterized before. In the SI we discuss how our theory relates to that of Zhang et al. in more detail (SI 6). Lastly, without our probabilistic approach to estimate tuning curve properties from noisy and biased neural data, Zhang et al. could only speculate on the actual distribution over field sizes—a speculation that we confirmed with our probabilistic approach.

Recent work used a model different from ours in describing the heterogeneity of field size and shape of neurons encoding one- to three-dimensional stimuli [5, 29]. They described neural tuning by a Gaussian process over these stimuli, followed by a threshold above which neurons become active [5, 29]. This allowed them to capture various tuning statistics, including the distributions of field sizes, of the number of fields, and of the gaps between them. For analytical tractability, our theory made slightly different assumptions about the distribution of field sizes. Nonetheless, as we have shown, our qualitative findings do not depend on these assumptions (SI 3). Therefore, while the information gain for Gaussian process-based models might differ quantitatively, our main findings for the impact of size and shape heterogeneity should also apply to them.

In pursuit of a theory that is relevant beyond its application here to place cells, we studied receptive field heterogeneity that is largely stochastic and unstructured. Specialized populations in the brain exhibit more structured heterogeneity. For example, grid cells exhibit periodic receptive fields that are organized into discrete modules of differently sized periods. A substantial body of literature has demonstrated how such structured and modular heterogeneity is useful, particularly in populations with periodic tuning curves [30–33]. Future work could be dedicated to unifying our results with existing computational accounts of grid cell tuning, probing the relative benefits of heterogeneity in structured and unstructured codes.

While we restrict the application of our tuning curve measurement analysis tool to estimating place field properties, we expect its application to be beneficial more generally. Indeed, explicitly accounting for uncertainty in measurements of tuning characteristics can help overcome many of the difficulties associated with estimating tuning properties with the STA alone. That said, unlike computing the STA, our probabilistic approach is time- and compute-intensive. While this did not prohibit our application, future work could be dedicated to improving the algorithmic efficiency of our probabilistic approach.

Our work focused on information maximization, but additional objectives may drive tuning heterogeneity. Indeed, recent theoretical work has argued that place field tuning heterogeneity emerges as a solution for learning to navigate the environment to maximize reward [34]. This study can be read as an account for how stimulus-driven heterogeneity emerges through learning. Future work could postulate how *non* stimulus-driven heterogeneity can similarly be shaped through learning. Other recent work has demonstrated that tuning heterogeneity can increase the representational capacity of neural populations, allowing populations to encode more quantities in more easily linearly separable patterns of neural activity [35]. Information theoretic measures such as ours have often been used to study populations close to the sensory periphery, where it seems logical that it is advantageous to encode stimuli with high fidelity [1, 12–14, 21]. There is no reason to presume that the brain abandons efficient codes once we move away from the periphery. Thus, our findings should equally apply to all neural codes, irrespective of whether they encode sensory or latent variables.

Our framework makes testable predictions about the interplay between tuning shape heterogeneity, tuning size heterogeneity, and encoded stimulus dimensionality that can be studied in populations across the brain beyond place cells. Our work also provides a template for testing these predictions in other neural populations, wherein uncertainty manifesting from biased or incomplete sampling can be explicitly accounted for and marshaled to refine estimates of neural tuning.

## Acknowledgements

We thank Brad Pfeiffer and David Foster for sharing their data with us. We thank Rachel Wilson, Christopher Harvey, Albert Lee, Camille Rullán Buxó, and Albert Chen for comments on an earlier version of the manuscript, and the Drugowitsch Lab for feedback on the work. This work was supported by a Kempner Graduate Fellowship (Z.C.) and NIH funding (U19NS118246 & R01NS116753; J.D.).

## Author contributions

Z.C. and J.D. conceived of the study. Z.C. performed the analysis. Z.C. and J.D. interpreted the results. Z.C. and J.D. wrote and revised the manuscript.

## Declarations of competing interests

We have no conflicts of interest to report.

## Code & data availability

Code and data used to produce the manuscript figures will be made available upon acceptance of the manuscript.

## Methods

### M1 Population activity model and population information

#### M1.1 Population activity model and tuning curves

##### Population activity

We assumed a population of *N* independent Poisson-spiking neurons that emits a population vector of spikes 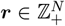 in response to a *D*-dimensional stimulus or latent variable ***s*** ∈ ℝ^*D*^ (for *D* = 2, ***s*** can be interpreted as the spatial position of an animal). Each neuron *n* had tuning curve *f* (***s***; Ω_*n*_), where Ω_*n*_ denotes the neuron’s tuning parameters, such that the likelihood of observation population activity vector ***r*** for stimulus ***s*** is given by

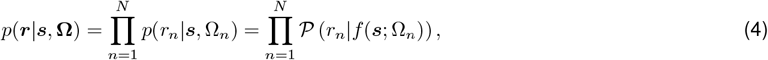

where 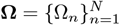 is the set of parameters of all neurons, and 𝒫 (*r*_*n*_ | *f* (***s***; Ω_*n*_)) denotes the Poisson probability of observing *r*_*n*_ spikes for firing rate *f* (***s***; Ω_*n*_).

##### Tuning curves

The tuning curve of each neuron consisted of one or multiple Gaussian-shaped subfields. In general, we assumed that each neuron could have an arbitrary number *P* of subfields (i.e., that their tuning curves were multimodal), such that its tuning is given by

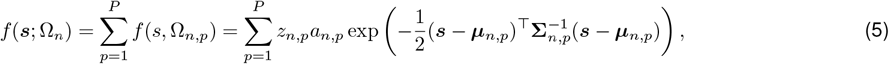

where Ω_*n,p*_ = {*z*_*n,p*_, *a*_*n,p*_, ***µ***_*n,p*_, **Σ**_*n,p*_} is the set of tuning curve parameters for the *p*-th subfield, and 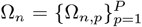. Each subfield *p* has gain *a*_*n,p*_ *≥* 0, tuning center ***µ***_*n,p*_ ∈ ℝ^*D*^, and tuning covariance **Σ**_*n,p*_ ∈ ℝ^*D×D*^. Furthermore, its binary indicator variable *z*_*n,p*_ ∈ {0, 1} determines whether the subfield is active (*z*_*n,p*_ = 1), or inactive (*z*_*n,p*_ = 0), in which case it does not contribute to a neuron’s tuning. We assumed that subfield activations were drawn from a Bernoulli distribution, *z*_*n,p*_ *∼* Bern(*ζ*), such that subfields were active with probability *ζ*. Thus, on average, each neuron has *Pζ* active subfields. For instances in which we only considered unimodal tuning curves, we set *P* = 1 and fix *z*_*n*,1_ = 1, such that each neuron’s tuning curve simplifies to

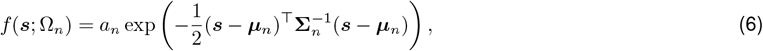

where here, to simplify notation, we have dropped the *p* index.

##### Tuning curve parameters that control size and shape heterogeneity

We assumed that the tuning curve parameters for each subfield of each neuron were independently drawn from the same distribution, Ω _*n,p*_ *∼ p*(Ω _*n,p*_), that is *p*(**Ω**) = Π_*n,p*_ *p*(Ω_*n,p*_). This distribution thus determined the tuning curve statistics of the whole neural population. We here describe how we parameterized the distribution over the tuning covariance, **Σ**_*n,p*_, which is the most critical aspect of our work, as it determines the receptive field size and shape. Details for the remaining subfield parameters can be found in the Supplementary Information.

We assumed that each tuning covariance was given by the diagonal matrix 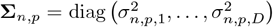 where *σ*_*n,p,i*_ denotes the field width for the *i*th stimulus dimension. The use of diagonal (i.e., axis-aligned) tuning covariances is not a limiting assumption, as for sufficiently large populations, arbitrary rotations of these tuning covariances have no impact on the encoded information (SI 2.2.6).

For homogeneous tuning shapes, we assumed an isotropic covariance, such that *σ*_*n,p,i*_ = *σ*_*n,p*_ for all *i* = 1, …, *D*, leading to circular (or, for *D >* 2, spherical/hyperspherical) tuning curves with width *σ*_*n,p*_. This width is drawn from a Gamma distribution *σ*_*n,p*_ *∼* Γ (*υ, β/υ*), such that its mean, ⟨*σ*_*n,p*_⟩ = *β*, is the same across all neurons and subfields. For heterogeneous tuning shapes, we assumed an anisotropic covariance where the width for each dimension *i* is independently drawn from *σ*_*n,p,i*_ ∼ Γ (*υ, β/υ*). This caused tuning shapes to be mostly elliptical (or, for *D >* 2, ellipsoidal/hyper-ellipsoidal), with mean widths ⟨*σ*_*n,p*,1_⟩ = *β* that are shared across all stimulus dimensions, neurons, and subfields. For *D* = 2 we allowed the field width *σ*_*n,p*,1_ and *σ*_*n,p*,2_ to be correlated by modeling them as a correlated bivariate Gamma distribution [36] with correlation coefficient *ρ* ∈ [0, 1], and where *σ*_1_ and *σ*_2_ share the same marginal distribution. This allowed us to continuously move between the fully heterogeneous case (at *ρ* = 0) and fully homogeneous case (at *ρ* = 1). For *D* = 1, shape heterogeneity does not apply, as tuning curves had a single width *σ*_*n,p*,1_ that was drawn according to *σ*_*n,p*,1_ *∼* Γ (*υ, β/υ*).

For all cases, size heterogeneity is controlled by the shape parameter *υ* of the Gamma distribution that the individual widths are drawn from. Smaller *υ*’s lead to a larger heterogeneity of tuning curve widths. As special cases, *υ* → 1 leads to the *σ*_*n,p,i*_’s to be distributed according to an exponential distribution, whereas, *υ* → *∞* leads to a fixed-width population, where all tuning curves share the same width, *β*. In this limit, the width becomes the same in all dimensions, such that tuning shapes become homogeneous, even if they are drawn from the heterogeneous shape distribution. The choice of a Gamma distribution to model tuning widths in the population is made both to align our work with previous studies [8], and because the Gamma distribution has positive support, and all of its higher-order moments exist, which we find is a necessary condition for computing the FI in closed-form for arbitrary stimulus dimensionality *D*.

In the main text, we express size heterogeneity through a re-parameterization of the Gamma shape parameter, *υ*. Specifically, we introduce, *α* = exp(1 − *v*), so that *α* = 0 corresponds to no size heterogeneity (and is the case of *υ* → *∞*), whereas *α* = 1 corresponds to the maximum degree of size heterogeneity considered in our model (*υ* = 1). While this re-parameterization aids in exposition of the results, we leave the derivations below in terms of *υ*, as this latter parameterization yields more succinct expressions.

##### Metabolic constraints

Neurons with wider tuning curves are active over a wider range of stimuli, and thus use more metabolic resources than more narrowly tuned neurons. We therefore considered two cases. In the metabolically unconstrained case, this extra metabolic expenditure was ignored, and neural gain for all neurons was drawn from *a*_*n*_ *∼* 𝒰 [*g*_mean_ − *G, g*_mean_ + *G*] (where *G* is an arbitrary constant in the range *G* ∈ [0, *g*_mean_] so that ⟨*a*_*n*_⟩ = *g*_mean_), irrespective of the width of their tuning curves. In the metabolically constrained case, their gains were adjusted to meet a fixed metabolic constraint *C* for each stimulus dimension *D*, that is

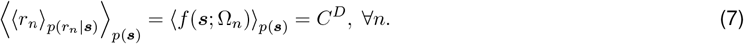

This implies that neural gains scale inversely with tJheir tuning curve widths, that is *a*_*n*_ = |**Σ**_*n*_|^−1*/*2^. To see why this is the case, note that we can rewrite 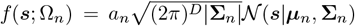. Then, 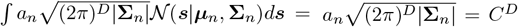, assuming that 𝒩 (***s***|***µ***_*n*_, **Σ**_*n*_) *≈* 0 outside of the considered stimulus range. For this quantity to be constant regardless of tuning curve width, we need to set *a*_*n*_ *∝* |**Σ**_*n*_|^−1*/*2^. As *C* is an arbitrary overall scaling parameter, we chose 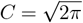 without loss of generality, resulting in *a*_*n*_ = |**Σ**_*n*_|^−1*/*2^.

#### M1.2 Measuring population information from neural population activity

We used Fisher information (FI) to quantify how much population activity ***r*** tells us about the encoded stimulus ***s***. For a population of independently spiking neurons, this FI sums across neurons, and for each neuron depends on its tuning parameters Ω_*n*_. In the large population limit, *N* → *∞*, the information contained in the population approaches *N***ℐ**(***s***), where **ℐ** (***s***) ∈ ℝ^*D×D*^ denotes the per-neuron FI matrix, averaged across possible neural tuning parameters, Ω_*n*_ *∼ p*(Ω_*n*_) (SI 2). We averaged this FI matrix across stimuli, 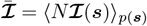, assuming a uniform stimulus distribution, ***s*** *∼* 𝒰 [*s*_min_, *s*_max_]^*D*^. Finally, we summarized it by its determinant, in line with how the determinant of a covariance matrix measures the generalized variance, and re-scaled it by *N* and *D*:

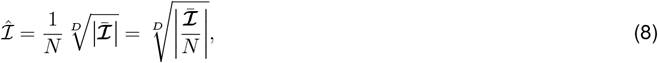

to make our information measure comparable across choices of *N* and *D*, in line with previous literature [8]. This choice of measuring population information captures both the local discriminability of the population code and also provides a lower bounds the error of the best unbiased decoder, which is achieved by a biologically realistic locally linear decoder (SI 2.1). When computing population information, we surveyed the FI in populations that exhibit size and shape heterogeneity under two main conditions: one in which tuning curve centers are unbounded (can exceed the stimulus space by an arbitrarily large extent) and one in which the tuning curve centers of the population are bounded by a finite extent beyond the stimulus space (and, as a special case, exactly bound to the stimulus space). We defer study of the latter case to the Supplementary Information, and present the results in the former case below. We show that the former case can be seen as the asymptotic encoded information, as we found it provides an upper bound on the bounded population information (SI 5).

#### M1.3 A summary of our analytical results

As we show in the Supplementary Information, the encoded information can be found by first computing the FI, averaged across population draws, for one neuron indexed in the population by *n*, **ℐ** _*n*_(***s***; Ω_*n*_), and then applying the transformations detailed in Eq. (8). To obtain the information gain in the various cases described below, we evaluated the encoded information for populations with a given degree of heterogeneity (in either size (*υ*) or shape (*ρ*)) and the encoded information in populations with homogeneous widths (*υ* → *∞*), and computed the ratio between these two expressions. For each case, we start by presenting expressions that relate the encoded information to tuning widths of a neuron, where all other tuning parameters have been marginalized. We then provide expressions for the information gain after marginalizing over the tuning widths under different assumptions of the statistics of the tuning widths (such as the degree of shape or size heterogeneity in the population). The various subscripts on the information,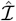, below correspond to the assumptions on the tuning statistics. Detailed derivations for all the following expressions are given in the Supplementary Information.

##### Unimodal tuning curves with no metabolic constraints

We found that, in general, the information of the *i*-th dimension of an encoded quantity for populations with unimodal tuning curves and no metabolic constraints follows

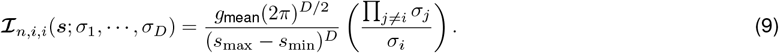

##### Size heterogeneity

When only considering size heterogeneity, taking the average over Eq. (9) and evaluating Eq. (8) yields

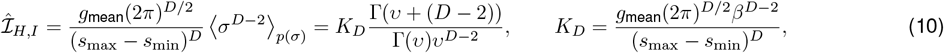

where Γ(·) is the Gamma function, *υ* is the degree of size heterogeneity, *D* is the dimensionality of the encoded quantity, and *K*_*D*_ is a constant that is dimensionality-dependent (hence the ·_*D*_ subscript). The information encoded by a population with homogeneous field sizes is found by taking the *υ* → *∞* limit, and is given by *K*_*D*_. Then, the information gain with size heterogeneity alone is

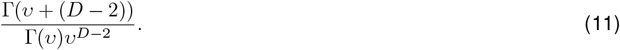

##### Size and maximal shape heterogeneity

When there is both size and shape heterogeneity (ie., *ρ* = 0), averaging over Eq. (9) and evaluating Eq. (8) yields,

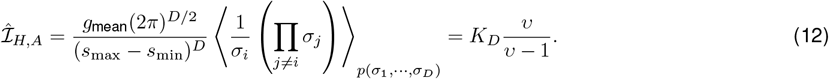

Again, the information encoded by a population with homogeneous field sizes is found by taking the *υ* → *∞* limit and is given by *K*_*D*_, so that the information gain over a homogeneously tuned population is

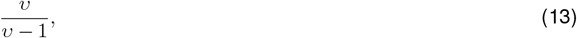

independent of the dimensionality of the encoded quantity.

##### Continuously varying shape heterogeneity for *D* = 2

For general dimensionality, we only considered the extremes of no shape heterogeneity (*ρ* = 1) and maximal shape heterogeneity (*ρ* = 0). For the case where encoded quantities are two-dimensional, we can derive the information gain in terms of a smoothly varying parameter *ρ* ∈ [0, 1] that controls the degree of shape heterogeneity. In this case, *σ*_1_, *σ*_2_ are distributed as a correlated bivariate Gamma distribution with correlation coefficient *ρ* ([36]), so that averaging over Eq. (9) and evaluating Eq. (8) with *D* = 2 yields

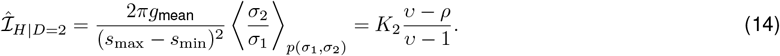

In this case, taking the *υ* → *∞* limit tells us that the information in a population with homogeneous tuning widths is given by *K*_2_, so that the information gain is

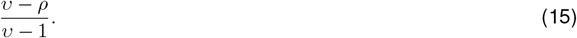

Plugging in *ρ* = 0 recovers Eq. (13), and plugging in *ρ* = 1 recovers Eq. (11) when *D* = 2.

##### Multimodal tuning curves with no metabolic constraints

We found that the encoded information for populations with multimodal tuning curves and no metabolic constraints along the *i*-th encoded dimension follows:

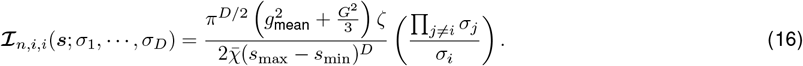

where 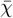 is a constant that captures the average firing rate of a neuron over the arena. The value of this constant depends on the assumptions regarding field size and shape heterogeneity.

##### Size heterogeneity

In the case of size heterogeneity but no shape heterogeneity, averaging over Eq. (16) yields

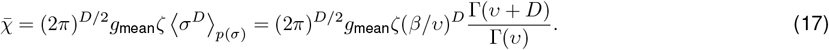

And,

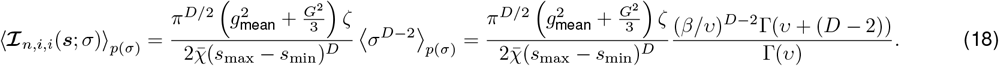

Combining these expressions and evaluating Eq. (8) yields the information:

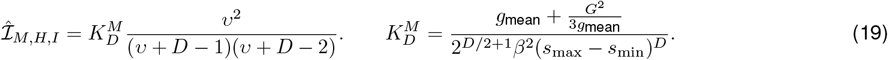

As before, the information gain is obtained by comparing the a fixed width population (*υ* → *∞*), which is 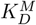, such that the information gain over a population with homogenenous widths is given by,

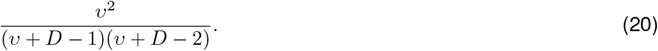

##### Size and maximal shape heterogeneity

In the case of size heterogeneity and maximal shape heterogeneity (*ρ* = 0), averaging over Eq. (16) yields

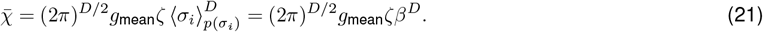

And,

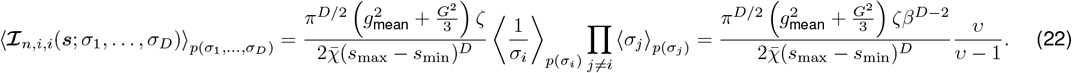

Combining these expressions and evaluating Eq. (8) yields the information:

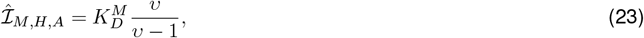

where 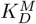 is the same as for the case where only size heterogeneity is assumed. The information in a population with homogeneous widths is again 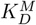, so that the information gain over a population with homogeneous widths in this case is

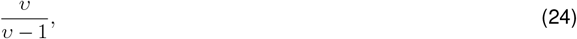

just as in the unimodal case.

##### Continuously varying shape heterogeneity for *D* = 2

As above, we explore continuously varying the shape heterogeneity in the multimodal *D* = 2 case. Here, averaging over Eq. (16) yields

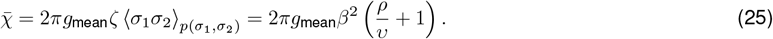

And

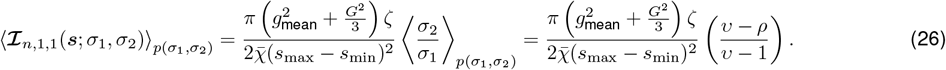

where, as in the unimodal case, *σ*_1_, *σ*_2_ are distributed as a correlated bivariate Gamma distribution with correlation coefficient *ρ*. Combining these expressions and evaluating Eq. (8) yields the information in the *D* = 2 case:

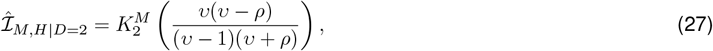

where 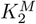 is the constant 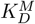 in the case where *D* = 2. The information for a population with homogeneous widths in this case is 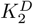, so that the information gain is,

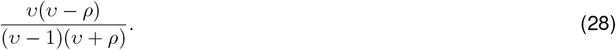

As in the unimodal case, plugging in *ρ* = 0 recovers Eq. (24), and plugging in *ρ* = 1 recovers Eq. (20) when *D* = 2.

##### Unimodal tuning curves with metabolic constraints

For the metabolically constrained case, we restricted our analysis to the unimodal case. Here, the information of the *i*-th dimension of the encoded quantity is given by

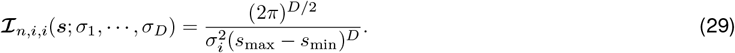

This equation tells us immediately that unlike in the metabolically unconstrained case, shape heterogeneity does not impact encoded information, as the information about the *i*-th dimension of the encoded quantity only depends upon the width of the receptive field along that dimension. Size heterogeneity does matter, however, and the information is obtained by averaging over *p*(*σ*_*i*_) and again evaluating Eq. (8):

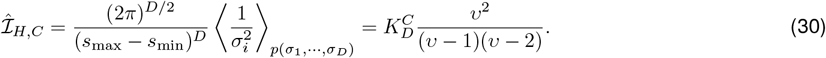

The information in a metabolically constrained population with homogenenous widths is 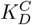, giving an information gain,

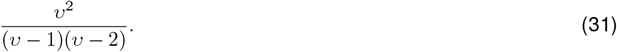

Unlike in the unimodal and multimodal cases without metabolic constraints, the information gain from just size heterogeneity alone is independent of the dimensionality of the encoded quantity.

### M2 Numerical simulations for verifying population information

To verify our analytic results, we performed simulations of neural populations over which we compute the encoded information (Eq. (8)). Given a population with *N* neurons, mean tuning curve size *β*, and heterogeneity *α*, we numerically simulated *M* populations. For each population, the *N* neurons are parameterized by samples from the distributions enumerated above and in the Supplementary Information (ie., gains, tuning curve widths, etc). As in [8], each individual population is presented with a single stimulus ***s***, uniformly drawn from the space of possible stimuli. For each population, we compute the Fisher information of each neuron at that stimulus ***s***, and, as the Fisher information is additive for independently stochastic neurons, we sum the per-neuron FI across all population neurons. Note that in our simulations, we compute the FI matrix derived in Eq. (SI.27) (ie., the expression achieved after analytically averaging over *r*_*n*_ *∼ p*(*r*_*n*_ | ***s***, Ω_*n*_) for a given set of tuning parameters Ω_*n*_ and stimulus ***s***), rather than performing Monte Carlo averaging to approximate this expectation. We previously verified that the Monte Carlo average converges to the analytic expectation for a reasonable number of samples, and opted to use the analytic expression directly, as it drastically reduces computation time of the simulations.

We perform the above procedure for populations of size *N* = 800, with *β*^−1^ = 2 and log *α* ∈ [−4, 0] (Extended Data Fig. 2), or sizes *N* ∈ {50, …, 51200}, over *β*^−1^ ∈ {1, …, 32} and log *α ≈* −2 (Extended Data Figs. 7 & 8), averaged across *M* = 10, 000 populations. For simulations with multimodal tuning curves, we set the maximum number of subfields to be *P* = 20, and set the field rate to be 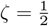, such that on average, each neuron has 10 subfields. For each population, we store the computed FI matrix, across all choices of *N* and *β*. Once we store this set of FI matrices, we can easily compute both our metric and that from [8] in order to verify the analytical relationship we developed between the two measures (Extended Data Fig. 8, SI 6).

The simulation code was implemented in Python 3.8.10, using NumPy 1.17.4 for vectorized computations and SciPy 1.11.1 for special functions. We primarily rely on a vectorized version of the above computations, and we verified through systematic testing that the vectorized and brute force methods produce the same results.

### M3 Estimating field shape and size heterogeneity from spiking data

#### M3.1 Data acquisition

We use data from Pfeiffer & Foster ([24], [37]), where the procedure for data acquisition is described. Animals were trained to forage in a 2 *×* 2m environment. The environment contained 36 wells with food, arranged in a 6 *×* 6 grid. Neural recordings were obtained through a microdrive array containing 40 gold-plated tetrodes implanted in dorsal CA1. Across 4 rats, data was obtained over 2 sessions, each (for a total of 8 sessions). Individual units were classified by [24]. The data was originally supplied in Matlab files. These files were passed through code designed to reformat the data into Python-compatible formats [25]. Units from the original work were classified manually based on spike waveforms [24, 37]. Inhibitory neurons were omitted from our analysis, and were identified mainly based on average firing rate [24, 25, 37].

The animal’s position was determined using head-mounted LED lights that were recorded at 60 Hz by an overhead video system.

#### M3.2 Estimating subfield parameters

Critical to estimating field size heterogeneity (the empirical 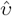) and shape heterogeneity (the empirical 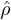) is accurately measuring the (sub)field widths of neurons from spiking data. We formalized this measurement process as a Bayesian inference problem. For a single neuron, we are given {(***s***_1_, *r*_1_), …, (***s***_*T*_, *r*_*T*_)} where ***s***_*i*_ ∈ ℝ^2^ is an animal’s position at time *i* and *r*_*i*_ ∈ ℤ_+_ is the number of spikes emitted by the neuron at time *i*. Then our aim was to compute the parameter posterior

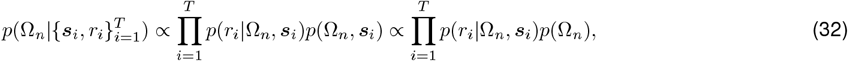

where the last proportionality arises from assuming that Ω_*n*_ and *s*_*i*_ are independent of each other.

#### M3.3 Generative model

Our generative model was designed to reflect the theory above. In other words, we assumed that each neuron can have *P* subfields, such that a neuron’s tuning is given by *f* (***s***; Ω_*n*_) + *m*_*n*_, where *f* (·) is given by Eq. (5), where 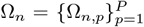 is the set of subfield parameters, and where we added a small constant firing baseline *m*_*n*_ to capture rare spurious spikes. Here, we permitted the tuning covariance matrix to rotate, expressing it as 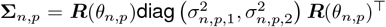. As such, the full set of parameters we wish to infer for each neuron is Ω_*n,p*_ = [*z*_*n,p*_, *a*_*n,p*_, ***µ***_*n,p*_, *σ*_*n,p*,1_, *σ*_*n,p*,2_, *θ*_*n,p*_, *m*_*n*_]. Subfield parameters of a given neuron were assumed to be independent. Lastly, we assumed each neuron in each small time bin *δt* to emit *r*_*i*_ spikes with Poisson probability *p* (*r*_*i*_ | Ω_*n*_, ***s***_*i*_) = 𝒫 (*r*_*i*_ | *f* (***s***_*i*_; Ω_*n*_)*δt*).

In lieu of prior knowledge about the distribution of subfield parameters, we opted to use uninformative priors for all parameters. As such, we assumed

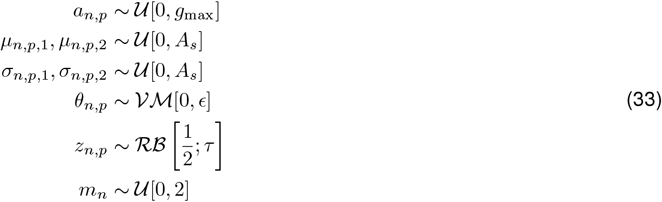

where is the relaxed Bernoulli distribution (a differentiable version of the Bernoulli distribution defined over [0, 1] with a temperature parameter (*τ*, which we set to 10^−4^) that controls how sharp the PDF is over [0, 1] [38]), *g*_max_ is the maximum possible gain of a subfield, *ϵ* is the precision parameter of the von-Mises distribution (which we typically set close to 0 to approximate a uniform distribution over the circle), and *A*_*s*_ is the arena size.

#### M3.4 Estimating the tuning parameter posterior

We use Markov Chain Monte Carlo (MCMC) sampling to separately estimate the tuning parameter posterior distribution for each neuron. Specifically, we used a no U-turn HMC sampler provided in NumPyro 0.12.1. For each neuron, we ran 4 sampling chains with 1,000 samples, after a burn-in phase of 1,000 samples. As the generative model’s firing rate function sums over individual subfields that can be active or inactive, its structure mirrors that of a finite approximation to an infinite mixture model. The associated parameter posterior turned out to be challenging for the sampler to navigate. Owing to the label switching / invariance inherent in fitting a mixture model, we also found that it was challenging to robustly quantify when a run of standard MCMC had converged. To circumvent these issues, we used neural transport ([39]) (implemented in NumPyro 0.12.1) upstream of MCMC. Briefly, neural transport uses a neural network to learn a differentiable mapping from an easily navigable sampling space (in this case, an isotropic Gaussian) to the goal posterior. Because the mapping must be differentiable, the neural transport approach implicitly collapses to a single invariant mode of the posterior. This solved the label switching phenomenon, and allowed us to better assess when MCMC had converged.

##### Measuring the number of subfields per-neuron

We approximated a Beta-Bernoulli mixture process with a finite Beta-Bernoulli mixture. We assumed that each neuron can have a maximum of 16 subfields. A subfield *p* of neuron *n* is considered “present” if the its indicator variable *z*_*n,p*_ exhibits an inferred posterior mean greater than 0.5. This is a convention that we chose to accommodate the use of a differentiable version of a Bernoulli random variable, which has support over all of [0, 1]. For each neuron, we counted the number of subfields as the number of indicator variables with inferred posterior mean greater than 0.5. Most posterior means of *z*_*n,p*_ had values either very close to 0 or 1, far away from the threshold.

##### Measuring convergence of MCMC

To measure convergence of MCMC, we first restricted our analysis to parameters of subfields which have a non-zero inferred indicator parameter *z*_*n,p*_. Over the 4 independent MCMC chains, we then compute the Split Gelman-Rubin (SGR) statistic over the samples, which measures the similarity of within-chain sample variance and between-chain sample variance. An SGR close to 1 indicates that the between-chain and within-chain variances are highly similar, suggesting that each chain is mixing well and properly sampling the posterior. We assessed that a sampling run on a neuron has converged if the SGR was less than 1.05.

##### Re-running when convergence fails

If convergence failed on a given neuron, we took two re-initialization approaches, depending on the source of the convergence failure. On some runs, the neural transport failed. In this case, we simply re-started the entire procedure, typically starting with a new random number seed. Alternatively, the neural transport succeeded, but the SGR was greater than 1.05. In this case, we found the parameter estimates associated with the sample from the previous run with the greatest log-likelihood, and re-initialized 4 new chains starting from this sample. This adaptive procedure allowed us to re-sample the posterior where all chains started from nearby locations in the posterior space.

##### Constraining the prior distribution support over gains to mitigate invariances

The use of the relaxed Bernoulli prior over the indicator variable, while useful for things like neural transport, introduced some invariances into the model. Namely, because *z*_*n,p*_ ∈ [0, 1], and because it is multiplied by each subfield’s gain *a*_*n,p*_, these quantities are inherently unidentifiable. This leads to difficulties with interpreting the posterior indicator variable samples. A convenient workaround for this issue was to set *g*_max_ independently for each neuron to the maximum value obtained in that neuron’s STA rate-map. We found that doing so reliably pushed the inferred value of the indicator variables for each subfield towards 0 and 1.

##### Visualizing marginal posteriors

To visualize marginal posteriors for inferred subfield parameters (such as in Fig. 4 in the main text), we applied kernel density estimation (KDE) with a Gaussian kernel whose bandwidth was chosen by Scott’s Rule ([40]) to the samples for a given parameter obtained using MCMC.

#### M3.5 Estimating 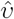 and 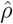

As a reminder, 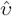 is the empirical shape parameter of the Gamma distribution of receptive field widths, and 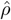 is the empirical correlation coefficient between the encoding dimension widths along each spatial dimension.

For obtaining both quantities, we started by estimating the maximum a-posteriori (MAP) widths for each subfield of each neuron. To do so, we took the posterior samples over the field widths *σ*_*n,p*,1_ and *σ*_*n,p*,2_ obtained using MCMC and used KDE with a Gaussian kernel whose bandwidth was chosen by Scott’s Rule ([40]) to approximate a continuous posterior density over all possible field widths in [0, *A*_*s*_]. We then approximated the MAP per-dimension field widths as 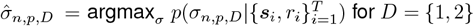 for *D* = {1, 2}.

##### Estimating 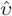

To estimate the empirical field size heterogeneity in a population, we took each estimated MAP width of each neuron, each subfield, and for both spatial dimensions, and obtained a distribution over the MAP widths by computing a weighted histogram of the MAP widths, where each MAP width was weighted by the inverse of its posterior variance, 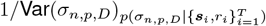. Finally, we densely sampled this new distribution, and computed the maximum likelihood (MLE) Gamma distribution shape parameter over these samples. To get a distribution over this parameter, we bootstrapped the MLE over samples from the weighted MAP width distribution.

##### Estimating 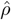

To obtain the empirical shape heterogeneity in the population, we computed the weighted correlation coefficient between the inferred MAP widths across all neurons and subfields. The weighted correlation coefficient is

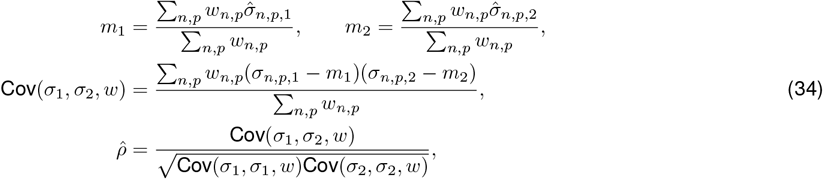

where

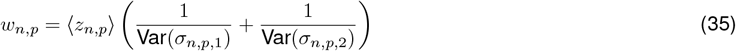

where the averages and variances in Eq. (35) are taken with respect to the approximate posterior distribution over the respective field width estimated by MCMC. To get a distribution over correlation coefficients, we bootstraped the weighted correlation over MAP widths.

##### Recovering 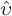 and 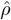 in synthetic datasets

We started by generating a synthetic population of neurons. In order to do this, we first ran the inference method on empirical data from a single run of the data from Pfeiffer and Foster [24]. From this, we obtained approximate empirical distributions of the model parameters, such as, the distribution of tuning curve centers *p*(***µ***_*n,p*_), the distribution of subfield gains *p*(*a*_*n,p*_), the approximate number of subfields per-neuron (this helps us set *ζ*, or the average field density, given that *P* = 16 in our finite mixture approximation), the distribution of baseline activity rates *p*(*m*_*n*_), and the distribution of tuning widths *p*(**Σ**_*n,p*_) (Extended Data Fig. 4).

Next, we generated a series of populations of 25-75 neurons, where either only the shape parameter, *υ*, of the Gamma distribution of synthetic field widths changed, or where only the principal synthetic field width correlation, *ρ*, changed between populations. All other synthetic population parameters were sampled according to the empirical parameter distributions obtained by running the inference method on real data. We then took real trajectories of rats from the Pfeiffer and Foster dataset, and computed the firing rate at each time point, *t*, according to *f* (***s***_*t*_; Ω_*n*_) + *m*_*n*_ where *f* (·) is given by Eq. (5), where Ω_*n*_ captures the sampled subfield parameters of the synthetic population, and where ***s***_*t*_ is the position of the animal at time *t*. We then generated spiking sequences according to the Poisson noise model described above and the computed firing rate.

Next, we applied our inference method to the spiking and position data. Finally, we applied the procedures above for estimating empirical 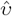 and 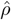 on the simulated spiking sequences.

We compared the reconstruction performance of our method to that obtained through using STA-based measurement tools. To obtain the STA-based estimates, we first computed the STA of each synthetic neuron in the population with 4×4 cm spatial bins. We then applied image segmentation algorithms in `skimage`, following methodology in [9]. We first apply a firing rate threshold that is either uniform across neurons (only consider bins in the STA that exceed 2 Hz for all neurons) or that is per-neuron (only consider bins in the STA that exceed a neuron’s average firing rate plus 1 standard deviation of the firing rate across spatial bins). Next, to impose a size threshold, we used the `skimage.morphology.remove_small_objects` function, with a threshold hyperparameter that dictates how large a contiguous region must be in order to be considered a region of interest in the STA (set to 8 cm^2^ [two bins] to impose a size cutoff, or to 0 cm^2^ to not impose a size cutoff). Finally, we used the `skimage.measure.regionprops` function, which fits ellipsoids around each contiguous region in the rate map that passes the thresholds outlined above. These region properties contained the estimated principal widths of each contiguous region of activity in the STA. We then pooled all principal widths across all neurons and subfields. We finally measured the empirical 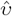 as the MLE Gamma shape over the pooled principal widths, and the 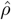 as the empirical correlation in the principal widths along each dimension of the pooled widths.

#### M3.6 Measuring eccentricity as a function of distance to nearest boundary

To test if shape heterogeneity is primarily driven by field distortions near boundaries, we measured the weighted correlation between distance of a subfield to its nearest boundary and a subfield’s eccentricity (lower eccentricity = shape more circular). For each subfield, we first compute the eccentricity, which is given by

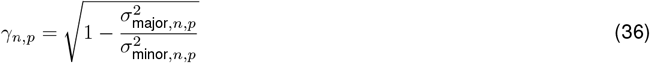

where *σ*_major,*n,p*_ is the MAP major subfield width (the larger of the two) of neuron *n*, subfield *p*, and *σ*_minor,*n,p*_ is the MAP minor subfield width (the smaller of the two). To compute the weighted correlation between *γ*_*n,p*_ and the subfield’s distance to its nearest boundary, we compute the weights as before in Eq. (35) and compute the correlation with Eq. (34).

### M4 Measuring the explanatory power of the STA and our inference method

#### M4.1 Fraction of deviance explained

We measured goodness of fit with the fraction of deviance explained by a given model (either the STA or our inference method) in comparison to to a null model for each neuron: 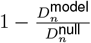 where,

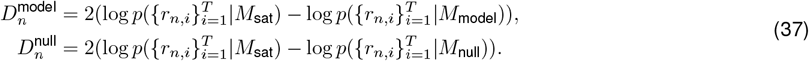

The saturated and null models both assumed that spikes are generated according to a Poisson process based on the firing rate. The null model assumed that the firing rate is uniform across the arena and equal to the average firing rate of the neuron, whereas the saturated model assumed that the firing rate corresponds to the observed spike count in each time bin. *M*_STA_ is obtained by computing the rate map with 4×4cm bins. For the inference method model, we used the maximum log likelihood sample, 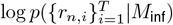, obtained when sampling the posterior through MCMC.

#### M4.2 Trajectory switch experiment

To demonstrate that the STA is biased by the trajectory taken by the animal, we devised a trajectory switch experiment. In this experiment, we generated a population of synthetic neurons, where, again, the parameters of each subfield of each neuron in the population followed the respective distributions obtained by our inference method on a single sample run of real data. As above, we took a real trajectory of a rat (trajectory A) from the Pfeiffer and Foster dataset, and computed the firing rate at each time point, *t*, according to *f* (***s***_*t*_; Ω_*n*_) + *m*_*n*_ where *f* (·) is given by Eq. (5), where Ω_*n*_ captures the sampled subfield parameters of the synthetic population, and where ***s***_*t*_ is the position of the animal at time *t*. We then generated spiking sequences according to the Poisson noise model described above and the computed firing rate. We then computed the STA from these generate spikes.

Next, we followed the same spike generation procedure above, but with a different trajectory (trajectory B). We then computed the FDE for spikes from each trajectory, but where the *M*_model_ for both sequences is the STA from trajectory A. We quantified the change in explanatory power as the percent change in the average deviance explained between the trajectories.

To show that our inference method is not hampered by such biases, we applied the same trajectory switch experiment, but where *M*_model_ for both sequences is the field parameters inferred by our method on spikes generated from trajectory A. As shown in the main text, we found only a minor change in the FDE as a result of the trajectory switch for our method.

## Extended Data

**Extended Data Figure 1:**
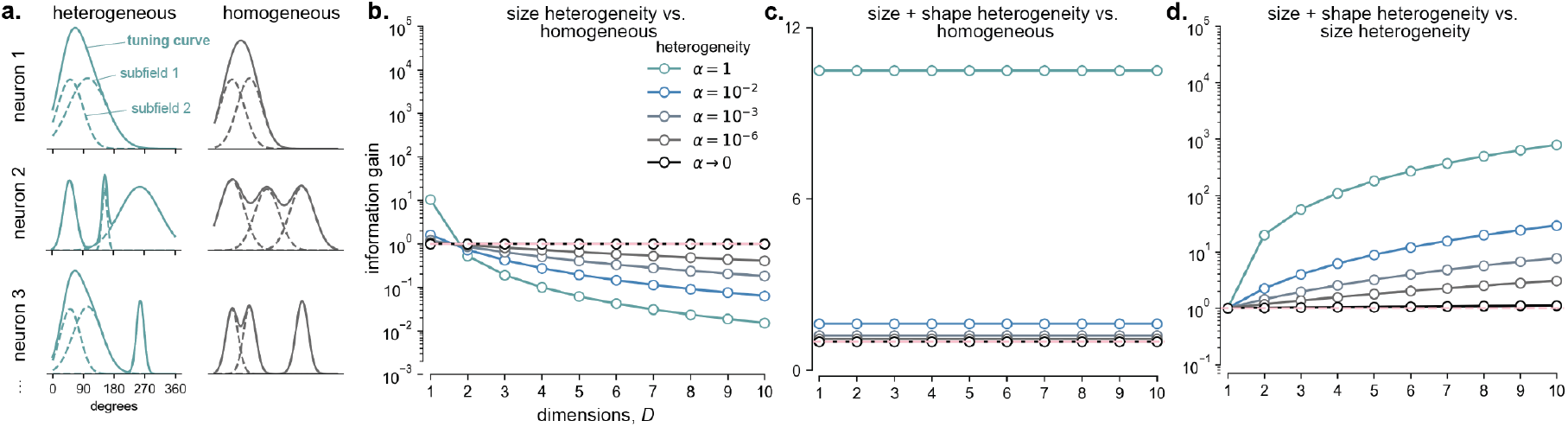
Information gain for multimodal tuning curves is dimensionality-dependent. **a**. Schematic of multimodal tuning curves (*D* = 1 shown here only) with receptive field heterogeneity (left) and without (right). Note that subfield gains can vary even if their receptive fields are homogeneous. **b**. Information gain with just size heterogeneity alone in the multimodal case (corresponds to unimodal case, **Fig. 1d** main text). **c**. Information gain with size and shape heterogeneity in the multimodal case (corresponds to unimodal case, **Fig. 2d** main text). **d**. Information gain with size and shape heterogeneity versus just size heterogeneity (corresponds to unimodal case, **Fig. 2c** main text). Even though the information gain for multimodal tuning curves exhibits a different dimensionality-dependence than for unimodal ones, it nonetheless predicts that for encoding two-dimensional stimuli, size and shape heterogeneity are beneficial. Indeed, unlike for unimodal tuning curves, size heterogeneity alone in the *D* = 2 case is actually harmful.

**Extended Data Figure 2:**
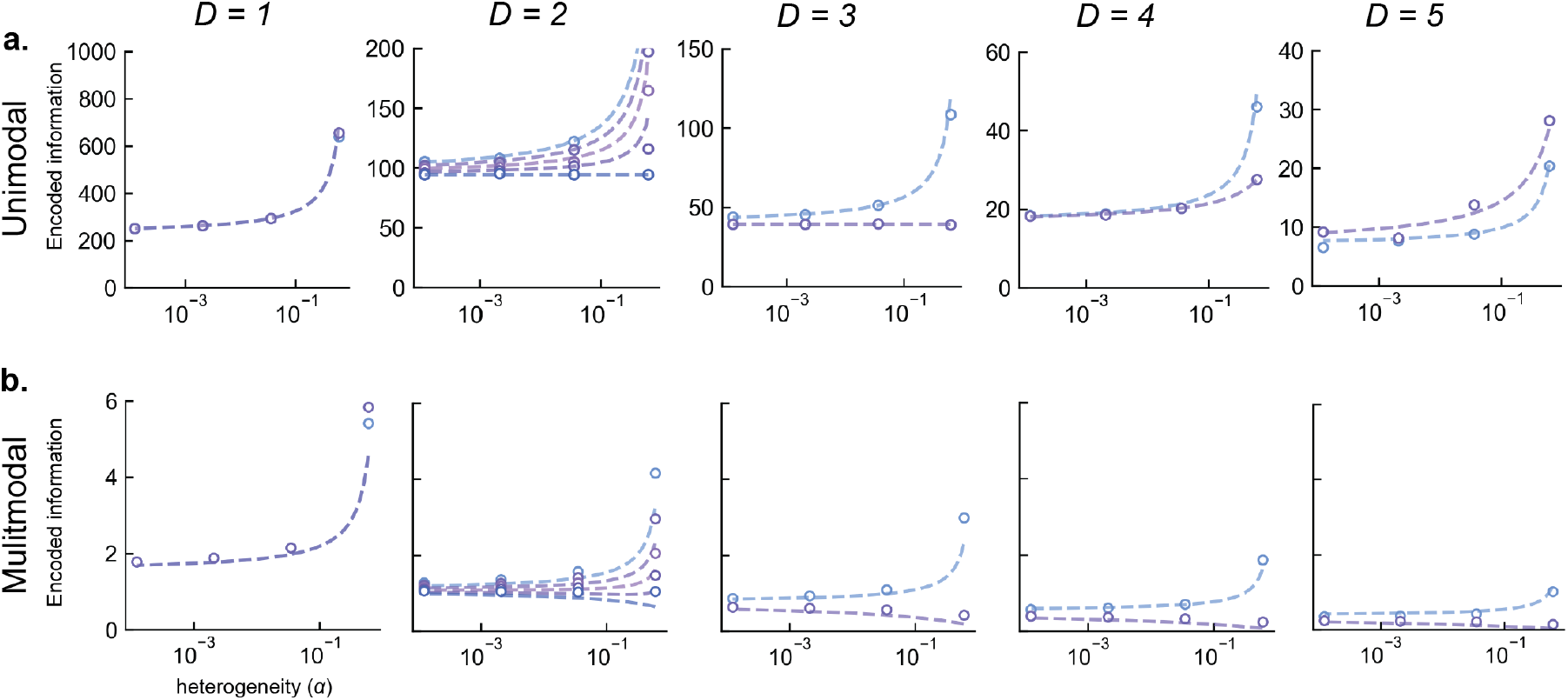
Verification of analytical expressions for encoded information via simulated populations. We verified our analytic theoretical predictions by empirically computing the encoded information in simulated populations with *N* = 800 neurons that exhibit unimodal (row **a**.) and multimodal (row **b**.) tuning curves across various stimulus dimensionalities (*D* ∈ {1, 2, 3, 4, 5}; dashed lines = theory, circles = simulations for selected *α*, see Methods M2 for details). For *D* ∈ {1, 3, 4, 5} we show the encoded information across *α* for populations with shape heterogeneity (light blue) and without shape heterogeneity (purple). For *D* = 2 we show the encoded information for different degrees of shape heterogeneity (colors).

**Extended Data Figure 3:**
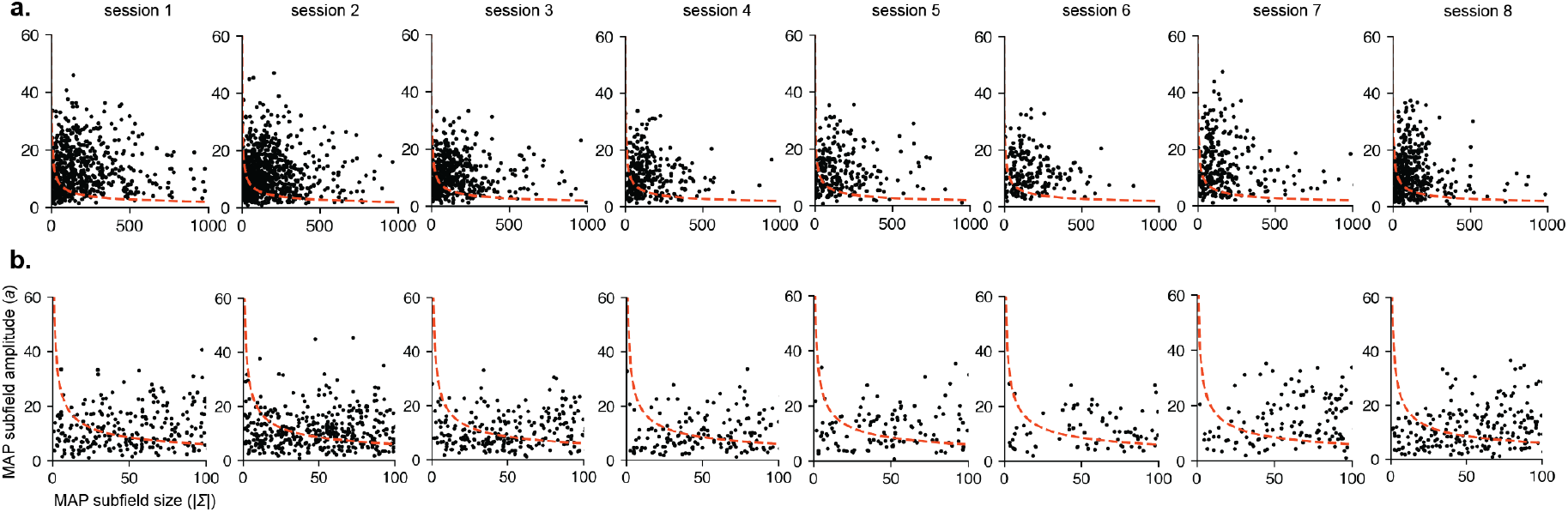
The tuning amplitude of rodent place cells does not appear to be determined by metabolic constraints. If the amplitude is sensitive to metabolic costs, we would expect subfields with larger receptive fields to have a smaller gain. **a**. Maximum a-posteriori (MAP) subfield sizes (here measured by the determinant of the tuning width covariance matrix) versus MAP subfield amplitude (dots = subfields) across all sessions. Dashed red line corresponds to theoretical amplitudes predicted by our theory of metabolic constraints (up to an arbitrary per session-scaling factor). **b**. Plots showing the same relationship for subfield sizes up to 100 to better illustrate the deviation from the theory for smaller subfields.

**Extended Data Figure 4:**
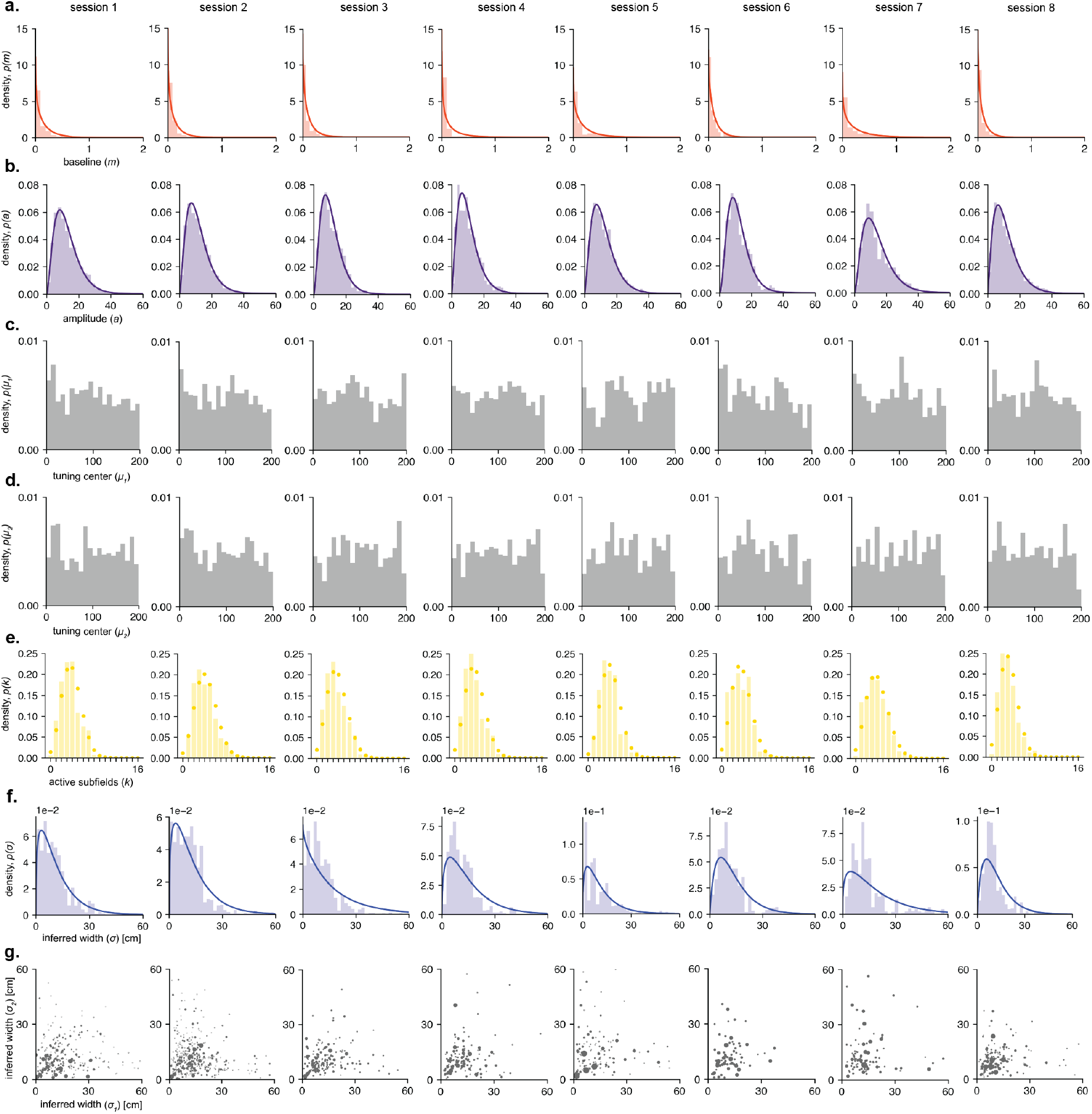
Tuning statistics of rodent place cells across sessions. **a**. Inferred baseline activity posterior across all neurons, red lines corresponds to maximum likelihood Gamma distribution fit. **b**. Inferred amplitude posterior across all subfields, purple lines corresponds to maximum likelihood Gamma distribution fit. We found that the amplitude distribution was best fit by a Gamma distribution. This differs from the assumption we made in our theory, where we assumed subfield amplitudes to be drawn from a uniform distribution. However, this has no consequence for the validity of our theory when applied to these data, as subfield amplitude contributes to information gain only through its mean, irrespective of the shape of the distribution itself. **c**. Inferred posterior distribution of subfield tuning curve centers along one spatial dimension, which are roughly uniformly distributed across the arena. **d**. Same as **c**. but for the alternate spatial dimension. **e**. Inferred posterior distribution over number of subfields per neuron. Yellow dot corresponds to beta-Binomial maximum likelihood fit. **f**. Distributions over inferred maximum a-posteriori (MAP) widths for each session, blue line the maximum likelihood fit Gamma distribution. **g**. Inferred MAP principal widths across each dimension (dots = subfields, size of dot is proportional to sum of the inverse of the marginal posterior variances of each principal width).

**Extended Data Figure 5:**
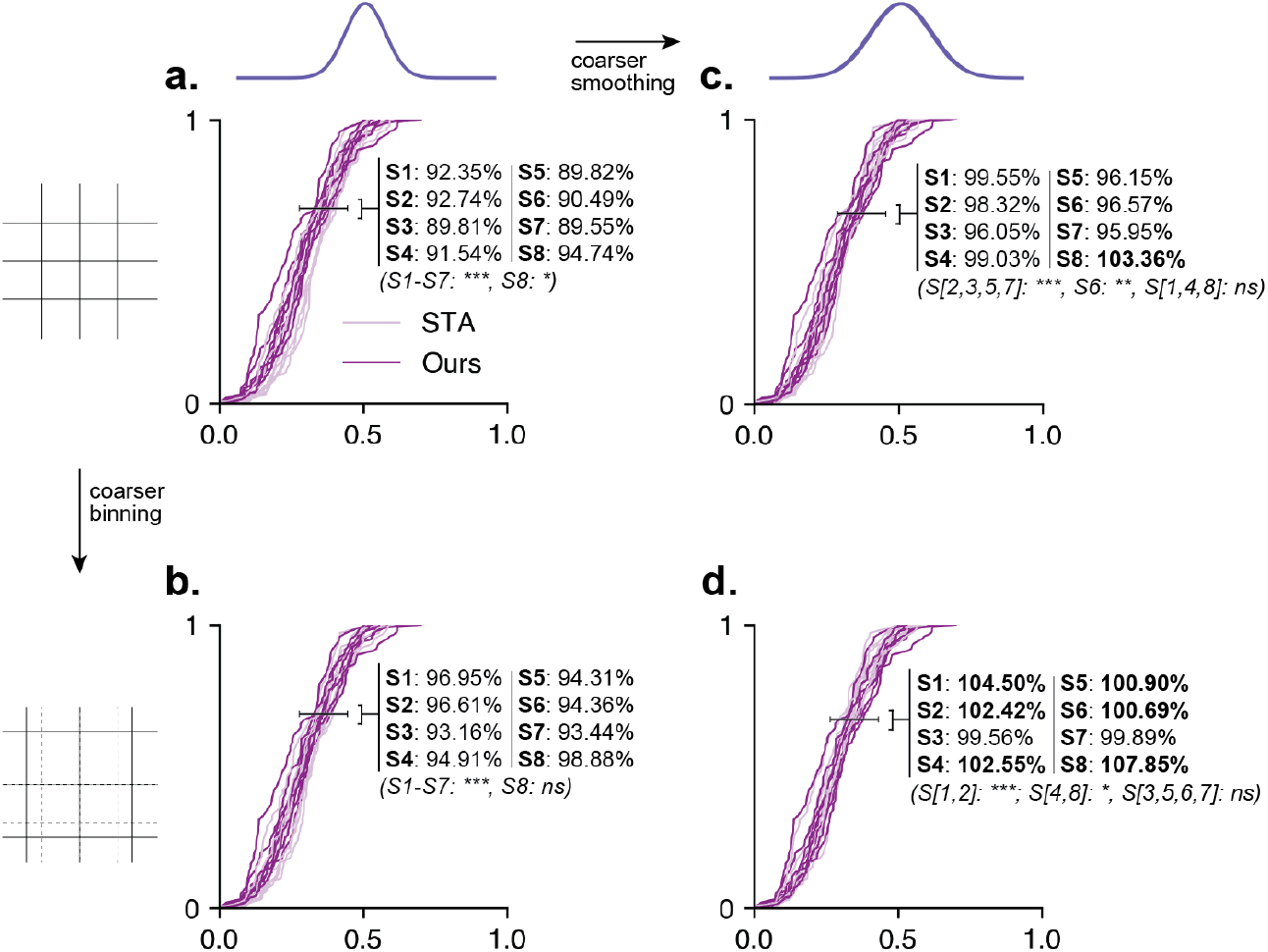
Fraction of deviance explained by the STA is sensitive to modest changes in hyperparameters. **a**. The baseline FDE reported in the main text corresponds to binning the 2×2m arena into 4×4cm bins and applying Gaussian smoothing with *σ* = 4cm. **b**. FDE CDFs with slightly coarser binning (5×5cm) but the same smoothing kernel. **c**. FDE CDFS with 4×4cm binning and applying Gaussian smoothing with *σ* = 5.2cm. **d**. FDE CDFs with 5×5cm binning and applying Gaussian smoothing with *σ* = 5.2cm. For all panels, session deviance of STA explained by our method is bolded if it exceeds 100% (in other words, if our method better explains the data). Significance indicated by *p*-values on paired t-test in **Table 1**.

**Table 1:** P-values for paired t-tests run for each session under different binning and smoothing hyperparameters for the STA, as shown in **Extended Data Fig. 5**.

|  | Panel **b.** | Panel **c.** | Panel **d.** |
| --- | --- | --- | --- |
| S1 | $p = 1.347 \times 10^{-5}$ | $p = 0.609$ | $p = 4.78 \times 10^{-6}$ |
| S2 | $p = 7.449 \times 10^{-19}$ | $p = 6.329 \times 10^{-5}$ | $p = 1.74 \times 10^{-7}$ |
| S3 | $p = 2.051 \times 10^{-19}$ | $p = 7.374 \times 10^{-5}$ | $p = 0.603$ |
| S4 | $p = 7.304 \times 10^{-7}$ | $p = 0.461$ | $p = 0.034$ |
| S5 | $p = 5.956 \times 10^{-11}$ | $p = 1.875 \times 10^{-5}$ | $p = 0.344$ |
| S6 | $p = 5.513 \times 10^{-9}$ | $p = 0.002$ | $p = 0.522$ |
| S7 | $p = 9.509 \times 10^{-13}$ | $p = 3 \times 10^{-4}$ | $p = 0.906$ |
| S8 | $p = 0.652$ | $p = 0.340$ | $p = 0.022$ |

**Extended Data Figure 6:**
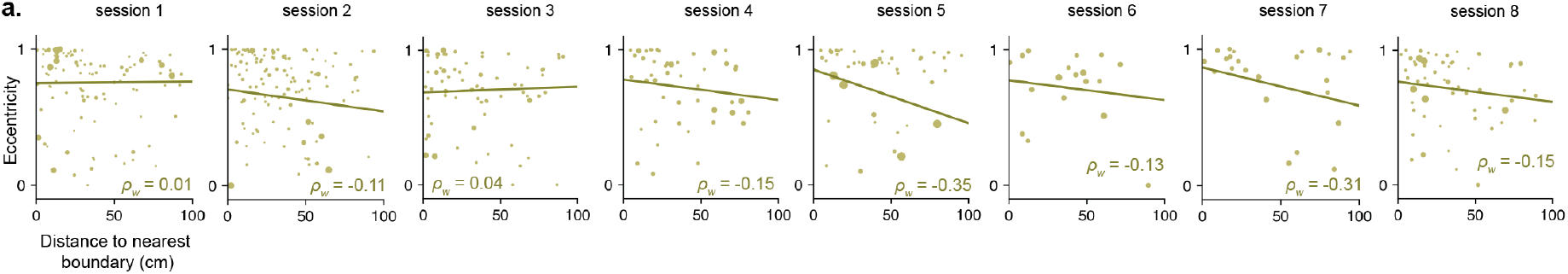
Shape heterogeneity is persistent in subfields across the arena. **a**. Distance to nearest boundary plotted against field eccentricity (lower eccentricity = subfield shape more circular; *ρ*_*w*_ = weighted correlations for each session [overlaid on plot], dots = subfields, dark line = weighted least squares regression where weights are the weights used to compute *ρ*_*w*_). Bootstrapped *p*(*ρ*_*w*_ *>* 0) for each session (*p*(*ρ*_*w*_ *>* 0) *<* 0.05 threshold for significance): S1: 0.49 (ns), S2: 0.27 (ns), S3: 0.58 (ns), S4: 0.12 (ns), S5: 0.06 (ns), S6: 0.30 (ns), S7: 0.019 (*), S8: 0.11.

**Extended Data Figure 7:**
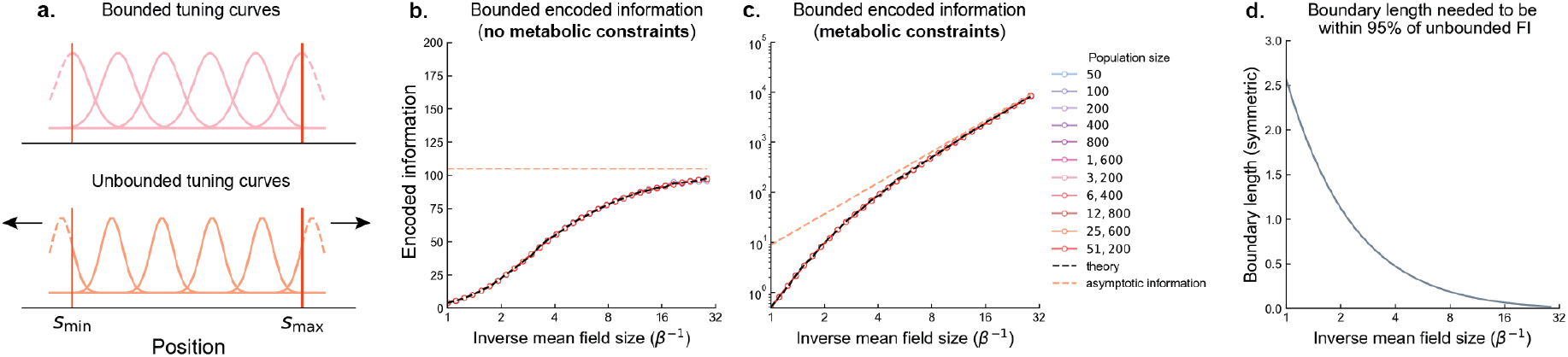
The influence of bounding tuning curve centers on encoded information. **a**. A schematic illustrating the difference between bounded tuning curves (top panel) and unbounded tuning curves (bottom panel). Unbounded tuning curve centers can span an arbitrarily large domain beyond the stimulus range, whereas bounded tuning curves can only exceed the stimulus range by finite amounts (and in the plots here–as in [8]–are rigidly bound to match the stimulus domain). **b**. Bounded encoded information (both theoretical predictions, and simulated population of different sizes *N*, see Methods M2) in the metabolically unconstrained case, which is systematically lower than the asymptotic encoded information. **c**. The same as **b**. but for metabolically constrained populations. **d**. A quantification of the influence of boundary effects as a function of average receptive field size. This shows how far outside the stimulus boundaries tuning curves centers must be placed in order to achieve 95% of the asymptotic encoded information in the metabolically unconstrained case. In the limit of infinitely small tuning curves, we recover full asymptotic encoded information, as suggested by panels **b**,**c**.

**Extended Data Figure 8:**
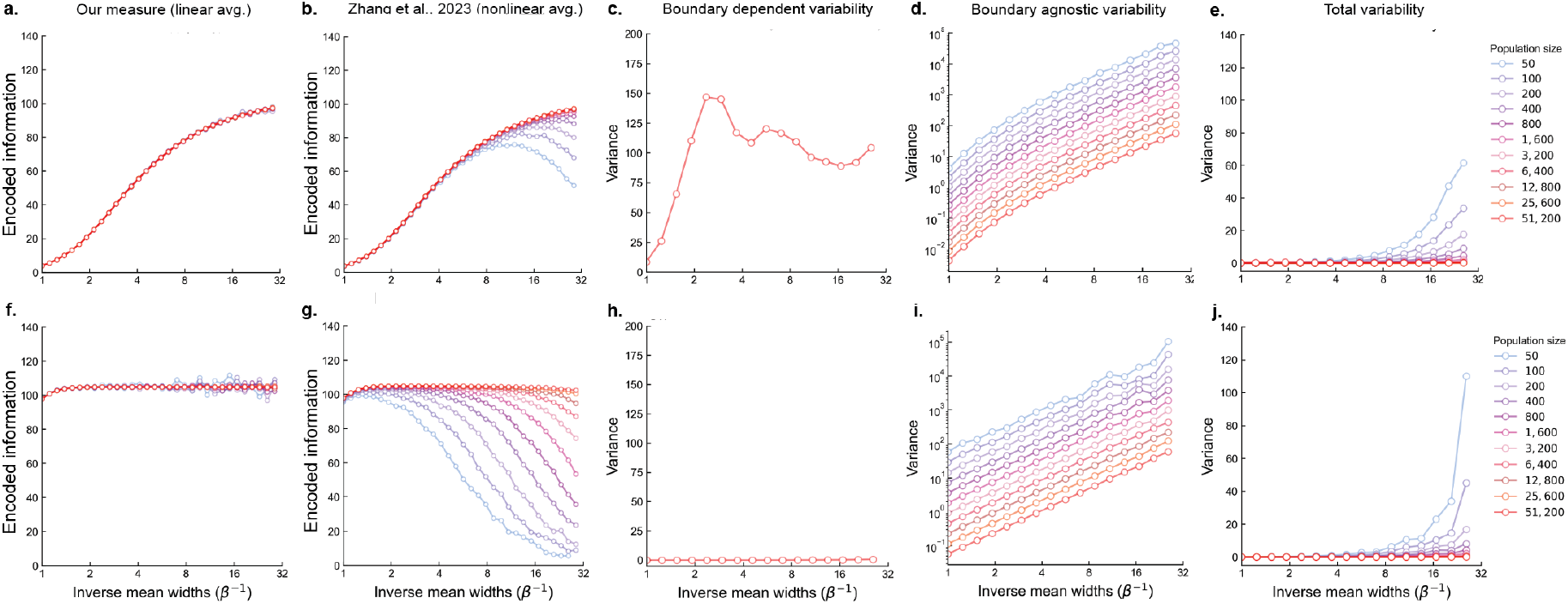
Asymptotic and bounded information variability is quenched in large populations. Panels in the top row show results when tuning curve centers are bounded to lie within the stimulus range, whereas the bottom row shows results when these centers can extend significantly beyond that range. **a**. Our measure of encoded information, using linear averaging of the population FI. **b**. Zhang et al.’s measure of encoded information, which we have shown (SI 6) equals ours minus components that depend on the variability of per-neuron encoded information, inversely scaled by the size of the population. Here we measure the encoded information in populations with heterogeneity *α* = 8 *×* 10^−2^ (*υ* = 3.5). At such a setting, the difference in our measures vanishes for populations as small is *N* = 800 neurons. **c**. If tuning curve centers are bounded, encoded information varies across the stimulus range. The resulting variance, which is shown here, additionally reduces Zhang et al.’s information measure. This source of variability is independent of population size, but is here estimated with a population of size *N* = 51, 200. **d**. Variation in per-neuron FI that is agnostic to whether tuning curve centers are bounded. **e**. The total variability (the sum of variabilities depicted in panels **c**. and **d**. divided by the linear encoded information depicted in **a**.). **f-j**. The same as **a-e**., but for the case of unbounded tuning curve centers. Note that in the unbounded case, boundary-dependent variability (**h**.) does not contribute to overall variability, as predicted by the theory. In both cases, as *N*→ *∞*, overall variability is driven towards 0 (**e**., **j**.), as predicted by the theory. This quenching in variability in larger populations drives the measure used in [8] to become indistinguishable from ours in larger populations.

## Notes

### Competing Interest Statement

The authors have declared no competing interest.

