## Supplemental Information for "A unifying theory of receptive field heterogeneity predicts hippocampal spatial tuning"

### 1 Outline

This supplement has the following structure. In Sec. 2, we first explain how we measure encoded information. In this section, we also discuss how we parameterize our model population. In Secs. 3-5, we derive analytical expressions for encoded information under different assumptions and relate them to each other. In Sec. 6.1, we discuss how our work applied to two-dimensional stimuli relates to that of Zhang et al. [1].

### 2 Measuring encoded information and model assumptions

#### 2.1 How we quantify information, and properties of this measure

We aim to quantify how encoded information in neural activity depends on shape and width heterogeneity of neural tuning curves in a population. To do so, we assume a population of $N$ Poisson spiking neurons emitting a population vector of spikes $\boldsymbol{r} \in \mathbb{N}_0^N$ in response to a stimulus $\boldsymbol{s} \in \mathbb{R}^D$ (where $D$ is the stimulus dimensionality; for $D = 2$, $\boldsymbol{s}$ can be interpreted as the spatial position of an animal), and where $s_i \sim \mathcal{U}[s_{\text{min}}, s_{\text{max}}]$. We assess encoded information with the Fisher information (FI) that the population activity $\boldsymbol{r}$ provides about $\boldsymbol{s}$. This yields a matrix $N\boldsymbol{\mathcal{I}}(\boldsymbol{s}) \in \mathbb{R}^{D\times D}$, where $\boldsymbol{\mathcal{I}}(\boldsymbol{s}) \in \mathbb{R}^{D\times D}$ denotes the average FI matrix per neuron (hence the $N$ pre-factor). We average this FI matrix across stimuli, $\bar{\boldsymbol{\mathcal{I}}} = \langle N\boldsymbol{\mathcal{I}}(\boldsymbol{s})\rangle_{p(\boldsymbol{s})}$, and summarize it by its determinant, in line with how the determinant of a covariance matrix measures the generalized variance. Furthermore, to make our information measure comparable across choices of $N$ and $D$, we—in line with previous literature [1]—re-scale it by

$$\hat{\mathcal{I}} = \frac{1}{N}\sqrt[D]{|\bar{\boldsymbol{\mathcal{I}}}|} = \sqrt[D]{\left|\frac{\bar{\boldsymbol{\mathcal{I}}}}{N}\right|}. \tag{1}$$

This choice of measuring encoded information captures both the local discriminability of the population code and also provides a lower bound on the decoding error of downstream populations, which we show in the following subsection. When computing encoded information, we will survey the FI in populations that exhibit shape and width heterogeneity under two main conditions: one in which tuning curve centers are unbounded (can exceed the stimulus space by an arbitrarily large extent) and one in which the tuning curve centers of the population are bounded by a finite extent beyond the stimulus space (and, as a special case, exactly bound to the stimulus space). We call the former case the "asymptotic encoded information," as we find it provides an upper bound on the bounded encoded information (more details provided in Sec 5).

##### 2.1.1 The encoded information lower bounds the stimulus decoding error

Let $\boldsymbol{\epsilon}(\boldsymbol{s})$ denote the $D$-dimensional vector that contains the stimulus decoding error across all stimulus dimensions, and let $\text{Cov}(\boldsymbol{\epsilon}(\boldsymbol{s}))$ be its covariance matrix. There are two common ways to summarize this covariance matrix as a scalar decoding error. The first is the normalized trace, $\text{Tr}\left(\text{Cov}(\boldsymbol{\epsilon}(\boldsymbol{s}))\right)/D$, which equals the mean-squared decoding error across stimulus dimensions. The second is the $D$th root of its determinant, $\sqrt[D]{|\text{Cov}(\boldsymbol{\epsilon}(\boldsymbol{s}))|}$, which we term the *generalized decoding error* (on account of the fact that it is inspired by the generalized variance of an estimator, or the determinant of the estimator covariance), that captures error relations across dimensions that might not be captured by the trace. For isotropic covariance matrices, these two measures become equivalent. In what follows we will show that our encoded information measure is a lower bound on both of these decoding errors.

Central to our approach is the Cramer-Rao bound that relates the covariance of unbiased estimators to the Fisher information matrix. In particular, for some given population parameters $\boldsymbol{\Omega}$ it states that

$$\text{Cov}(\boldsymbol{\epsilon}(\boldsymbol{s};\boldsymbol{\Omega})) \succeq \boldsymbol{\mathcal{I}}(\boldsymbol{s};\boldsymbol{\Omega})^{-1}, \tag{2}$$

where we have assumed that $\boldsymbol{\epsilon}(\boldsymbol{s};\boldsymbol{\Omega})$ is the decoding error of an unbiased estimator, that is $\langle\boldsymbol{\epsilon}(\boldsymbol{s};\boldsymbol{\Omega})\rangle_{p(\boldsymbol{r}|\boldsymbol{s};\boldsymbol{\Omega})} = \boldsymbol{0}$, and where $\boldsymbol{A} \succeq \boldsymbol{B}$ denotes the Loewner order that implies that $\boldsymbol{A} - \boldsymbol{B} \in \texttt{SPSD}$, where SPSD is the set of symmetric positive semi-definite matrices.

In Appendix A.3, we show that the generalized decoding error lower bounds the mean-squared discrimination error, that is, $\text{Tr}\left(\text{Cov}(\boldsymbol{\epsilon}(\boldsymbol{s}))\right)/D \geq \sqrt[D]{|\text{Cov}(\boldsymbol{\epsilon}(\boldsymbol{s}))|}$. As such, a measure that lower bounds the generalized decoding error also lower bounds the mean-squared decoding error.

To show that the $D$-th root of this generalized variance is bounded by our encoded information measure, let us assume that the FI matrix $\boldsymbol{\mathcal{I}}(\boldsymbol{s};\boldsymbol{\Omega})$ is symmetric positive definite (SPD). This assumption would only be violated if the FI along some stimulus direction becomes zero, which is only the case in degenerate scenarios (e.g., at some extremes of the parameter ranges). Then, as $\boldsymbol{A} \succ \boldsymbol{B}$ implies that $|\boldsymbol{A}| > |\boldsymbol{B}|$, the Cramer-Rao bound, together with $\sqrt[D]{\cdot}$ being an increasing function implies that

$$\left\langle \sqrt[D]{|\text{Cov}(\boldsymbol{\epsilon}(\boldsymbol{s};\boldsymbol{\Omega})|}\right\rangle_{p(\boldsymbol{s},\boldsymbol{\Omega})} \geq \left\langle \sqrt[D]{|\boldsymbol{\mathcal{I}}(\boldsymbol{s};\boldsymbol{\Omega})^{-1}|}\right\rangle_{p(\boldsymbol{s},\boldsymbol{\Omega})} = \left\langle \frac{1}{\sqrt[D]{|\boldsymbol{\mathcal{I}}(\boldsymbol{s};\boldsymbol{\Omega})|}}\right\rangle_{p(\boldsymbol{s},\boldsymbol{\Omega})} \tag{3}$$

where the final equality follows from the fact that $|\boldsymbol{A}^{-1}|= 1/|\boldsymbol{A}|$. Furthermore, as we show in Appendix A.4, the mapping $\boldsymbol{X} \mapsto |\boldsymbol{X}|^{-1/D}$ is convex over the set of symmetric positive definite (SPD) matrices. Therefore, by Jensen's inequality we have $\left\langle 1/\sqrt[D]{|\boldsymbol{\mathcal{I}}(\boldsymbol{s};\boldsymbol{\Omega})|}\right\rangle_{p(\boldsymbol{s},\boldsymbol{\Omega})} \geq 1/\sqrt[D]{|\langle \boldsymbol{\mathcal{I}}(\boldsymbol{s};\boldsymbol{\Omega})\rangle_{p(\boldsymbol{s},\boldsymbol{\Omega})}|}$, leading to the final bound,

$$\left\langle \sqrt[D]{|\mathsf{Cov}(\boldsymbol{\epsilon}(\boldsymbol{s};\boldsymbol{\Omega})|}\right\rangle_{p(\boldsymbol{s},\boldsymbol{\Omega})} \geq \frac{1}{\sqrt[D]{|\langle \boldsymbol{\mathcal{I}}(\boldsymbol{s};\boldsymbol{\Omega})\rangle_{p(\boldsymbol{\Omega})}|}} = \frac{1}{N\hat{\mathcal{I}}}. \tag{4}$$

Therefore, the inverse of our encoded information measure, scaled by population size $N$, is a lower bound on the $D$th root of the generalized decoding error, and thus also a lower bound on the mean-squared decoding error.

Note that Zhang & Sejnowski in [2] also used an argument based on the Cramer-Rao bound to motivate their encoded information measure. In fact, it can be shown that, as long as $\langle \boldsymbol{\mathcal{I}}(\boldsymbol{s};\boldsymbol{\Omega})\rangle_{p(\boldsymbol{s},\boldsymbol{\Omega})} = \lambda \boldsymbol{I}$ holds, their measure equals ours. Thus, a similar bound applies to their work. In Sec. 6.1 we further explore how the encoded information measure we introduce relates to an alternative measures used in [1].

##### 2.1.2 A simple, locally linear decoder achieves the optimal decoding error

We measure encoded information through the Fisher information. As we have seen in the previous section, one rationale for this approach is that the encoded information consequently lower-bounds the decoding error of a minimum variance unbiased decoder. This bound is agnostic to the precise form of the decoder—instead it dictates only what error even the "best" unbiased decoder can achieve. While we will show below that heterogeneous tuning achieves higher encoded information (and thus, a lower achievable decoding error in principle), one might reasonably speculate that such decoding error could only be achieved by a highly biologically implausible decoder. Here, we show that the optimal decoder of the model population we consider is a simple, locally linear decoder. This is true regardless of the tuning properties of the underlying population. Indeed, we show that the tuning properties only influence the magnitude of the linear decoding weights. That the optimal decoding error can be achieved by such a simple decoder merits the focus of this work on the encoded information.

To show this, we generalize the results of [3] to arbitrary stimulus dimensionality $D$. The authors in [3] show that for any population whose emission probability ($p(\boldsymbol{r}|s)$) is in the exponential family with linear sufficient statistics, the *minimum variance unbiased locally linear decoder* (MVULLD) is "optimal" in the sense that it is efficient with respect to the Cramer-Rao bound. In other words, the variance of the estimator for the MVULLD is equal to the inverse of the Fisher information. The authors in [3] showed this for $s \in \mathbb{R}$. Here, we show it is also the case for $\boldsymbol{s} \in \mathbb{R}^D$.

As in [3], we start by assuming that the likelihood of observing population activity vector $\boldsymbol{r} \in \mathbb{R}^N$ in response to stimulus $\boldsymbol{s} \in \mathbb{R}^D$ takes the form of an exponential family with linear sufficient statistics,

$$p(\boldsymbol{r}|\boldsymbol{s}) = g(\boldsymbol{s})\Phi(\boldsymbol{r})\exp(\boldsymbol{h}(\boldsymbol{s})^T\boldsymbol{r}) \tag{5}$$

where $g(\boldsymbol{s}) = \frac{1}{\int \Phi(\boldsymbol{r})\exp(h(\boldsymbol{s})^T\boldsymbol{r})d\boldsymbol{r}}$, and $\Phi(\cdot)$ and $\boldsymbol{h}(\cdot)$ are known functions. It is easy to show that the population model that we describe further below (assuming sufficiently dense tuning) satisfies this assumption [4]. Then, as in [3], we will first write down the expression for the Fisher information for this model population. We will then show that the covariance of the MVULLD is equal to the inverse of the model Fisher information, thereby saturing the Cramer-Rao bound, and suggesting that the MVULLD is the optimal decoder for the model population.

To find the Fisher information matrix associated with the above $p(\boldsymbol{r}|\boldsymbol{s})$, note that $\boldsymbol{h}(\boldsymbol{s}) : \mathbb{R}^D \to \mathbb{R}^N$ such that $\nabla_s\left(\boldsymbol{h}(\boldsymbol{s})^\top\boldsymbol{r}\right) = \nabla\boldsymbol{h}(\boldsymbol{s})^\top\boldsymbol{r}$, where $\nabla_s\boldsymbol{h}(\boldsymbol{s})$ denotes the $N \times D$ Jacobian matrix of $\boldsymbol{h}(\cdot)$ evaluated at $\boldsymbol{s}$. Then, some algebra shows that $\nabla_s \log g(\boldsymbol{s}) = -\nabla_s\boldsymbol{h}(\boldsymbol{s})^\top\boldsymbol{f}(\boldsymbol{s})$, where $\boldsymbol{f}(\boldsymbol{s}) = \langle\boldsymbol{r}\rangle_{p(\boldsymbol{r}|\boldsymbol{s})}$ denotes the mean population activity in response to stimulus $\boldsymbol{s}$. Together, this allows us to compute the Fisher information matrix by

$$\begin{aligned}\boldsymbol{\mathcal{I}}(\boldsymbol{s}) &= \left\langle \left(\nabla_s \log p(\boldsymbol{r}|\boldsymbol{s})\right)\left(\nabla_s \log p(\boldsymbol{r}|\boldsymbol{s})\right)^\top\right\rangle_{p(\boldsymbol{r}|\boldsymbol{s})} \\ &= \left\langle \left(\nabla_s\boldsymbol{h}(\boldsymbol{s})^\top(\boldsymbol{r}-\boldsymbol{f}(\boldsymbol{s})\right)\left(\nabla_s\boldsymbol{h}(\boldsymbol{s})^\top(\boldsymbol{r}-\boldsymbol{f}(\boldsymbol{s}))\right)^\top\right\rangle_{p(\boldsymbol{r}|\boldsymbol{s})} \\ &= \nabla_s\boldsymbol{h}(\boldsymbol{s})^\top\left\langle(\boldsymbol{r}-\boldsymbol{f}(\boldsymbol{s}))(\boldsymbol{r}-\boldsymbol{f}(\boldsymbol{s}))^\top\right\rangle_{p(\boldsymbol{r}|\boldsymbol{s})}\nabla_s\boldsymbol{h}(\boldsymbol{s}) \\ &= \nabla_s\boldsymbol{h}(\boldsymbol{s})^\top\boldsymbol{\Sigma}(\boldsymbol{s})\nabla_s\boldsymbol{h}(\boldsymbol{s}),\end{aligned} \tag{6}$$

where $\boldsymbol{\Sigma}(\boldsymbol{s}) = \mathsf{cov}(\boldsymbol{r}|\boldsymbol{s})$ is the $N \times N$ population activity noise covariance. To express the FI matrix in terms of $\boldsymbol{f}(\boldsymbol{s})$, we furthermore generalize [4] to multidimensional $\boldsymbol{s}$ to show that $\nabla_s\boldsymbol{f}(\boldsymbol{s}) = \nabla_s\langle\boldsymbol{r}\rangle_{p(\boldsymbol{r}|\boldsymbol{s})} = \boldsymbol{\Sigma}(\boldsymbol{s})\nabla_s\boldsymbol{h}(\boldsymbol{s})$ (again requiring some algebra [3, 4]), where $\nabla_s\boldsymbol{f}(\boldsymbol{s})$ denotes the $N \times D$ Jacobian matrix of $\boldsymbol{f}(\cdot)$ evaluated at $\boldsymbol{s}$. Using in turn $\nabla_s\boldsymbol{h}(\boldsymbol{s}) = \boldsymbol{\Sigma}(\boldsymbol{s})^{-1}\nabla_s\boldsymbol{f}(\boldsymbol{s})$ allows us to re-write the Fisher information as

$$\boldsymbol{\mathcal{I}}(\boldsymbol{s}) = \nabla_s\boldsymbol{f}(\boldsymbol{s})^\top\boldsymbol{\Sigma}(\boldsymbol{s})^{-1}\nabla_s\boldsymbol{f}(\boldsymbol{s}). \tag{7}$$

What remains is to show that the MVULLD covariance saturates to the inverse of this expression. We follow the derivation from [3]. First, we assume that the decoded stimulus $\hat{s}$ is computed as a linear projection of population activity through a set of weights $\boldsymbol{W} \in \mathbb{R}^{N\times D}$. The unbiased linear estimator is given as

$$\hat{\boldsymbol{s}} - \boldsymbol{s}_0 = \boldsymbol{W}^\top \left(\boldsymbol{r} - \boldsymbol{f}(\boldsymbol{s}_0)\right). \tag{8}$$

Indeed the expectation on the RHS around $\boldsymbol{s}_0$ is $\boldsymbol{W}^\top \left(\langle \boldsymbol{r}\rangle_{p(\boldsymbol{r}|\boldsymbol{s}_0)} - \boldsymbol{f}(\boldsymbol{s}_0)\right) = \boldsymbol{0}$, confirming that this estimator is unbiased. Then, our aim is to find decoding weights $\boldsymbol{W}$ that are locally unbiased, such that

$$\nabla_s \langle \hat{\boldsymbol{s}}\rangle_{p(\boldsymbol{r}|\boldsymbol{s})} = \boldsymbol{I} \tag{9}$$

which yields constraints

$$\boldsymbol{W}^T \nabla_s \boldsymbol{f}(\boldsymbol{s}) = \boldsymbol{I}. \tag{10}$$

To obtain the MVULLD, we aim to minimize the local estimator variances summed across stimulus dimensions, as given by the trace of the local estimator covariance. The expression for the estimator covariance around $\boldsymbol{s}_0$:

$$\left\langle \left(\boldsymbol{W}^\top(\boldsymbol{r} - \boldsymbol{f}(\boldsymbol{s}_0))\right)\left(\boldsymbol{W}^\top(\boldsymbol{r} - \boldsymbol{f}(\boldsymbol{s}_0))\right)^\top \right\rangle_{p(\boldsymbol{r}|\boldsymbol{s}_0)} = \boldsymbol{W}^\top \left\langle (\boldsymbol{r} - \boldsymbol{f}(\boldsymbol{s}_0))(\boldsymbol{r} - \boldsymbol{f}(\boldsymbol{s}_0))^\top \right\rangle_{p(\boldsymbol{r}|\boldsymbol{s}_0)} \boldsymbol{W} = \boldsymbol{W}^\top \boldsymbol{\Sigma} \boldsymbol{W} \tag{11}$$

where $\boldsymbol{\Sigma} = \boldsymbol{\Sigma}(\boldsymbol{s}_0)$ is the noise covariance matrix of the population activity around $\boldsymbol{s}_0$. Then, we aim to solve the following optimization problem:

$$\min \frac{1}{2}\mathrm{Tr}(\boldsymbol{W}^\top \boldsymbol{\Sigma} \boldsymbol{W}) \quad \text{s.t.} \quad \boldsymbol{W}^T \nabla_s \boldsymbol{f}(\boldsymbol{s}) = \boldsymbol{I}, \tag{12}$$

which admits the following Lagrangian

$$\mathcal{L}(\boldsymbol{W}, \boldsymbol{\Lambda}) = \frac{1}{2}\mathrm{Tr}(\boldsymbol{W}^\top \boldsymbol{\Sigma} \boldsymbol{W}) - \mathrm{Tr}(\boldsymbol{\Lambda}^\top(\boldsymbol{W}^\top \nabla_s \boldsymbol{f}(\boldsymbol{s}) - \boldsymbol{I})). \tag{13}$$

The gradients of the Lagrangian with respect to the parameters are given by:

$$\nabla_{\boldsymbol{W}} \mathcal{L}(\boldsymbol{W}, \boldsymbol{\Lambda}) = \frac{1}{2}(\boldsymbol{\Sigma} + \boldsymbol{\Sigma}^\top)\boldsymbol{W} - \nabla_s \boldsymbol{f}(\boldsymbol{s})\boldsymbol{\Lambda}^\top \tag{14}$$

$$\nabla_{\boldsymbol{\Lambda}} \mathcal{L}(\boldsymbol{W}, \boldsymbol{\Lambda}) = \boldsymbol{W}^\top \nabla_s \boldsymbol{f}(\boldsymbol{s}) - \boldsymbol{I} \tag{15}$$

By symmetry of the nosie covariance matrix, $\frac{1}{2}(\boldsymbol{\Sigma} + \boldsymbol{\Sigma}^\top) = \boldsymbol{\Sigma}$. Then the above gradients yield the two optimality conditions

$$\boldsymbol{0} = \boldsymbol{\Sigma}\boldsymbol{W} - \nabla_s \boldsymbol{f}(\boldsymbol{s})\boldsymbol{\Lambda}^\top, \tag{16}$$

$$\boldsymbol{0} = \boldsymbol{W}^\top \nabla_s \boldsymbol{f}(\boldsymbol{s}) - \boldsymbol{I}. \tag{17}$$

From this we see that the optimal $\boldsymbol{W}^* = \boldsymbol{\Sigma}^{-1}\nabla_s \boldsymbol{f}(\boldsymbol{s})\boldsymbol{\Lambda}^\top$. The second condition tells us that $\boldsymbol{W}^\top \nabla_s \boldsymbol{f}(\boldsymbol{s}) = \boldsymbol{I}$, thus confirming unbiasedness. Substituting the optimal $\boldsymbol{W}^*$ into this second condition and solving for $\boldsymbol{\Lambda}$ results in $\boldsymbol{\Lambda}^* = \left(\nabla_s \boldsymbol{f}(\boldsymbol{s})^\top \boldsymbol{\Sigma}^{-1} \nabla_s \boldsymbol{f}(\boldsymbol{s})\right)^{-1}$. Substituting this expression back into the original expression for $\boldsymbol{W}^*$ results in the optimal weights $\boldsymbol{W}^* = \boldsymbol{\Sigma}^{-1}\nabla_s \boldsymbol{f}(\boldsymbol{s}) \left(\nabla_s \boldsymbol{f}(\boldsymbol{s})^\top \boldsymbol{\Sigma}^{-1} \nabla_s \boldsymbol{f}(\boldsymbol{s})\right)^{-1}$. Finally, with some algebra, we find that

$$\boldsymbol{W}^{*\top} \boldsymbol{\Sigma} \boldsymbol{W}^* = \left(\nabla_s \boldsymbol{f}(\boldsymbol{s})^\top \boldsymbol{\Sigma}^{-1} \nabla_s \boldsymbol{f}(\boldsymbol{s})\right)^{-1} = \boldsymbol{\mathcal{I}}(\boldsymbol{s})^{-1}. \tag{18}$$

In other words, the error covariance exactly equals the inverse Fisher information, suggesting that the locally linear estimator $\boldsymbol{W}^*$ is optimal in that it is efficient with respect to the Cramer-Rao bound.

#### 2.2 Population activity model, and some general properties of this model

##### 2.2.1 Population activity and tuning curves

We assume $N$ independent stochastic Poisson spiking neurons with tuning curves $f(\boldsymbol{s}; \Omega_n)$, such that the likelihood of observing population activity vector $\boldsymbol{r}$ for stimulus $\boldsymbol{s}$ is given by

$$p(\boldsymbol{r}|\boldsymbol{s}, \boldsymbol{\Omega}) = \prod_{n=1}^{N} p(r_n|\boldsymbol{s}, \Omega_n) = \prod_{n=1}^{N} \frac{f(\boldsymbol{s}; \Omega_n)^{r_n} \exp\left(-f(\boldsymbol{s}; \Omega_n)\right)}{r_n!}, \tag{19}$$

where $\Omega_n$ is the set of tuning curve parameters for neuron $n$, and $\boldsymbol{\Omega} = \{\Omega_n\}_{n=1}^{M}$ is the set of parameters of all neurons in the population.

The tuning curve of each neuron consist of one or multiple Gaussian-shaped subfields. In general, we assume that each neuron can have an arbitrary maximum number $P$ of subfields (i.e., that their tuning curves are multimodal), such that its tuning is given by

$$f(\boldsymbol{s};\Omega_n) = \sum_{p=1}^{P} f(s,\Omega_{n,p}) = \sum_{p=1}^{P} z_{n,p} a_{n,p} \exp\left(-\frac{1}{2}(\boldsymbol{s}-\boldsymbol{\mu}_{n,p})^\top \boldsymbol{\Sigma}_{n,p}^{-1}(\boldsymbol{s}-\boldsymbol{\mu}_{n,p})\right), \tag{20}$$

where $\Omega_{n,p} = \{z_{n,p}, a_{n,p}, \boldsymbol{\mu}_{n,p}, \boldsymbol{\Sigma}_{n,p}\}$ is the set of tuning curve parameters for the $p$th subfield, and $\Omega_n = \{\Omega_{n,p}\}_{p=1}^{P}$. Each subfield $p$ has gain $a_{n,p} \geq 0$, tuning center $\boldsymbol{\mu}_{n,p} \in \mathbb{R}^D$, and tuning covariance $\boldsymbol{\Sigma}_{n,p} \in \mathbb{R}^{D\times D}$. Furthermore, its binary indicator variable $z_{n,p} \in \{0,1\}$ determines whether the subfield is active ($z_{n,p} = 1$), or inactive ($z_{n,p} = 0$), in which case it doesn't contribute to a neuron's tuning. For instances in which we only consider unimodal tuning curves, we set $P = 1$ and fix $z_{n,1} = 1$, such that each neuron's tuning curve simplifies to

$$f(\boldsymbol{s};\Omega_n) = a_n \exp\left(-\frac{1}{2}(\boldsymbol{s}-\boldsymbol{\mu}_n)^\top \boldsymbol{\Sigma}_n^{-1}(\boldsymbol{s}-\boldsymbol{\mu}_n)\right), \tag{21}$$

where here, to simplify notation, we drop the $p$ index.

##### 2.2.2 Tuning curve parametrization

We assume that tuning curve parameters are drawn independently and identically distributed across neurons and subfields, that is $p(\boldsymbol{\Omega}) = \prod_{n=1,p=1}^{N,P} p(\Omega_{n,p})$. Furthermore, within each subfield, we assume independence across subfield activation, gain, tuning mean, and covariance, that is $p(\Omega_{n,p}) = p(z_{n,p})p(a_{n,p})p(\boldsymbol{\mu}_{n,p})p(\boldsymbol{\Sigma}_{n,p})$. We next specify each of these distributions.

**Subfield activations,** $z_{n,p}$**.** Except for unimodal tuning ($P = 1$), we assume that subfield activations are drawn from a Bernoulli distribution, $z_{n,p} \sim \text{Bern}(\zeta)$, such that subfields are active with probability $\zeta$. Thus, on average, each neuron will have $P\zeta$ active subfields. For unimodal tuning $P = 1$ we instead fix $z_{n,1} = 1$, as already mentioned further above.

**Gains,** $a_{n,p}$**.** Please consult the next section.

**Tuning centers,** $\boldsymbol{\mu}_{n,p}$**.** Receptive field centers are assumed to uniformly cover some square region $[c_{\text{min}}, c_{\text{max}}]^D$, such that the center in the $i$th dimension is drawn from $\mu_{n,p,i} \sim \mathcal{U}[c_{\text{min}}, c_{\text{max}}]$.

**Tuning covariances,** $\boldsymbol{\Sigma}_{n,p}$**.** How we parameterize the tuning covariance is the most critical aspect of our work, as it determines both the heterogeneity of tuning widths varies across neurons (and subfields), and (for $D > 1$) the heterogeneity of tuning shape for each of these subfields. As we justify in a later section, we only consider axis-aligned tuning curves with a diagonal tuning covariance given by $\boldsymbol{\Sigma}_{n,p} = \text{diag}\left(\sigma_{n,p,1}^2, \dots \sigma_{n,p,D}^2\right)$.

First, we describe how we determine shape heterogeneity. For homogeneous tuning shapes, we assume an isotropic covariance, such that $\sigma_{n,p,i} = \sigma_{n,p}$ for all $i = 1,\dots,D$, leading to circular (or, for $D > 2$, spherical/hyperspherical) tuning curves with width $\sigma_{n,p}$. This width is drawn from a Gamma distribution $\sigma_{n,p} \sim \Gamma(\upsilon, \beta/\upsilon)$, such that its mean, $\langle\sigma_{n,p}\rangle = \beta$, is the same across all neurons and subfields. For heterogeneous tuning shapes, we assume an anisotropic covariance where the width for each dimension $i$ is independently drawn from $\sigma_{n,p,i} \sim \Gamma(\upsilon, \beta/\upsilon)$. This leads to tuning shapes that are mostly elliptical (or, for $D > 2$, ellipsoidal/hyper-ellipsoidal), with mean widths $\langle\sigma_{n,p,1}\rangle = \beta$ that are shared across all stimulus dimensions, neurons, and subfields. For $D = 2$ we allow the field width $\sigma_{n,p,1}$ and $\sigma_{n,p,2}$ to be correlated by modeling them as a correlated bivariate Gamma distribution [5] with correlation coefficient $\rho \in [0,1]$, and where $\sigma_1$ and $\sigma_2$ share the same marginal distribution. This allows us to continuously move between the fully heterogeneous case (at $\rho = 0$) and fully homogeneous case (at $\rho = 1$). For $D = 1$, shape heterogeneity does not apply, as tuning curves have a single width $\sigma_{n,p,1}$ that is drawn according to $\sigma_{n,p,1} \sim \Gamma(\upsilon, \beta/\upsilon)$.

For all cases, width heterogeneity is controlled by the shape parameter $\upsilon$ of the Gamma distribution that the individual widths are drawn from. Smaller $\upsilon$'s lead to a larger heterogeneity of tuning curve widths. As special cases, $\upsilon \to 1$ leads to the $\sigma_{n,p,i}$'s to be distributed according to an exponential distribution, whereas, $\upsilon \to \infty$ leads to a fixed-width population, where all tuning curves share the same width, $\beta$. In this limit, the width becomes the same in all dimensions, such that tuning shapes become homogeneous, even if they are drawn from the heterogeneous shape distribution. The choice of a Gamma distribution to model tuning widths in the population is made both to align our work with previous studies [1], and because the Gamma distribution has positive support, and all of its higher-order moments exist, which we find is a necessary condition for computing the FI in closed-form for arbitrary stimulus dimensionality $D$.

In the main text, we express size heterogeneity through a re-parameterization of the Gamma shape parameter, $\upsilon$. Specifically, we introduce, $\alpha = \exp(1-\upsilon)$, so that $\alpha = 0$ corresponds to no size heterogeneity (and is the case of $\upsilon \to \infty$), whereas $\alpha = 1$ corresponds to the maximum degree of size heterogeneity considered in our model ($\upsilon = 1$). While this re-parameterization aids in exposition of the results, we leave the derivations below in terms of $\upsilon$, as this latter parameterization yields more succinct expressions.

##### 2.2.3 Metabolic constraints

Neurons with wider tuning curves are active over a wider range of stimuli, and thus use more metabolic resources than more narrowly tuned neurons. We therefore consider two cases. In the metabolically unconstrained case, this extra metabolic expenditure is ignored, and neural gain for all neurons and subfields is drawn independently from $a_{n,p} \sim \mathcal{U}[g_{\text{mean}} - G, g_{\text{mean}} + G]$ (where $G$ is an arbitrary constant in the range $G \in [0, g_{\text{mean}}]$ so that $\langle a_{n,p} \rangle = g_{\text{mean}}$), irrespective of the width of their tuning curves. For the metabolically constrained case we only consider unimodal tuning, and adjust the gains to meet a fixed metabolic constraint $C$ for each stimulus dimension $D$, that is

$$\left\langle \langle r_n \rangle_{p(r_n|\boldsymbol{s})} \right\rangle_{p(\boldsymbol{s})} = \langle f(\boldsymbol{s}; \Omega_n) \rangle_{p(\boldsymbol{s})} = C^D, \ \forall n. \tag{22}$$

This implies that neural gains scale inversely with their tuning curve widths, that is $a_n = |\boldsymbol{\Sigma}_n|^{-1/2}$. To see why this is the case, note that we can rewrite $f(\boldsymbol{s}; \Omega_n) = a_n \sqrt{(2\pi)^D |\boldsymbol{\Sigma}_n|} \mathcal{N}(\boldsymbol{s}|\boldsymbol{\mu}_n, \boldsymbol{\Sigma}_n)$. Then, $\int a_n \sqrt{(2\pi)^D |\boldsymbol{\Sigma}_n|} \mathcal{N}(\boldsymbol{s}|\boldsymbol{\mu}_n, \boldsymbol{\Sigma}_n) d\boldsymbol{s} = a_n \sqrt{(2\pi)^D |\boldsymbol{\Sigma}_n|} = C^D$, assuming that $\mathcal{N}(\boldsymbol{s}|\boldsymbol{\mu}_n, \boldsymbol{\Sigma}_n) \approx 0$ outside of the considered stimulus range. For this quantity to be constant regardless of tuning curve width, we need to set $a_n \propto |\boldsymbol{\Sigma}_n|^{-1/2}$. As $C$ is an arbitrary overall scaling parameter, we chose $C = \sqrt{2\pi}$ without loss of generality, resulting in $a_n = |\boldsymbol{\Sigma}_n|^{-1/2}$.

##### 2.2.4 The average encoded information equals the average FI per neuron

Our goal is to compute the average (over draws of neural populations, $\boldsymbol{\Omega}$, and across all neurons in each population) FI per neuron in a population of neurons tuned with stochastic parameters. In other words, we are attempting to calculate, $\boldsymbol{\mathcal{I}}(s) = \langle \boldsymbol{\mathcal{I}}(s; \boldsymbol{\Omega})/N \rangle_{p(\boldsymbol{\Omega})}$, where $\boldsymbol{\mathcal{I}}(s; \boldsymbol{\Omega})$ is the FI matrix of population $\boldsymbol{\Omega}$ at stimulus $s$. For independently spiking neurons, the FI of a neural population is the sum of the individual FI's of each neuron in the population, that is $\boldsymbol{\mathcal{I}}(s; \boldsymbol{\Omega}) = \sum_{n=1}^{N} \boldsymbol{\mathcal{I}}_n(s; \Omega_n)$ (Appendix A.1). Then,

$$\boldsymbol{\mathcal{I}}(\boldsymbol{s}) = \frac{1}{N} \langle \boldsymbol{\mathcal{I}}(\boldsymbol{s}; \boldsymbol{\Omega}) \rangle_{p(\boldsymbol{\Omega})} = \frac{1}{N} \left\langle \sum_{n=1}^{N} \boldsymbol{\mathcal{I}}_n(\boldsymbol{s}; \Omega_n) \right\rangle_{p(\boldsymbol{\Omega})} = \frac{1}{N} \sum_{n=1}^{N} \langle \boldsymbol{\mathcal{I}}_n(\boldsymbol{s}; \Omega_n) \rangle_{p(\Omega_n)} = \frac{1}{N} \sum_{n=1}^{N} \boldsymbol{\mathcal{I}}_n(\boldsymbol{s}) = \boldsymbol{\mathcal{I}}_n(\boldsymbol{s}), \tag{23}$$

where we have defined $\boldsymbol{\mathcal{I}}_n(\boldsymbol{s}) = \langle \boldsymbol{\mathcal{I}}_n(\boldsymbol{s}; \Omega_n) \rangle_{p(\Omega_n)}$, and where the final equality follows because all parameters are independently and identically distributed across neurons in the population. This means that to compute the value of interest, we need only to compute $\boldsymbol{\mathcal{I}}_n(s)$, or the FI of a single neuron in the population, averaged over possible tuning curve parameters drawn from $p(\Omega_n)$.

##### 2.2.5 The FI matrix for a single neuron with known tuning parameters

To find the average FI of a single neuron $n$, let us first consider its FI for a given set of parameters $\Omega_n$, which is defined as

$$\boldsymbol{\mathcal{I}}_n(\boldsymbol{s}; \Omega_n) = \left\langle \nabla_{\boldsymbol{s}} \log p(r_n|\boldsymbol{s}, \Omega_n) \nabla_{\boldsymbol{s}} \log p(r_n|\boldsymbol{s}, \Omega_n)^\top \right\rangle_{p(r|\boldsymbol{s}, \Omega_n)}. \tag{24}$$

To compute this, note that

$$\nabla_{\boldsymbol{s}} \log p(r_n|\boldsymbol{s}, \Omega_n) = \nabla_{\boldsymbol{s}} f(\boldsymbol{s}; \Omega_n) \left( \frac{r_n}{f(\boldsymbol{s}; \Omega_n)} - 1 \right). \tag{25}$$

In the multimodal case,

$$\nabla_{\boldsymbol{s}} f(\boldsymbol{s}; \Omega_n) = \sum_{p=1}^{P} f(s, \Omega_{n,p}) \boldsymbol{u}_{n,p} \tag{26}$$

where we have defined $\boldsymbol{u}_{n,p} = \boldsymbol{\Sigma}_{n,p}^{-1}(\boldsymbol{\mu}_{n,p} - \boldsymbol{s})$. Substituting this expression into Eq. (25) and averaging over $p(r_n|s, \Omega_{n,p})$ yields

$$\boldsymbol{\mathcal{I}}_n(\boldsymbol{s}; \Omega_n) = \frac{1}{f(\boldsymbol{s}; \Omega_n)} \sum_{p=1}^{P} \sum_{q=1}^{P} f(\boldsymbol{s}; \Omega_{n,p}) f(s; \Omega_{n,q}) \boldsymbol{u}_{n,p} \boldsymbol{u}_{n,q}^\top, \tag{27}$$

with matrix elements given by

$$\mathcal{I}_{n,i,j}(\boldsymbol{s}; \Omega_n) = \frac{1}{f(s; \Omega_n)} \sum_{p=1}^{P} \sum_{q=1}^{P} \frac{1}{\sigma_{n,p,i}^2 \sigma_{n,q,j}^2} (s_i - \mu_{n,p,i})(s_j - \mu_{n,q,j}) f(\boldsymbol{s}; \Omega_{n,p}) f(\boldsymbol{s}; \Omega_{n,q}). \tag{28}$$

In the unimodal case, Eq. (27) reduces to

$$\mathcal{I}_n(\boldsymbol{s};\Omega_n) = f(\boldsymbol{s};\Omega_n)\boldsymbol{u}_n\boldsymbol{u}_n^\top., \tag{29}$$

with matrix elements

$$\mathcal{I}_{n,i,j}(\boldsymbol{s};\Omega_n) = \frac{1}{\sigma_{n,i}^2\sigma_{n,j}^2}(s_i - \mu_{n,i})(s_j - \mu_{n,j})f(\boldsymbol{s};\Omega_n). \tag{30}$$

For both the unimodal and the multimodal case, average FI of a single neuron, $\mathcal{I}_n(\boldsymbol{s})$, is found by marginalizing the respective expression for $\mathcal{I}_n(\boldsymbol{s};\Omega_n)$ over $p(\Omega_n)$, as we will do in later sections.

###### 2.2.6 Axis-aligned and rotated tuning curves yield the same encoded information

We will generally work with axis-aligned tuning curves. A natural question is how strongly our results depend upon this assumption. Here, we will show that axis-aligned and rotated populations achieve the same Fisher information for general dimensionality $D$ (and thereby the same encoded information). For brevity, we demonstrate this only in the unimodal case, though we verify that this also holds in the multimodal case in simulations. Substituting our expressions for $\boldsymbol{u}_n$ into the FI (in matrix form) for a single neuron for arbitrary tuning covariance $\boldsymbol{\Sigma}_n$, Eq. (29), we get

$$\mathcal{I}_n(\boldsymbol{s};\Omega_n) = f(\boldsymbol{s};\Omega_n)\boldsymbol{\Sigma}_n^{-1}(\boldsymbol{\mu}_n - s)(\boldsymbol{\mu}_n - s)^\top\boldsymbol{\Sigma}_n^{-\top}. \tag{31}$$

We first marginalize over tuning curve centers, $p(\boldsymbol{\mu})$. To do so we note that $f(\boldsymbol{s};\Omega_n) = a_n\sqrt{(2\pi)^D|\boldsymbol{\Sigma}_n|}\mathcal{N}(\boldsymbol{\mu}_n|\boldsymbol{s},\boldsymbol{\Sigma}_n)$, such that

$$\left\langle f(\boldsymbol{s};\Omega_n)(\boldsymbol{\mu}_n - s)(\boldsymbol{\mu}_n - s)^\top\right\rangle_{p(\boldsymbol{\mu})} = a_n\sqrt{(2\pi)^D|\boldsymbol{\Sigma}_n|}\left\langle\mathcal{N}(\boldsymbol{\mu}_n|\boldsymbol{s},\boldsymbol{\Sigma}_n)(\boldsymbol{\mu}_n - s)(\boldsymbol{\mu}_n - s)^\top\right\rangle_{p(\boldsymbol{\mu})} \propto a_n\sqrt{(2\pi)^D|\boldsymbol{\Sigma}_n|}\boldsymbol{\Sigma}_n, \tag{32}$$

where, to avoid boundary effects, we have assumed that these tuning curve centers extend significantly beyond the range of possible stimulus values, and the last expression is proportional up to the $p(\boldsymbol{\mu})$ scaling. As a result, the marginal of the full expression becomes

$$\begin{aligned}\langle\mathcal{I}_n(\boldsymbol{s};\Omega_n)\rangle_{p(\boldsymbol{\mu})} &= \left\langle f(\boldsymbol{s};\Omega_n)\boldsymbol{\Sigma}_n^{-1}(\boldsymbol{\mu} - s)(\boldsymbol{\mu} - s)^\top\boldsymbol{\Sigma}_n^{-\top}\right\rangle_{p(\boldsymbol{\mu})}\\ &= \boldsymbol{\Sigma}_n^{-1}\left\langle f(\boldsymbol{s};\Omega_n)(\boldsymbol{\mu} - s)(\boldsymbol{\mu} - s)^\top\right\rangle_{p(\boldsymbol{\mu})}\boldsymbol{\Sigma}_n^{-\top} \propto \frac{1}{\sqrt{|\boldsymbol{\Sigma}_n^{-1}|}}\boldsymbol{\Sigma}_n^{-1}\boldsymbol{\Sigma}_n\boldsymbol{\Sigma}_n^{-\top} = \frac{\boldsymbol{\Sigma}_n^{-1}}{\sqrt{|\boldsymbol{\Sigma}_n^{-1}|}},\end{aligned} \tag{33}$$

where the final expression is again true up to a multiplicative constant. We will use the remainder of this supplement to evaluate these expressions in closed-form, but for now leave these expressions un-evaluated for ease of exposition.

Axis-aligned tuning curves exhibit diagonal covariance tuning matrices. To achieve rotated tuning curves, we can rewrite the tuning covariance as,

$$\boldsymbol{\Sigma}_n = \boldsymbol{R}\boldsymbol{L}_n\boldsymbol{R}^\top, \qquad \text{such that} \qquad \boldsymbol{\Sigma}_N^{-1} = \boldsymbol{R}^\top\boldsymbol{L}_n^{-1}\boldsymbol{R}, \tag{34}$$

where we have defined $\boldsymbol{L}_n = \text{diag}(\sigma_{n,1}^2,\ldots,\sigma_{n,D}^2)$, and where $\boldsymbol{R}$ is an orthogonal rotation matrix. Then we can rewrite the above expression as

$$\langle\mathcal{I}_n(\boldsymbol{s};\Omega_n)\rangle_{p(\boldsymbol{\mu})} \propto \frac{1}{\sqrt{|\boldsymbol{R}^\top\boldsymbol{L}_n^{-1}\boldsymbol{R}|}}\boldsymbol{R}\boldsymbol{L}_n^{-1}\boldsymbol{R}^\top = \frac{1}{\sqrt{|\boldsymbol{L}_n^{-1}|}}\boldsymbol{R}\boldsymbol{L}_n^{-1}\boldsymbol{R}^\top \tag{35}$$

where the second equality follows from the fact that $\boldsymbol{R}$ is an orthogonal matrix. Now, we marginalize over $p(\sigma)$:

$$\langle\mathcal{I}_n(\boldsymbol{s};\Omega_n)\rangle_{p(\boldsymbol{\mu},\sigma)} \propto \left\langle\frac{1}{\sqrt{|\boldsymbol{L}_n^{-1}|}}\boldsymbol{R}\boldsymbol{L}_n^{-1}\boldsymbol{R}^\top\right\rangle_{p(\sigma)} = \left\langle\boldsymbol{R}\frac{\boldsymbol{L}_n^{-1}}{\sqrt{|\boldsymbol{L}_n^{-1}|}}\boldsymbol{R}^\top\right\rangle_{p(\sigma)} = \boldsymbol{R}\left\langle\frac{\boldsymbol{L}_n^{-1}}{\sqrt{|\boldsymbol{L}_n^{-1}|}}\right\rangle_{p(\sigma)}\boldsymbol{R}^\top \tag{36}$$

We leave the marginalization un-evaluated for now. Except, note that because $\boldsymbol{L}_n$ is a diagonal matrix with either identical elements along the diagonal (isotropic) or i.i.d. elements along the diagonal (anisotropic), the marginalization will yield a scaled version for the identity matrix. We will call this matrix $\boldsymbol{C} = c\boldsymbol{I}$ with the prefactor $c$. Then it is clear that rotations will not impact the FI:

$$\langle\mathcal{I}_n(\boldsymbol{s};\Omega_n)\rangle_{p(\boldsymbol{\mu},\sigma)} \propto \boldsymbol{R}\left\langle\frac{\boldsymbol{L}_n^{-T}}{\sqrt{|\boldsymbol{L}_n^{-1}|}}\right\rangle_{p(\sigma)}\boldsymbol{R}^\top = \boldsymbol{R}\boldsymbol{C}\boldsymbol{R}^\top = c\boldsymbol{R}\boldsymbol{R}^\top \propto \boldsymbol{I}. \tag{37}$$

Given that tuning rotations yield no effect on the FI, and they are cumbersome to work with (especially in higher dimensions), we will proceed working exclusively with axis-aligned tuning curves.

Our derivation shows that the encoded information is independent of how tuning curves are rotated for any stimulus dimensionality. Thus, we are generalizing [1] that used a uniform tuning curve rotations for $D = 2$ only, and did not demonstrate that tuning curve rotations had no impact on encoded information.

### 3 Asymptotic encoded information for general stimulus dimensionality

We consider two cases of how the tuning curve centers cover the stimulus range. We will start with the asymptotic (unbounded) case, which puts no restriction on where the neurons' tuning curve centers can fall relative to the stimuli these populations encode. More specifically, recalling that $\mu_{n,i} \in \mathcal{U}[c_{\min}, c_{\max}]$, we assume that $c_{\min} \ll s_{\min}$ and $c_{\max} \gg s_{\max}$. Later, we will consider the bounded case, in which tuning curve centers can only exceed the stimulus range by finite amounts.

For the unbounded case, we examine an adjusted average FI that takes into account that the tuning curve centers may span an infinitely large range, but the stimulus range is fixed: We compute $\hat{\mathcal{I}}/\eta$ where $\eta = \left(\frac{s_{\max}-s_{\min}}{c_{\max}-c_{\min}}\right)^D$ and where $\eta$ can be thought of as the fraction of stimulus-responsive neurons in the population. In what follows, we will drop the $\cdot_n$ subscript from the various tuning curve parameters to keep the notation uncluttered.

#### 3.1 Unimodal tuning curves

We will start with the simpler unimodal case. We begin by marginalizing Eq. (30) over tuning curve centers, $p(\boldsymbol{\mu})$:

$$\begin{aligned} \langle \boldsymbol{\mathcal{I}}_{n,i,j}(\boldsymbol{s};\Omega_n)\rangle_{p(\boldsymbol{\mu})} &= \frac{1}{\eta}\left(\frac{1}{(c_{\text{min}}-c_{\text{max}})^D}\int_{c_{\text{min}}}^{c_{\text{max}}} \boldsymbol{\mathcal{I}}_{n,i,j}(\boldsymbol{s};\Omega_n)d\boldsymbol{\mu}\right) \\ &= \frac{1}{(s_{\text{min}}-s_{\text{max}})^D}\int_{c_{\text{min}}}^{c_{\text{max}}} \frac{1}{\sigma_i^2\sigma_j^2}(s_i-\mu_i)(s_j-\mu_j)f(\boldsymbol{s};\Omega_n)d\boldsymbol{\mu}. \end{aligned} \tag{38}$$

In the limit $(c_{\text{min}}, c_{\text{max}}) \to (-\infty, \infty)$, the above becomes

$$\begin{aligned} \langle \boldsymbol{\mathcal{I}}_{n,i,j}(\boldsymbol{s};\Omega_n)\rangle_{p(\boldsymbol{\mu})} &= \frac{1}{(s_{\text{min}}-s_{\text{max}})^D}\int_{-\infty}^{\infty} \frac{1}{\sigma_i^2\sigma_j^2}(s_i-\mu_i)(s_j-\mu_j)f(\boldsymbol{s};\Omega_n)d\boldsymbol{\mu} \\ &= \frac{a(2\pi)^{D/2}\Sigma_{i,j}}{\sigma_i^2\sigma_j^2\sqrt{|\boldsymbol{\Sigma}^{-1}|}(s_{\max}-s_{\min})^D}. \end{aligned} \tag{39}$$

The tuning curve covariance matrix $\boldsymbol{\Sigma}$ is diagonal, such that only the diagonal terms of the FI matrix are non-zero, and are given by

$$\boldsymbol{\mathcal{I}}_{n,i,i}(\boldsymbol{s};a,\sigma_1,\cdots,\sigma_D) = \frac{a(2\pi)^{D/2}\prod_{k=1}^D\sigma_k}{\sigma_i^2(s_{\max}-s_{\min})^D} = \frac{a(2\pi)^{D/2}\prod_{k\neq i}\sigma_k}{\sigma_i(s_{\max}-s_{\min})^D}, \tag{40}$$

where we have used $1/\sqrt{|\boldsymbol{\Sigma}^{-1}|} = \sqrt{|\boldsymbol{\Sigma}|} = \sqrt{\prod_{k=1}^D \sigma_k^2} = \prod_{k=1}^D \sigma_k$.

**Maximal shape heterogeneity (anisotropic tuning covariance).** Averaging next over the independently drawn $\sigma_1, \cdots, \sigma_D$ results in

$$\langle \boldsymbol{\mathcal{I}}_{n,i,i}(\boldsymbol{s};a,\sigma_1,\cdots,\sigma_D)\rangle_{p(\sigma_1,\cdots,\sigma_D)} = \frac{a(2\pi)^{D/2}}{(s_{\max}-s_{\min})^D}\left\langle \frac{1}{\sigma_i}\left(\prod_{j\neq i}\sigma_j\right)\right\rangle_{p(\sigma_1,\cdots,\sigma_D)} = \frac{a(2\pi)^{D/2}\beta^{D-2}}{(s_{\max}-s_{\min})^D}\left(\frac{\upsilon}{\upsilon-1}\right). \tag{41}$$

Finally, averaging over gains results in,

$$\boxed{\hat{\mathcal{I}}_{H,A} = K_D\frac{\upsilon}{\upsilon-1}, \qquad K_D = \frac{(2\pi)^{D/2}\beta^{D-2}g_{\text{mean}}}{(s_{\max}-s_{\min})^D}}, \tag{42}$$

where the $\cdot_D$ subscript on $K$ indicates that the term is dimensionality-dependent. In the fixed width limit, $\upsilon \to \infty$, the encoded information reduces to

$$\boxed{\hat{\mathcal{I}}_F = K_D} \tag{43}$$

So, we find that the information gain of having heterogeneously sized tuning widths in the population is given by:

$$\boxed{\frac{\hat{\mathcal{I}}_{H,A}}{\hat{\mathcal{I}}_F} = \frac{\upsilon}{\upsilon-1}\frac{K_D}{K_D} = \frac{\upsilon}{\upsilon-1}}, \tag{44}$$

independent of dimensionality.

It is important to observe at this point that the $\upsilon \to 1$ limit turns the Gamma distribution underlying the distribution of tuning curve widths into an exponential distribution, like the one assumed in [1]. In this limit, however, the integral $\langle 1/\sigma_i\rangle$ that is required to take the average over these tuning curve widths does not converge. This is also reflected in the above FI expressions which go to infinity in this limit. Thus, any attempt to approximate this integral by Monte Carlo methods, as done in [1], will lead to numerical instabilities, and potentially spurious result.

**No shape heterogeneity (isotropic tuning covariance).** Returning to Eq. (41) and averaging additionally over tuning curve gains,

$$\langle \boldsymbol{\mathcal{I}}_{n,i,i}(\boldsymbol{s}; a, \sigma) \rangle_{p(\sigma,a)} = \frac{\langle a \rangle_{p(a)} (2\pi)^{D/2}}{(s_{\max} - s_{\min})^D} \left\langle \frac{1}{\sigma} \left( \prod_{j \neq i} \sigma \right) \right\rangle_{p(\sigma)} = K_D \frac{\Gamma(\upsilon + (D-2))}{\Gamma(\upsilon)\upsilon^{D-2}}, \tag{45}$$

where $K_D$ is the same as in Eq. (42), and where we have used the general expression for the moments of a Gamma distribution: $\langle \sigma^k \rangle = (\beta/\upsilon)^k \frac{\Gamma(\upsilon+k)}{\Gamma(\upsilon)}$. So that,

$$\boxed{\hat{\mathcal{I}}_{H,I} = K_D \frac{\Gamma(\upsilon + (D-2))}{\Gamma(\upsilon)\upsilon^{D-2}}} \tag{46}$$

Comparing this expression to Eq. (42) shows that the seemingly minor assumption of equal field widths across all dimensions has a strong effect on how the population's information depends on tuning heterogeneity.

In particular, in contrast to Eq. (44), the information gain is now dimensionality-dependent:

$$\boxed{\frac{\hat{\mathcal{I}}_{H,I}}{\hat{\mathcal{I}}_F} = \frac{\Gamma(\upsilon + (D-2))}{\Gamma(\upsilon)\upsilon^{D-2}} \frac{K_D}{K_D} = \frac{\Gamma(\upsilon + (D-2))}{\Gamma(\upsilon)\upsilon^{D-2}}} \tag{47}$$

The difference in information gain when compared to anisotropic tuning covariances is particularly stark for $D = 2$ and $D = 3$. For anisotropic tuning curves, the information gain, Eq. (44) is larger for more homogeneous tuning widths and approaches infinity with $\upsilon \to 1$. For isotropic tuning covariances, the information gain for $D = 2$ becomes

$$\frac{\hat{\mathcal{I}}_{H,I}}{\hat{\mathcal{I}}_F} \overset{D=2}{=} \frac{\Gamma(\upsilon)}{\Gamma(\upsilon)} = 1. \tag{48}$$

and, indeed, for $D = 3$:

$$\frac{\hat{\mathcal{I}}_{H,I}}{\hat{\mathcal{I}}_F} \overset{D=3}{=} \frac{\Gamma(\upsilon + 1)}{\Gamma(\upsilon)\upsilon} = 1 \tag{49}$$

where the final equality follows from the definition of the Gamma function: $\Gamma(\upsilon + 1) = \upsilon\Gamma(\upsilon)$. That is, in the absence of tuning shape heterogeneity, the heterogeneity of place field sizes ceases to influence the encoded information. For both of these cases, an information gain subject to heterogeneity is only possible if tuning curves exhibit broad shape heterogeneity (anisotropic). We will investigate this phenomenon further in the following subsection.

**Continuously varying the shape heterogeneity for $D = 2$.** For populations encoding 2$D$ stimuli, we can further characterize the joint impact of heterogeneity in both field shape and size. Recall that in the $D = 2$ case only, $\sigma_1$ and $\sigma_2$ are distributed as a correlated bivariate Gamma distribution, with correlation coefficient $\rho$. When $D = 2$, Eq. (41) becomes

$$\langle \boldsymbol{\mathcal{I}}_{n,1,1}(\boldsymbol{s}; a, \sigma_1, \sigma_2) \rangle_{p(\sigma_1,\sigma_2)} = \frac{2\pi a}{(s_{\max} - s_{\min})^2} \left\langle \frac{\sigma_2}{\sigma_1} \right\rangle_{p(\sigma_1,\sigma_2)} \tag{50}$$

Define $\iota \doteq \sigma_1/\sigma_2$. From [5], we know that

$$\langle \iota \rangle = \frac{\Gamma(\upsilon + 1)\Gamma(\upsilon - 1)}{\Gamma(\upsilon)\Gamma(\upsilon)} \, {}_2F_1(-1, 1; \upsilon; \rho) \tag{51}$$

where ${}_2F_1(\cdot)$ is the Gaussian hypergeometric function. Noting that $\Gamma(\upsilon) = (\upsilon - 1)\Gamma(\upsilon - 1)$, $\Gamma(\upsilon + 1) = \upsilon\Gamma(\upsilon)$, and that ${}_2F_1(-1, 1; \upsilon; \rho) = \frac{\upsilon - \rho}{\upsilon}$, we obtain that

$$\langle \boldsymbol{\mathcal{I}}_{n,1,1}(\boldsymbol{s}; a, \sigma_1, \sigma_2) \rangle_{p(\sigma_1,\sigma_2)} = \langle \boldsymbol{\mathcal{I}}_{n,2,2}(\boldsymbol{s}; a, \sigma_1, \sigma_2) \rangle_{p(\sigma_1,\sigma_2)} = \frac{2\pi a}{(s_{\max} - s_{\min})^2} \left( \frac{\upsilon - \rho}{\upsilon - 1} \right). \tag{52}$$

Performing the final marginalization over $p(a)$ and using Eq. (1), we find that

$$\boxed{\hat{\mathcal{I}}_{H|D=2} = K_2 \left( \frac{\upsilon - \rho}{\upsilon - 1} \right)} \tag{53}$$

where $K_2$ is the constant $K_D$ defined above in the $D = 2$ case. As before, in the fixed-width limit,

$$\boxed{\hat{\mathcal{I}}_{F|D=2} = K_2} \tag{54}$$

So that in general, the information gain in the $D = 2$ case is

$$\boxed{\frac{\hat{\mathcal{I}}_{H|D=2}}{\hat{\mathcal{I}}_{F|D=2}} = \frac{K_2}{K_2}\left(\frac{\upsilon - \rho}{\upsilon - 1}\right) = \frac{\upsilon - \rho}{\upsilon - 1}.} \tag{55}$$

This expression allows us to understand the information gain in populations as a function of two smoothly varying parameters controlling field size and shape heterogeneity ($\upsilon$ and $\rho$, respectively). Note that for $\rho \to 0$, when the receptive fields widths are uncorrelated across spatial dimensions,

$$\hat{\mathcal{I}}_{H|D=2,\rho=0} = K_2\left(\frac{\upsilon}{\upsilon - 1}\right) \tag{56}$$

Because $\upsilon \geq 1$, the prefactor $\frac{\upsilon}{\upsilon-1} > 1$ (except when $\upsilon \to \infty$ when the prefactor is equal to 1), and is largest as $\upsilon \to 1$.

Conversely, when $\rho \to 1$, where there is no shape heterogeneity (but nonetheless heterogeneous in size across the population),

$$\hat{\mathcal{I}}_{H|D=2,\rho=1} = K_2 = \hat{\mathcal{I}}_{F|D=2}. \tag{57}$$

which exactly coincides with the fixed-width case. In other words, for any strength of heterogeneity in population field sizes, as the correlation between field radii across spatial dimensions for individual neurons approaches 1 (i.e., they become equal), the computational advantage of heterogeneously sized fields across the population vanishes. This matches our results in the previous section.

**Information gain does not depend on the chosen distribution type (general $D$).** We have so far assumed that the field widths, $\sigma_1, \ldots, \sigma_D$ are drawn from a Gamma distribution. We here show that the information gain does not depend on the specific choice of distribution. In other words, encoded information benefits from receptive field heterogeneity irrespective of how exactly the receptive fields vary in shape and size.

To show this, let us start with the case of maximal shape heterogeneity, for which the field width's $\sigma_i$ are drawn independently across the different stimulus dimensions. In this case, by Eq. (41) tells us that the information gain is given by

$$\frac{\hat{\mathcal{I}}_{H,A}}{\hat{\mathcal{I}}_F} = \frac{\left\langle\frac{1}{\sigma}\right\rangle \prod_{j\neq i}\langle\sigma\rangle}{\frac{1}{\beta}\prod_{j\neq i}\beta} = \left\langle\frac{1}{\sigma}\right\rangle\beta. \tag{58}$$

This expression arises from the fact that the dimension-normalized Fisher information is the geometric mean across the information along each stimulus dimension $i$, and that, furthermore, the per-dimension information is the same across all dimensions. Furthermore, for $\hat{\mathcal{I}}_F$ we have used the fact that $\sigma \to \beta$ in the limit of no heterogeneity, and, by definition $\langle\sigma\rangle = \beta$, such that the two products cancel each other. Lastly, as $1/\sigma$ is a strictly convex function in $\sigma$ we have that $\langle 1/\sigma\rangle > 1/\langle\sigma\rangle = 1/\beta$ (assuming that $p(\sigma)$ is not a delta function). As a results $\langle 1/\sigma\rangle\,\beta > 1$, such that

$$\frac{\hat{\mathcal{I}}_{H,A}}{\hat{\mathcal{I}}_F} > 1, \tag{59}$$

irrespective of the exact shape of $p(\sigma)$, as long as it is not a delta function. Once it becomes a delta function, the $\sigma$'s become constants, such that the information gain becomes one.

For no shape heterogeneity we have $\sigma_1 = \cdots = \sigma_D = \sigma$. In this case, Eq. (45) tells us that the information gain is given by

$$\frac{\hat{\mathcal{I}}_{H,I}}{\hat{\mathcal{I}}_F} = \frac{\left\langle\frac{1}{\sigma}\prod_{j\neq i}\sigma\right\rangle}{\frac{1}{\beta}\prod_{j\neq i}\beta} = \frac{\left\langle\sigma^{D-2}\right\rangle}{\beta^{D-2}}. \tag{60}$$

It is easy to see that $\sigma^{D-2}$ is strictly convex for non-negative $\sigma$ if $D = 1$ and $D > 3$. For these cases, we thus have $\left\langle\sigma^{D-2}\right\rangle > \langle\sigma\rangle^{D-2} = \beta^{D-2}$, where we again have assumed that $p(\sigma)$ is not a delta function. For $D = 2$ we get $\left\langle\sigma^{D-2}\right\rangle = 1 = \beta^{D-2}$. For $D = 3$ we have $\left\langle\sigma^{D-2}\right\rangle = \langle\sigma\rangle = \beta = \beta^{D-2}$. Taken together, this implies

$$\frac{\hat{\mathcal{I}}_{H,I}}{\hat{\mathcal{I}}_F} = 1 \quad \text{if } D \in \{2,3\}, \qquad \frac{\hat{\mathcal{I}}_{H,I}}{\hat{\mathcal{I}}_F} > 1 \quad \text{otherwise}, \tag{61}$$

irrespective of the exact shape of $p(\sigma)$ as long as it is not a delta function. Once it becomes a delta function, the information gain becomes one for all $D$.

**Information gain due to shape heterogeneity does not depend on the chosen distribution type ($D = 2$).** Let us now focus on the contribution of shape heterogeneity to information gain for $D = 2$. For this case we have previously assumed that the per-dimension widths of neurons' receptive fields are drawn from a correlated bivariate Gamma distribution with correlation strength $\rho$. We see in this case that decorrelating the principal widths leads to higher encoded information. Let us now show that this information gain due to shape heterogeneity does not depend on the specific type of distribution we have chosen. As we have seen in the previous section, the information gain due to size heterogeneity alone (i.e., no shape heterogeneity, $\rho = 1$) is one for $D = 2$, irrespective of the shape of $p(\sigma)$. Furthermore, the information gain for maximal shape heterogeneity (i.e., $\rho = 0$) is above one for $D = 2$, again irrespective of the shape of $p(\sigma)$. This alone already tells us that shape heterogeneity guarantees an increase for $D = 2$, irrespective of the shape of $p(\sigma)$.

To gain further insight into how encoded information depends on $\rho$ between the two extremes, $\rho \in [0, 1]$, note that, by Eq. (50), encoded information depends on the average ratio of these field widths, that is $\hat{\mathcal{I}} \propto \langle \sigma_1/\sigma_2 \rangle_{p(\sigma_1,\sigma_2)}$. We then observe that an approximate Taylor expansion of the ratio of two random variables tells us that

$$\left\langle \frac{\sigma_1}{\sigma_2} \right\rangle_{p(\sigma_1,\sigma_2)} \approx \frac{\langle \sigma_1 \rangle_{p(\sigma_1)}}{\langle \sigma_2 \rangle_{p(\sigma_2)}} - \frac{\text{Cov}(\sigma_1, \sigma_2)}{\langle \sigma_2 \rangle^2_{p(\sigma_2)}} + \frac{\langle \sigma_1 \rangle_{p(\sigma_1)}}{\langle \sigma_2 \rangle^3_{p(\sigma_2)}} \text{Var}(\sigma_2), \tag{62}$$

regardless of the specific choice of $p(\sigma_1, \sigma_2)$. Assuming that the marginal distributions of $\sigma_1$ and $\sigma_2$ are identical, such that $\langle \sigma_1 \rangle_{p(\sigma_1)} = \langle \sigma_2 \rangle_{p(\sigma_2)}$, $\text{Var}(\sigma_1) = \text{Var}(\sigma_2)$, and $\text{Cov}(\sigma_1, \sigma_2) = \rho\sqrt{\text{Var}(\sigma_1)\text{Var}(\sigma_2)} = \rho\text{Var}(\sigma_2)$,

$$\begin{aligned} \left\langle \frac{\sigma_1}{\sigma_2} \right\rangle_{p(\sigma_1,\sigma_2)} &\approx \frac{\langle \sigma_1 \rangle_{p(\sigma_1)}}{\langle \sigma_2 \rangle_{p(\sigma_2)}} - \frac{\rho\text{Var}(\sigma_2)}{\langle \sigma_2 \rangle^2_{p(\sigma_2)}} + \frac{\text{Var}(\sigma_2)}{\langle \sigma_2 \rangle^2_{p(\sigma_2)}} \\ &= 1 + \frac{\text{Var}(\sigma_1)}{\langle \sigma_1 \rangle^2_{p(\sigma_1)}}(1 - \rho) \end{aligned} \tag{63}$$

Note that the ratio $\frac{\text{Var}(\sigma_1)}{\langle \sigma_1 \rangle^2_{p(\sigma_1)}} > 0$ is positive as long as the distribution has positive support. This is a mild, sensible assumption about the distribution of receptive field widths, which are necessarily nonnegative. This result therefore tells us that decorrelating principal widths *always* improves encoded information, regardless of the choice of $p(\sigma_1, \sigma_2)$. It also tells us that the amount of information gained is approximately[1] linearly proportional to the ratio of the marginal width variance and the square of the average marginal width, where the proportionality constant linearly increases as the correlation strength between principal widths decreases.

Intriguingly, this tells us that generally, populations with (i) maximally decorrelated principal widths ($\rho = 0$), and which (ii) have maximally variable marginal widths ($\text{Var}(\sigma)$) while maintaining minimal marginal average field width can be expected to achieve the highest encoded information.

#### 3.2 Multimodal tuning curves

For the multimodal case, we will in most cases omit the $\cdot_n$ index on parameters to keep the notation uncluttered, but will retain the subfield index, $\cdot_p$, to avoid confusion. Unlike the unimodal case, the normalizing factor for the multimodal case is $\eta = (s_{\max} - s_{\min})^D$. We will discuss why this is the case below. We begin, as above, by averaging over tuning curve centers:

$$\begin{aligned} &\langle \boldsymbol{\mathcal{I}}_{n,i,j}(\boldsymbol{s}; \Omega_n) \rangle_{p(\boldsymbol{\mu})} = \\ &\frac{1}{\eta}\left( \frac{1}{(c_{\max} - c_{\min})^{2D}} \int_{c_{\min}}^{c_{\max}} \int_{c_{\min}}^{c_{\max}} \frac{1}{f(\boldsymbol{s}; \Omega_n)} \sum_{p=1}^{P} \sum_{q=1}^{P} \frac{1}{\sigma^2_{p,i}\sigma^2_{q,j}} (s_i - \mu_{p,i})(s_j - \mu_{q,j}) f(\boldsymbol{s}; \Omega_{n,p}) f(\boldsymbol{s}; \Omega_{n,q}) d\boldsymbol{\mu}_p d\boldsymbol{\mu}_q \right) \end{aligned} \tag{64}$$

Note that such averaging would seem more complicated than in the unimodal case: while the product of two Gaussian densities ($f(\boldsymbol{s}; \Omega_{n,p}), f(\boldsymbol{s}; \Omega_{n,q})$) is still proportional to a Gaussian density, we now must rescale everything by $\frac{1}{f(\boldsymbol{s};\Omega_n)}$, leading to complications in marginalizing over $p(\boldsymbol{\mu})$. To circumvent this, we make a simplifying assumption: We assume that the possible number of subfields per-neuron is quite large, or $P \to \infty$. In this case,

$$f(\boldsymbol{s}; \Omega_n) = \frac{P}{P} f(\boldsymbol{s}; \Omega_n) = P\left( \frac{1}{P} \sum_{p=1}^{P} f(\boldsymbol{s}; \Omega_{n,p}) \right) \xrightarrow[P \to \infty]{} P \left\langle f(\boldsymbol{s}; \Omega_{n,p}) \right\rangle_{p(\Omega_{n,p})} \equiv P\chi(\boldsymbol{s}) \tag{65}$$

where $\chi(\boldsymbol{s}) = \langle f(\boldsymbol{s}; \Omega_{n,p}) \rangle_{p(\Omega_n)}$. In other words, as the number of subfields becomes large, the normalizing factor $\frac{1}{f(\boldsymbol{s};\Omega_n)}$ becomes independent of the parameters of individual subfields and is merely dependent on $\boldsymbol{s}$. This permits us to ignore

[1] We emphasize that understanding the impact of anisotropy for the case of general $p(\sigma_1, \sigma_2)$ and $\rho \in [0, 1]$ is found by an approximation of the true encoded information. Indeed, if we plug in the known quantities for the bivariate correlated Gamma case into Eq. (63), we find that $\hat{\mathcal{I}} \propto 1 + \frac{1-\rho}{v}$, which differs from Eq. (55). Importantly, though, the approximation agrees that encoded information is linear in $(1 - \rho)$.

it while marginalizing over the subfield parameters. This assumption may be interpreted as meaning that each individual neuron has roughly constant activity over the entire stimulus range, which seems to violate the intuitive notion of a receptive field in the first place. Thus, these derivations should only be understood as an asymptotic approximation of the encoded information contained in populations with multimodal tuning curves. That said, we find in simulations that the theory is a valid approximation satisfied by assuming only $P = 20$ subfields, and with a field rate of $\zeta = 0.5$, such that on average, each neuron has only 10 active subfields (Extended Data Fig. 2). In the asymptotic case, $\chi(s)$ can easily computed by first marginalizing over $p(\boldsymbol{\mu})$ and then over $p(a)$ and $p(v)$:

$$\begin{aligned}
\langle f(\boldsymbol{s};\Omega_{n,p})\rangle_{p(\boldsymbol{\mu})} &= \frac{1}{(c_{\max}-c_{\min})^D}\int_{c_{\min}}^{c_{\max}} f(\boldsymbol{s};\Omega_{n,p})d\boldsymbol{\mu} = \frac{(2\pi)^{D/2}}{(c_{\max}-c_{\min})^D}\frac{a_n z_n}{\sqrt{|\boldsymbol{\Sigma}^{-1}|}} = \frac{(2\pi)^{D/2}}{(c_{\max}-c_{\min})^D}a_{n,p}z_{n,p}\sqrt{|\boldsymbol{\Sigma}|} \\
\langle f(\boldsymbol{s};a,z,\sigma_1,\cdots\sigma_D)\rangle_{p(a,v)} &= \frac{(2\pi)^{D/2}}{(c_{\max}-c_{\min})^D}\langle a_n\rangle_{p(a)}\langle z_n\rangle_{p(z)}\sqrt{|\boldsymbol{\Sigma}|} = \frac{(2\pi)^{D/2}g_{\text{mean}}\zeta}{(c_{\max}-c_{\min})^D}\sqrt{|\boldsymbol{\Sigma}|} \\
\chi(\boldsymbol{s}) &= \frac{(2\pi)^{D/2}g_{\text{mean}}\zeta}{(c_{\max}-c_{\min})^D}\left\langle\sqrt{|\boldsymbol{\Sigma}|}\right\rangle_{p(\sigma)}.
\end{aligned} \tag{66}$$

For now, we will skip the final marginalization over $p(\sigma)$, as it depends on specific field shape heterogeneity across the population. Notably, $\chi(s)$ is independent of $s$, and so we will drop its explicit dependence on $s$. Returning to Eq. (64), and taking the $(c_{\text{min}}, c_{\text{max}}) \to (-\infty,\infty)$ limit:

$$\begin{aligned}
\langle \mathcal{I}_{n,i,j}(\boldsymbol{s};\Omega_n)\rangle_{p(\boldsymbol{\mu})} &= \frac{1}{P\bar{\chi}(s_{\max}-s_{\min})^D}\int_{-\infty}^{\infty}\int_{-\infty}^{\infty}\sum_{p=1}^{P}\sum_{q=1}^{P}\frac{1}{\sigma_{p,i}^2\sigma_{q,j}^2}(s_i-\mu_{p,i})(s_j-\mu_{q,j})f(\boldsymbol{s};\Omega_{n,p})f(\boldsymbol{s};\Omega_{n,q})d\boldsymbol{\mu}_p d\boldsymbol{\mu}_q \\
&= \frac{1}{P\bar{\chi}(s_{\max}-s_{\min})^D}\Bigg(\sum_{p=1}^{P}\int_{-\infty}^{\infty}\frac{1}{\sigma_{p,i}^2\sigma_{p,j}^2}(s_i-\mu_{p,i})(s_j-\mu_{p,j})f(\boldsymbol{s};\Omega_{n,p})^2 d\boldsymbol{\mu}_p + \\
&\qquad \frac{1}{(c_{\max}-c_{\min})^D}\sum_{p\neq q}\int_{-\infty}^{\infty}\int_{-\infty}^{\infty}\frac{1}{\sigma_{p,i}^2\sigma_{q,j}^2}(s_i-\mu_{p,i})(s_j-\mu_{q,j})f(\boldsymbol{s};\Omega_{n,p})f(\boldsymbol{s};\Omega_{n,q})d\boldsymbol{\mu}_p d\boldsymbol{\mu}_q\Bigg)
\end{aligned} \tag{67}$$

where $\bar{\chi} = (2\pi)^{D/2}g_{\text{mean}}\zeta\left\langle\sqrt{|\boldsymbol{\Sigma}|}\right\rangle_{p(\sigma)}$ (where the $(c_{\max}-c_{\min})^{-D}$ fully cancels out the $(c_{\max}-c_{\min})^D$ normalizing factor contained in the marginalization over $d\boldsymbol{\mu}_p$ and partially cancels out the $(c_{\max}-c_{\min})^{2D}$ normalization factor in the pairwise field marginalization term). This is the reason for the change of the normalizing factor $\eta$ for the multimodal case. In the unimodal case, $\eta$ helps to normalize the encoded information, given the assumption of tuning curve centers that span far beyond the stimulus range. In the multimodal case, this normalization is effectively performed by $\chi(s)$, such that we need not account for it explicitly through $\eta$. Note that the cross-subfield terms in the $\sum_{p\neq q}\cdot$ sum will always be zero: The centers of each subfield are independently sampled, allowing us to re-arrange the integrals above such that each $p \neq q$ term is zero:

$$\begin{aligned}
&\sum_{p\neq q}\int_{-\infty}^{\infty}\int_{-\infty}^{\infty}\frac{1}{\sigma_{p,i}^2\sigma_{q,j}^2}(s_i-\mu_{p,i})(s_j-\mu_{q,j})f(\boldsymbol{s};\Omega_{n,p})f(\boldsymbol{s};\Omega_{n,q})d\boldsymbol{\mu}_p d\boldsymbol{\mu}_q\Big) = \\
&\qquad \sum_{p\neq q}\left(\int_{-\infty}^{\infty}\frac{(s_i-\mu_{p,i})}{\sigma_{p,i}^2}f(\boldsymbol{s};\Omega_{n,p})d\boldsymbol{\mu}_p\right)\left(\int_{-\infty}^{\infty}\frac{(s_j-\mu_{q,j})}{\sigma_{q,j}^2}f(\boldsymbol{s};\Omega_{n,q})d\boldsymbol{\mu}_q\right) = 0.
\end{aligned} \tag{68}$$

Here, the integrals over $\boldsymbol{\mu}_p$ and $\boldsymbol{\mu}_q$ cause $\mu_{p,i}$ and $\mu_{p,j}$ to become $s_i$ and $s_j$, respectively, causing each of the brackets to evaluate to zero. As such, we can ignore the interaction terms between the subfields. Then,

$$\begin{aligned}
\langle \mathcal{I}_{n,i,j}(\boldsymbol{s};\Omega_n)\rangle_{p(\boldsymbol{\mu})} &= \frac{1}{P\bar{\chi}(s_{\max}-s_{\min})^D}\left(\sum_{p=1}^{P}\int_{-\infty}^{\infty}\frac{1}{\sigma_{p,i}^2\sigma_{p,j}^2}(s_i-\mu_{p,i})(s_j-\mu_{p,j})f(\boldsymbol{s};\Omega_{n,p})^2 d\boldsymbol{\mu}_p\right) \\
&= \frac{1}{\bar{\chi}(s_{\max}-s_{\min})^D}\int_{-\infty}^{\infty}\frac{1}{\sigma_{p,i}^2\sigma_{p,j}^2}(s_i-\mu_{p,i})(s_j-\mu_{p,j})f(\boldsymbol{s};\Omega_{n,p})^2 d\boldsymbol{\mu}_p \\
&= \frac{\pi^{D/2}a_p^2 z_p^2}{2\bar{\chi}(s_{\max}-s_{\min})^D}\frac{\sqrt{|\boldsymbol{\Sigma}_p|}}{\sigma_{p,i}^2\sigma_{p,j}^2}\Sigma_{p|i,j}
\end{aligned} \tag{69}$$

where the second-to-last line follows from the fact that the subfield parameters are identically distributed, so that the sum of the average over the $P$ subfields is equal to $P$ times the average of one subfield. The final line follows from observing that $f(\boldsymbol{s};\Omega_n,p)^2 = a_p^2 z_p^2\sqrt{\pi^D|\boldsymbol{\Sigma}_p|}\mathcal{N}(\boldsymbol{s}|\boldsymbol{\mu}_p,\boldsymbol{\Sigma}_p/2)$, such that the result of the marginalization is proportional to $\Sigma_{p|i,j}$. Proceeding forward, as there are no interaction terms between subfields, we can now drop the subfield index $\cdot_p$. The tuning curves are axis-aligned, such that $\mathbf{\Sigma}_p$ is diagonal. Consequently, only the diagonal elements of $\langle \mathcal{I}_{n,i,j}(\boldsymbol{s};\Omega_n)\rangle_{p(\boldsymbol{\mu})}$ are non-zero, and are given by:

$$\langle \mathcal{I}_{n,i,i}(\boldsymbol{s};\Omega_n)\rangle_{p(\boldsymbol{\mu})} = \frac{\pi^{D/2}a^2 z^2}{2\bar{\chi}(s_{\max}-s_{\min})^D}\frac{\sqrt{|\mathbf{\Sigma}|}}{\sigma_i^4}\Sigma_{i,i} \tag{70}$$

Marginalizing over $p(a)$ and $p(z)$ yields

$$\langle \mathcal{I}_{n,i,i}(\boldsymbol{s};a,z,\sigma_1,\cdots,\sigma_D)\rangle_{p(a,z)} = \frac{\pi^{D/2}\left\langle a^2\right\rangle_{p(a)}\left\langle z^2\right\rangle_{p(z)}}{2\bar{\chi}(s_{\max}-s_{\min})^D}\frac{\sqrt{|\mathbf{\Sigma}|}}{\sigma_i^4}\Sigma_{i,i} = \frac{\pi^{D/2}\left(g_{\text{mean}}^2+\frac{G^2}{3}\right)\zeta}{2\bar{\chi}(s_{\max}-s_{\min})^D}\frac{\sqrt{|\mathbf{\Sigma}|}}{\sigma_i^4}\Sigma_{i,i}, \tag{71}$$

which, using $\sqrt{|\mathbf{\Sigma}|}=\prod_{j=1}^{D}\sigma_j$ and $\Sigma_{i,i}=\sigma_i^2$ can be further simplified to

$$\mathcal{I}_{n,i,i}(\boldsymbol{s};\sigma_1,\cdots,\sigma_D) = \frac{\pi^{D/2}\left(g_{\text{mean}}^2+\frac{G^2}{3}\right)\zeta}{2\bar{\chi}(s_{\max}-s_{\min})^D}\frac{\prod_{j\neq i}\sigma_j}{\sigma_i}. \tag{72}$$

All that remains is to marginalize over $p(\sigma)$, which we will now do for different tuning covariance assumptions.

**No shape heterogeneity (isotropic tuning covariance).** We evaluate the remaining component of $\bar{\chi}$ and the FI terms separately:

$$\bar{\chi} = (2\pi)^{D/2}g_{\text{mean}}\zeta\left\langle\sqrt{|\mathbf{\Sigma}|}\right\rangle_{p(\sigma)} = (2\pi)^{D/2}g_{\text{mean}}\zeta\left\langle\sigma^D\right\rangle_{p(\sigma)} = (2\pi)^{D/2}g_{\text{mean}}\zeta(\beta/\upsilon)^D\frac{\Gamma(\upsilon+D)}{\Gamma(\upsilon)}. \tag{73}$$

And,

$$\langle \mathcal{I}_{n,i,i}(\boldsymbol{s};\sigma)\rangle_{p(\sigma)} = \frac{\pi^{D/2}\left(g_{\text{mean}}^2+\frac{G^2}{3}\right)\zeta}{2\bar{\chi}(s_{\max}-s_{\min})^D}\left\langle\sigma^{D-2}\right\rangle_{p(\sigma)} = \frac{\pi^{D/2}\left(g_{\text{mean}}^2+\frac{G^2}{3}\right)\zeta}{2\bar{\chi}(s_{\max}-s_{\min})^D}\frac{(\beta/\upsilon)^{D-2}\Gamma(\upsilon+(D-2))}{\Gamma(\upsilon)}. \tag{74}$$

Simplifying this expression, we get that in the multimodal, isotropic case,

$$\boxed{\hat{\mathcal{I}}_{M,H,I} = K_D^M\frac{\upsilon^2}{(\upsilon+D-1)(\upsilon+D-2)} \qquad K_D^M = \frac{g_{\text{mean}}+\frac{G^2}{3g_{\text{mean}}}}{2^{D/2+1}\beta^2(s_{\max}-s_{\min})^D}} \tag{75}$$

where as in the unimodal case, $K_D^M$ is dimensionality-dependent (hence the $\cdot_D$ subscript), and we distinguish it from $K_D$ of the unimodal case with the $\cdot^M$ superscript. In the fixed-width limit ($\upsilon\to\infty$), this reduces to

$$\boxed{\hat{\mathcal{I}}_{M,F} = K_D^M.} \tag{76}$$

So that in the multimodal isotropic case, the information gain scales as

$$\boxed{\frac{\hat{\mathcal{I}}_{M,H,I}}{\hat{\mathcal{I}}_{M,F}} = \frac{K_D^M}{K_D^M}\frac{\upsilon^2}{(\upsilon+D-1)(\upsilon+D-2)} = \frac{\upsilon^2}{(\upsilon+D-1)(\upsilon+D-2)}.} \tag{77}$$

Unlike in the unimodal case, the multimodal case appears to predict that heterogeneous widths without shape heterogeneity are increasingly harmful for encoding precision as the dimensionality of the stimulus increases. Indeed, for only $D=1$ is there any information gain achieved with a population of heterogeneously sized, but not heterogeneously shaped, receptive fields. With $D=2$, we already see that, conditioned on having no field shape heterogeneity, a population with heterogeneously sized fields has *lower* information than a comparably tuned fixed-width population:

$$\frac{\hat{\mathcal{I}}_{M,H,I|D=2}}{\hat{\mathcal{I}}_{M,F|D=2}} = \frac{\upsilon^2}{(\upsilon+1)\upsilon} < 1, \quad \forall\upsilon\geq 1. \tag{78}$$

But note that this information gain approaches 1 as $\upsilon\to\infty$. Indeed, the information gain is smallest for $\upsilon=1$. In other words, when there is no shape heterogeneity in the population, the *lowest* encoded information is achieved by populations with maximal width heterogeneity.

**Maximal shape heterogeneity (anisotropic tuning covariance).** Returning to Eq. (72), and re-evaluating $\bar{\chi}$ and the FI terms under the anisotropic assumption:

$$\bar{\chi} = (2\pi)^{D/2} g_{\text{mean}} \zeta \left\langle \sqrt{|\boldsymbol{\Sigma}|} \right\rangle_{p(\sigma)} = (2\pi)^{D/2} g_{\text{mean}} \zeta \left\langle \sigma_i \right\rangle_{p(\sigma)}^{D} = (2\pi)^{D/2} g_{\text{mean}} \zeta \beta^{D}. \tag{79}$$

And,

$$\left\langle \boldsymbol{\mathcal{I}}_{n,i,i}(\boldsymbol{s};\sigma_1,\ldots,\sigma_D) \right\rangle_{p(\sigma_1,\ldots,\sigma_D)} = \frac{\pi^{D/2}\left(g_{\text{mean}}^2 + \frac{G^2}{3}\right)\zeta}{2\bar{\chi}(s_{\max}-s_{\min})^D} \left\langle \frac{1}{\sigma_i} \right\rangle_{p(\sigma_i)} \prod_{j\neq i} \left\langle \sigma_j \right\rangle_{p(\sigma_j)} = \frac{\pi^{D/2}\left(g_{\text{mean}}^2 + \frac{G^2}{3}\right)\zeta\beta^{D-2}}{2\bar{\chi}(s_{\max}-s_{\min})^D} \frac{\upsilon}{\upsilon-1}. \tag{80}$$

Simplifying this expression, we get that in the multimodal, anisotropic case,

$$\boxed{\hat{\mathcal{I}}_{M,H,A} = K_D^M \frac{\upsilon}{\upsilon-1}} \tag{81}$$

where $K_M$ is the same constant as in the multimodal isotropic case. And, again, in the fixed-width limit, $\hat{\mathcal{I}}_{M,F} = K_M$. Then, the information gain scales as

$$\boxed{\frac{\hat{\mathcal{I}}_{M,H,A}}{\hat{\mathcal{I}}_{M,F}} = \frac{K_D^M}{K_D^M}\frac{\upsilon}{\upsilon-1} = \frac{\upsilon}{\upsilon-1}.} \tag{82}$$

As in the unimodal case, the information gain for the anisotropic case is dimensionality independent.

**Continuously varying the shape heterogeneity for $D = 2$.** Returning to Eq. 72 for the case of $D = 2$:

$$\boldsymbol{\mathcal{I}}_{n,1,1}(\boldsymbol{s};\sigma_1,\sigma_2) = \frac{\pi\left(g_{\text{mean}}^2 + \frac{G^2}{3}\right)\zeta}{2\bar{\chi}(s_{\max}-s_{\min})^2}\frac{\sigma_2}{\sigma_1}. \tag{83}$$

and $\bar{\chi} = 2\pi g_{\text{mean}}\zeta \left\langle \sigma_1\sigma_2 \right\rangle_{p(\sigma)}$. We will average over these expressions separately, where, again, we assume in the $D = 2$ case that $\sigma_1, \sigma_2$ are sampled from a correlated bivariate Gamma distribution. Starting with $\bar{\chi}$:

$$\bar{\chi} = 2\pi g_{\text{mean}}\zeta \left\langle \sigma_1\sigma_2 \right\rangle_{p(\sigma_1,\sigma_2)} = 2\pi g_{\text{mean}} \left(\text{Cov}(\sigma_1,\sigma_2) + \left\langle \sigma_1 \right\rangle_{p(\sigma_1)} \left\langle \sigma_2 \right\rangle_{p(\sigma_2)}\right) = 2\pi g_{\text{mean}} \left(\rho\sqrt{\text{Var}(\sigma_1)\text{Var}(\sigma_2)} + \beta^2\right) \tag{84}$$

where the final equality follows from the definition of the correlation coefficient. Then, because $\sigma_1, \sigma_2$ are identically distributed such that $\sqrt{\text{Var}(\sigma_1)\text{Var}(\sigma_2)} = \text{Var}(\sigma_1)$,

$$\bar{\chi} = 2\pi g_{\text{mean}}\zeta \left\langle \sigma_1\sigma_2 \right\rangle_{p(\sigma_1,\sigma_2)} = 2\pi g_{\text{mean}} \left(\frac{\rho\beta^2}{\upsilon} + \beta^2\right) = 2\pi g_{\text{mean}}\beta^2 \left(\frac{\rho}{\upsilon} + 1\right). \tag{85}$$

The remaining terms dependent on $\sigma_1, \sigma_2$ are evaluated in the same way as in the unimodal case, where the simplified expression for the expectation of the ratio of correlated bivariate Gamma variables yields a final encoded information:

$$\boxed{\hat{\mathcal{I}}_{M,H|D=2} = K_2^M \left(\frac{\upsilon(\upsilon-\rho)}{(\upsilon-1)(\upsilon+\rho)}\right).} \tag{86}$$

The fixed-width limit yields:

$$\boxed{\hat{\mathcal{I}}_{M,F|D=2} = K_2^M} \tag{87}$$

such that the information gap in the $2D$ multimodal case is

$$\boxed{\frac{\hat{\mathcal{I}}_{M,H|D=2}}{\hat{\mathcal{I}}_{M,F|D=2}} = \frac{K_2^M}{K_2^M}\left(\frac{\upsilon(\upsilon-\rho)}{(\upsilon-1)(\upsilon+\rho)}\right) = \frac{\upsilon(\upsilon-\rho)}{(\upsilon-1)(\upsilon+\rho)}.} \tag{88}$$

This reflects the intuition results from earlier. Indeed, when $\rho = 1$ (ie., there is no shape heterogeneity), the information gap becomes

$$\frac{\hat{\mathcal{I}}_{M,H|D=2,\rho=1}}{\hat{\mathcal{I}}_{M,F|D=2,\rho=1}} = \frac{\upsilon(\upsilon-1)}{(\upsilon-1)(\upsilon+1)} = \frac{\upsilon}{\upsilon+1} < 1, \quad \forall \upsilon \geq 1. \tag{89}$$

Again, in the fixed-width limit, the information gain becomes 1, suggesting that if there's no shape heterogeneity in the population, any amount of width heterogeneity is harmful for population coding.

Conversely, when $\rho = 0$,

$$\frac{\hat{\mathcal{I}}_{M,H|D=2,\rho=0}}{\hat{\mathcal{I}}_{M,F|D=2,\rho=0}} = \frac{\upsilon}{\upsilon - 1} > 1, \quad \forall \upsilon \geq 1. \tag{90}$$

and indeed, the information gap is unbounded as $\upsilon \to 1$. While the unimodal and multimodal cases make different predictions for dimensionality $D > 2$, both models predict the same field characteristics that maximize encoded information for the $D = 2$ case. Indeed, both models suggest that a population encoding a 2$D$ stimulus can maximize encoded information by maximizing the heterogeneity of both field sizes and shapes across the population.

### 4 Asymptotic encoded information for metabolically constrained populations

We turn now to metabolically constrained populations while leaving the range of tuning centers unbounded. All of the computations above remain the same, except that we here fix the gain to $a_n = |\mathbf{\Sigma}_n|^{-1/2}$ (unlike above, where we assumed $a_n \sim \mathcal{U}[g_{\text{mean}} - G, g_{\text{mean}} + G]$).

#### 4.1 Unimodal tuning curves

As above, we start with averaging over tuning curve centers,

$$\begin{aligned} \langle \mathcal{I}_{n,i,j}(\boldsymbol{s}; \Omega_n) \rangle_{p(\boldsymbol{\mu})} &= \frac{1}{\eta} \left( \frac{1}{(c_{\text{min}} - c_{\text{max}})^D} \int_{c_{\text{min}}}^{c_{\text{max}}} \mathcal{I}_{n,i,j}(\boldsymbol{s}; \Omega_n) d\boldsymbol{\mu} \right) \\ &= \frac{1}{(s_{\max} - s_{\min})^D} \int_{c_{\text{min}}}^{c_{\text{max}}} \frac{1}{\sigma_i^2 \sigma_j^2} (s_i - \mu_i)(s_j - \mu_j) f(\boldsymbol{s}; \Omega_n) d\boldsymbol{\mu} \\ &\underset{\lim_{(c_{\min}, c_{\max}) \to (-\infty, \infty)}}{=} \frac{(2\pi)^{D/2} \Sigma_{i,j}}{\sigma_i^2 \sigma_j^2 (s_{\max} - s_{\min})^D}. \end{aligned} \tag{91}$$

As the tuning curve covariance matrix in the general $D$ case is diagonal, only the diagonal terms of the FIM are non-zero, and are given by

$$\mathcal{I}_{n,i,i}(\boldsymbol{s}; \sigma_1, \cdots, \sigma_D) = \frac{(2\pi)^{D/2}}{\sigma_i^2 (s_{\max} - s_{\min})^D}. \tag{92}$$

Averaging over $\sigma_1, \cdots, \sigma_D$ finally results in

$$\langle \mathcal{I}_{n,i,i}(\boldsymbol{s}; \sigma_1, \cdots, \sigma_D) \rangle_{p(\sigma_1, \cdots, \sigma_D)} = \frac{(2\pi)^{D/2}}{(s_{\max} - s_{\min})^D} \left\langle \frac{1}{\sigma_i^2} \right\rangle_{p(\sigma_1, \cdots, \sigma_D)} = K_D^C \frac{\upsilon^2}{(\upsilon - 1)(\upsilon - 2)}, \tag{93}$$

resulting in

$$\boxed{\hat{\mathcal{I}}_{H,C} = K_D^C \frac{\upsilon^2}{(\upsilon - 1)(\upsilon - 2)}, \qquad K_D^C = \frac{C^D}{\beta^2 (s_{\max} - s_{\min})^D}} \tag{94}$$

where, as in the cases covered above, the constant $K_D^C$ is dimensionality-dependent (hence the $\cdot_D$ subscript), and we use the $\cdot^C$ superscript to distinguish this metabolically constrained case from the constant in the metabolically unconstrained case. In the fixed-width limit, $\upsilon \to \infty$,

$$\boxed{\hat{\mathcal{I}}_{F,C} = K_D^C.} \tag{95}$$

So that the information gain in the metabolically constrained case is

$$\boxed{\frac{\hat{\mathcal{I}}_{H,C}}{\hat{\mathcal{I}}_{F,C}} = \frac{K_D^C}{K_D^C} \frac{\upsilon^2}{(\upsilon - 1)(\upsilon - 2)} = \frac{\upsilon^2}{(\upsilon - 1)(\upsilon - 2)}.} \tag{96}$$

Intriguingly, in the metabolically constrained case, width heterogeneity impacts encoded information, but shape heterogeneity has no impact. Further, the impact of width heterogeneity is independent of stimulus dimensionality.

#### 4.2 Comparing metabolically unconstrained and metabolically constrained populations with matched metabolic costs

Here we ask if metabolically constrained populations (lower gain for wider tuning) feature better encoded information than metabolically unconstrained populations (same mean gain across neurons) if we match their per-neuron metabolic costs.

For tuning function $f(\boldsymbol{s};\Omega_n)$, The average per-neuron metabolic cost is given by $\langle f(\boldsymbol{s};\Omega_n)\rangle_{p(\boldsymbol{s},\boldsymbol{\Omega})}$. For metabolically constrained populations, this cost is by definition $\langle f(\boldsymbol{s};\Omega_n)\rangle_{p(\boldsymbol{s},\boldsymbol{\Omega})} = C^D$. For metabolically unconstrained populations, this cost becomes $\langle f(\boldsymbol{s};\Omega_n)\rangle_{p(\boldsymbol{s},\boldsymbol{\Omega})} = g_{\text{mean}}\,(2\pi)^{D/2}\left\langle\sqrt{|\boldsymbol{\Sigma}_n|}\right\rangle_{p(\Omega_n)}$, where the expectation depends on whether the turning covariance is anisotropic or isotropic.

**Maximal shape heterogeneity (anisotropic tuning covariance).** For an anisotropic tuning covariance we have

$$\left\langle\sqrt{|\boldsymbol{\Sigma}_n|}\right\rangle_{p(\Omega_n)} = \left\langle\sqrt{\prod_{i=1}^{D}\sigma_{n,i}^2}\right\rangle_{p(\Omega_n)} = \prod_{i=1}^{D}\langle\sigma_{n,i}\rangle = \beta^D. \tag{97}$$

such that $\langle f(\boldsymbol{s};\Omega_n)\rangle_{p(\boldsymbol{s},\boldsymbol{\Omega})} = g_{\text{mean}}\,(2\pi)^{D/2}\,\beta^D$. To ensure that this per-neuron metabolic cost equals the one of an metabolically constrained population, $C^D$, we thus require the overall gain mean to scale as

$$g_{\text{mean}} = \left(\frac{C}{\sqrt{2\pi}\beta}\right)^D, \tag{98}$$

with the tuning curve width parameter $\beta$. Substituting this gain into the expression for $\hat{\mathcal{I}}_{H,A}$, Eq. (42), yields

$$\hat{\mathcal{I}}_{H,A} = \frac{C^D\upsilon}{(\upsilon-1)(s_{\text{max}-s_{\text{min}}})^2}\beta^{-2}. \tag{99}$$

Comparing this expression to the metabolically constrained case, Eq. (94), shows that

$$\hat{\mathcal{I}}_{H,E} = \frac{\upsilon}{\upsilon-2}\hat{\mathcal{I}}_{H,A}, \tag{100}$$

that is, for small $\upsilon > 2$, the metabolically constrained population outperforms the metabolically unconstrained one. However, this difference vanishes quickly with decreasing tuning curve width heterogeneity, and $\hat{\mathcal{I}}_{H,E} \to \hat{\mathcal{I}}_{H,A}$ once we approach fixed-width codes, $\upsilon \to \infty$. Therefore, metabolically constrained codes only outperform the encoded information of metabolically unconstrained codes with matched metabolic costs in regimes in which tuning curve sizes are highly heterogeneous, but feature the same encoded information once tuning curve size heterogeneity becomes small.

**No shape heterogeneity (isotropic tuning covariance).** For an isotropic tuning covariance we have

$$\left\langle\sqrt{|\boldsymbol{\Sigma}_n|}\right\rangle_{p(\Omega_n)} = \left\langle\sigma_n^D\right\rangle_{p(\Omega_n)} = \left(\frac{\beta}{\upsilon}\right)^D\frac{\Gamma(\upsilon+D)}{\Gamma(\upsilon)}, \tag{101}$$

where we have again made use of the general expression for the moments of a gamma distribution, such that $\langle f(\boldsymbol{s};\Omega_n)\rangle_{p(\boldsymbol{s},\boldsymbol{\Omega})} = g_{\text{mean}}\,(2\pi)^{D/2}\,(\beta/\upsilon)^D\,\frac{\Gamma(\upsilon+D)}{\Gamma(\upsilon)}$. To again match the metabolic costs of an metabolically constrained code, we now require the neural gain to scale as

$$g_{\text{mean}} = \left(\frac{C\upsilon}{\sqrt{2\pi}\beta}\right)^D\frac{\Gamma(\upsilon)}{\Gamma(\upsilon+D)}, \tag{102}$$

with the tuning curve width parameters, $\beta$ and $\upsilon$. Substituting this gain into the expression for $\hat{\mathcal{I}}_{H,I}$ yields

$$\hat{\mathcal{I}}_{H,I} = \frac{C^D\beta^{-2}}{(s_{\text{max}}-s_{\text{min}})^2}\frac{\upsilon^2\Gamma(\upsilon+(D-2))}{\Gamma(\upsilon+D)} = \frac{C^D\beta^{-2}}{(s_{\text{max}}-s_{\text{min}})^2}\frac{\upsilon^2}{(\upsilon-1)(\upsilon-2)+D(2\upsilon-3+D)}, \tag{103}$$

where we have used $\Gamma(\upsilon+D) = (\upsilon+(D-1))(\upsilon+(D-2))\Gamma(\upsilon+(D-2))$ for the second equality. Comparing this expression to $\hat{\mathcal{I}}_{H,E}$ shows that

$$\hat{\mathcal{I}}_{H,E} = \left(1+\frac{D(2\upsilon-3+D)}{(\upsilon-1)(\upsilon-2)}\right)\hat{\mathcal{I}}_{H,I}. \tag{104}$$

Therefore, as for the anisotropic tuning covariance case, metabolically constrained codes only outperform metabolically unconstrained codes with matched metabolic costs in regimes in which tuning curve sizes are highly heterogeneous, but feature the same encoded information once tuning curve size heterogeneity becomes small.

### 5 Encoded information in populations with bounded tuning centers

Let us now consider bounded populations, in which tuning curve centers can only exceed the stimulus space by finite amounts, which precludes taking the $(c_{\min}, c_{\max}) \to (-\infty, \infty)$ limit. For tractability, we furthermore only consider unimodal tuning curves and two-dimensional stimuli, $D = 2$.

#### 5.1 Metabolically unconstrained populations

As before, we start by averaging over tuning curve centers. Letting

$$\xi_i = \text{erf}\left(\frac{s_i - c_{\min}}{\sqrt{2}\sigma_i}\right) - \text{erf}\left(\frac{c_{\max} - s_i}{\sqrt{2}\sigma_i}\right), \tag{105}$$

$$\lambda_i = \exp\left(\frac{-(s_i - c_{\min})^2}{2\sigma_i^2}\right)(c_{\min} - s_i) + \exp\left(\frac{-(c_{\max} - s_i)^2}{2\sigma_i^2}\right)(s_i - c_{\max}), \tag{106}$$

$$\omega(c_0, c_1) = \exp\left(-\frac{(s_2 - c_0)^2}{2\sigma_2^2} - \frac{(s_1 - c_1)^2}{2\sigma_1^2}\right), \tag{107}$$

we find

$$\begin{aligned}\langle \boldsymbol{\mathcal{I}}_{n,1,1}(\boldsymbol{s}; \Omega_n)\rangle_{p(\boldsymbol{\mu})} &= \frac{1}{\eta}\left(\frac{1}{(c_{\min} - c_{\max})^2}\int_{c_{\min}}^{c_{\max}}\int_{c_{\min}}^{c_{\max}} \boldsymbol{\mathcal{I}}_{n,1,1}(\boldsymbol{s}; \Omega_n) d\mu_1 d\mu_2\right) \\ &= \frac{a\sigma_2\xi_2}{2\sigma_1^2(s_{\max} - s_{\min})^2}\left(\sqrt{2\pi}\lambda_1 + \pi\sigma_1\xi_1\right),\end{aligned} \tag{108}$$

and, similarly,

$$\langle \boldsymbol{\mathcal{I}}_{n,2,2}(\boldsymbol{s}; \Omega_n)\rangle_{p(\boldsymbol{\mu})} = \frac{a\sigma_1\xi_1}{2\sigma_2^2(s_{\max} - s_{\min})^2}\left(\sqrt{2\pi}\lambda_2 + \pi\sigma_2\xi_2\right), \tag{109}$$

$$\begin{aligned}\langle \boldsymbol{\mathcal{I}}_{n,1,2}(\boldsymbol{s}; \Omega_n)\rangle_{p(\boldsymbol{\mu})} &= \langle \boldsymbol{\mathcal{I}}_{n,2,1}(\boldsymbol{s}; \Omega_n)\rangle_{p(\boldsymbol{\mu})} \\ &= \frac{a}{(s_{\max} - s_{\min})^2}\left(\omega(c_{\max}, c_{\max}) - \omega(c_{\max}, c_{\min}) - \omega(c_{\min}, c_{\max}) + \omega(c_{\min}, c_{\min})\right).\end{aligned} \tag{110}$$

Unlike in the unbounded case, each of the FI terms remains stimulus-dependent even after averaging over tuning curve centers. We further simplify these expressions by explicitly considering expectations over stimuli. Assuming that all $s$ are drawn uniformly from the stimulus range,

$$\langle \boldsymbol{\mathcal{I}}_{n,i,j}(\boldsymbol{s}; a, \sigma_1, \sigma_2)\rangle_{p(\boldsymbol{s}|a,\sigma_1,\sigma_2)} = \frac{1}{(s_{\max} - s_{\min})^2}\int_{s_{\min}}^{s_{\max}}\int_{s_{\min}}^{s_{\max}} \boldsymbol{\mathcal{I}}_{n,i,j}(\boldsymbol{s}; a, \sigma_1, \sigma_2) ds_1 ds_2 \tag{111}$$

These integrals can be computed analytically. Let

$$Q(\sigma) = -\exp\left(-\frac{(c_{\min} - s_{\min})^2}{2\sigma^2}\right) + \exp\left(-\frac{(c_{\max} - s_{\min})^2}{2\sigma^2}\right) + \exp\left(-\frac{(c_{\min} - s_{\max})^2}{2\sigma^2}\right) - \exp\left(-\frac{(c_{\max} - s_{\max})^2}{2\sigma^2}\right), \tag{112}$$

$$\begin{aligned}P(\sigma) = &-(c_{\min} - s_{\min})\,\text{erf}\left(\frac{c_{\min} - s_{\min}}{\sqrt{2}\sigma}\right) + (s_{\min} - c_{\max})\,\text{erf}\left(\frac{s_{\min} - c_{\max}}{\sqrt{2}\sigma}\right) \\ &+ (s_{\max} - c_{\min})\,\text{erf}\left(\frac{s_{\max} - c_{\min}}{\sqrt{2}\sigma}\right) - (c_{\max} - s_{\max})\,\text{erf}\left(\frac{c_{\max} - s_{\max}}{\sqrt{2}\sigma}\right),\end{aligned} \tag{113}$$

$$S(\sigma) = \text{erf}\left(\frac{c_{\min} - s_{\min}}{\sqrt{2}\sigma}\right) - \text{erf}\left(\frac{s_{\min} - c_{\max}}{\sqrt{2}\sigma}\right) - \text{erf}\left(\frac{s_{\max} - c_{\min}}{\sqrt{2}\sigma}\right) + \text{erf}\left(\frac{s_{\max} - c_{\max}}{\sqrt{2}\sigma}\right). \tag{114}$$

Then,

$$\boldsymbol{\mathcal{I}}_{n,1,1}(a, \sigma_1, \sigma_2) = \frac{a\sigma_2}{2\sigma_1(s_{\max} - s_{\min})^4}\left[4\sigma_1 Q(\sigma_1) + \sqrt{2\pi}P(\sigma_1)\right]\left[\sigma_2 Q(\sigma_2) + \sqrt{\frac{\pi}{2}}P(\sigma_2)\right], \tag{115}$$

$$\boldsymbol{\mathcal{I}}_{n,2,2}(a, \sigma_1, \sigma_2) = \frac{a\sigma_1}{2\sigma_2(s_{\max} - s_{\min})^4}\left[4\sigma_2 Q(\sigma_2) + \sqrt{2\pi}P(\sigma_2)\right]\left[\sigma_1 Q(\sigma_1) + \sqrt{\frac{\pi}{2}}P(\sigma_1)\right], \tag{116}$$

$$\boldsymbol{\mathcal{I}}_{n,1,2}(a, \sigma_1, \sigma_2) = \boldsymbol{\mathcal{I}}_{n,2,1}(a, \sigma_1, \sigma_2) = -\frac{\pi\sigma_1\sigma_2}{2(s_{\max} - s_{\min})^4}S(\sigma_1)S(\sigma_2). \tag{117}$$

**Recovering the unbounded case.** As a sanity check, let us confirm that we recover the unbounded solution in the $(c_{\min}, c_{\max}) \to (-\infty, \infty)$ limit. In this limit, the only $\{c_{\min}, c_{\max}\}$-dependent functions become $Q(\sigma) \to 0$, $P(\sigma) \to 2(s_{\min} - s_{\max})$, and $S(\sigma) \to 0$, such that

$$\lim_{(c_{\min}, c_{\max}) \to (-\infty, \infty)} \mathcal{I}_{n,1,1}(a, \sigma_1, \sigma_2) = \frac{2\pi a}{(s_{\max} - s_{\min})^2} \frac{\sigma_2}{\sigma_1}, \tag{118}$$

$$\lim_{(c_{\min}, c_{\max}) \to (-\infty, \infty)} \mathcal{I}_{n,2,2}(a, \sigma_1, \sigma_2) = \frac{2\pi a}{(s_{\max} - s_{\min})^2} \frac{\sigma_1}{\sigma_2}, \tag{119}$$

$$\lim_{(c_{\min}, c_{\max}) \to (-\infty, \infty)} \mathcal{I}_{n,1,2}(a, \sigma_1, \sigma_2) = \mathcal{I}_{n,2,1}(a, \sigma_1, \sigma_2) = 0. \tag{120}$$

Averaging over $\sigma_1, \sigma_2$ and $a$ recovers the unbounded, metabolically unconstrained case above.

**A simplifying assumption:** $(c_{\min}, c_{\max}) = (s_{\min}, s_{\max})$**.** The simplifying assumption taken here reflects the simulation paradigm deployed in [1], and simplifies the analytic treatment of the bounded case. With this assumption, $P(\cdot)$, $Q(\cdot)$ and $S(\cdot)$, which we now denote $P_s(\cdot)$, $Q_s(\cdot)$ and $S_s(\cdot)$, become

$$Q_s(\sigma) = 2\left(\exp\left(-\frac{(s_{\max} - s_{\min})^2}{2\sigma^2}\right) - 1\right), \tag{121}$$

$$P_s(\sigma) = 2(s_{\max} - s_{\min}) \,\text{erf}\left(\frac{s_{\max} - s_{\min}}{\sqrt{2}\sigma}\right), \tag{122}$$

$$S_s(\sigma) = 0. \tag{123}$$

Then the FI terms become:

$$\mathcal{I}_{n,1,1}(a, \sigma_1, \sigma_2) = \frac{a\sigma_2}{2\sigma_1(s_{\max} - s_{\min})^4}\left[4\sigma_1 Q_s(\sigma_1) + \sqrt{2\pi} P_s(\sigma_1)\right]\left[\sigma_2 Q_s(\sigma_2) + \sqrt{\frac{\pi}{2}} P_s(\sigma_2)\right], \tag{124}$$

$$\mathcal{I}_{n,2,2}(a, \sigma_1, \sigma_2) = \frac{a\sigma_1}{2\sigma_2(s_{\max} - s_{\min})^4}\left[4\sigma_2 Q_s(\sigma_2) + \sqrt{2\pi} P_s(\sigma_2)\right]\left[\sigma_1 Q_s(\sigma_1) + \sqrt{\frac{\pi}{2}} P_s(\sigma_1)\right], \tag{125}$$

$$\mathcal{I}_{n,1,2}(a, \sigma_1, \sigma_2) = \mathcal{I}_{n,2,1}(a, \sigma_1, \sigma_2) = 0. \tag{126}$$

We finally can take (approximate) expectations over $\sigma_1, \sigma_2$. To do this, we take advantage of the fact that $\sigma_1, \sigma_2 \sim \Gamma(\upsilon, \beta/\upsilon)$ and that they are draw independently. Furthermore, we can rewrite the erf$(\cdot)$ and $\exp(\cdot)$ terms as power series:

$$\text{erf}\left(\frac{1}{x}\right) = \sum_{i=0}^{\infty} \frac{(-1)^i}{i!\,(2i+1)} x^{-2i-1}, \qquad \exp\left(-\frac{1}{x^2}\right) = \sum_{i=0}^{\infty} \frac{(-1)^i}{i!} x^{-2i}. \tag{127}$$

Then, if $x \sim \Gamma(\upsilon, \beta/\upsilon)$ with sufficiently large $\upsilon$,

$$\left\langle x^p \text{erf}\left(\frac{1}{x}\right)\right\rangle_{p(x)} \approx \sum_{i=0}^{\tilde{\upsilon}} \frac{(-1)^i}{i!\,(2i+1)} \beta^{-2i-1+p} \approx \beta^p \text{erf}\left(\frac{1}{\beta}\right) \tag{128}$$

$$\left\langle x^p \exp\left(-\frac{1}{x^2}\right)\right\rangle_{p(x)} \approx \sum_{i=0}^{\tilde{\upsilon}} \frac{(-1)^i}{i!} \beta^{-2i+p} \approx \beta^p \exp\left(-\frac{1}{\beta^2}\right), \tag{129}$$

with $\langle \text{erf}(1/x)\rangle \approx \text{erf}(1/\beta)$ and $\langle \exp(-1/x^2)\rangle \approx \exp(-1/\beta^2)$ as special cases, and where $\tilde{\upsilon}$ is upper limit on the number of terms in the power series that converge, and which is a function of the shape parameter[2] $\upsilon$. With this approximation, marginalizing over $\sigma_1$ and $\sigma_2$ in $\mathcal{I}_{n,1,1}(a, \sigma_1, \sigma_2)$ results in

$$\begin{aligned}
\mathcal{I}_{n,1,1}(a) &= \frac{a}{2(s_{\min} - s_{\max})^4}\left\langle \sigma_1^{-1}\sigma_2\left[4\sigma_1 Q_s(\sigma_1) + \sqrt{2\pi} P_s(\sigma_1)\right]\left[\sigma_2 Q_s(\sigma_2) + \sqrt{\frac{\pi}{2}} P_s(\sigma_2)\right]\right\rangle_{p(\sigma_1,\sigma_2)} \\
&= \frac{a}{2(s_{\min} - s_{\max})^4}\Bigg(4\langle Q_s(\sigma_1)\rangle_{p(\sigma_1)}\langle \sigma_2^2 Q_s(\sigma_2)\rangle_{p(\sigma_2)} + \sqrt{2\pi}\langle \sigma_1^{-1} P_s(\sigma_1)\rangle_{p(\sigma_1)}\langle \sigma_2^2 Q_s(\sigma_2)\rangle_{p(\sigma_2)} \\
&\quad + 2\sqrt{2\pi}\langle Q_s(\sigma_1)\rangle_{p(\sigma_1)}\langle \sigma_2 P_s(\sigma_2)\rangle_{p(\sigma_2)} + \pi\langle \sigma_1^{-1} P_s(\sigma_1)\rangle_{p(\sigma_1)}\langle \sigma_2 P_s(\sigma_2)\rangle_{p(\sigma_2)}\Bigg) \\
&\approx \frac{a}{2(s_{\max} - s_{\min})^4}\left[4\beta Q_s(\beta) + \sqrt{2\pi} P_s(\beta)\right]\left[\beta Q_s(\beta) + \sqrt{\frac{\pi}{2}} P_s(\beta)\right],
\end{aligned} \tag{130}$$

[2]This is a rough approximation, which neglects $\upsilon$ pre-factors obtained with averaging. We verified in simulations that, for sufficiently large $\upsilon$, this is a reasonable approximation.

and, similarly,

$$\boldsymbol{\mathcal{I}}_{n,2,2}(a) \approx \frac{a}{2(s_{\max}-s_{\min})^4}\left[4\beta Q_s(\beta)+\sqrt{2\pi}P_s(\beta)\right]\left[\beta Q_s(\beta)+\sqrt{\frac{\pi}{2}}P_s(\beta)\right], \tag{131}$$

$$\boldsymbol{\mathcal{I}}_{n,1,2}(a) = \boldsymbol{\mathcal{I}}_{n,2,1}(a) = 0. \tag{132}$$

Finally, averaging over the tuning curve gains yields the final form for the bounded tuning curve center average FI per neuron. Using $\hat{B}(\beta) \doteq \left[4\beta Q_s(\beta)+\sqrt{2\pi}P_s(\beta)\right]\left[\beta Q_s(\beta)+\sqrt{\frac{\pi}{2}}P_s(\beta)\right]$, where the function $\hat{B}$ quantifies the influence of boundary effects on the informativeness of a population code as a function of the population's average tuning curve width, we again apply Eq. (1) to find the final expression

$$\boxed{\hat{\mathcal{I}}_{H,B} \approx K_{H,B}\hat{B}(\beta), \quad K_{H,B} = \frac{g_{\text{mean}}}{2(s_{\max}-s_{\min})^4}.} \tag{133}$$

**Limiting cases of $\beta$.** To better understand the impact of boundary effects, let us consider several limiting cases of $\beta$.

**Increasingly narrow tuning.** For increasingly narrow tuning curves, when $\beta \to 0$ (or $\beta^{-1} \to \infty$ in plots), $P_s(\cdot)$ and $Q_s(\cdot)$ become $Q_s(\beta) \to -2$ and $P_s(\beta) \to 2(s_{\max}-s_{\min})$, such that $\hat{B}(\beta) \to 4(s_{\max}-s_{\min})^2$, and

$$\boxed{\lim_{\beta\to 0}\hat{\mathcal{I}}_{H,B} \approx K_{H,B}, \quad K_{H,B} = \frac{2\pi g_{\text{mean}}}{(s_{\max}-s_{\min})^2}} \tag{134}$$

which is equivalent to the metabolically unconstrained, unbounded case above. In other words, as tuning curves become more narrow (particularly relative to the stimulus bounds), boundary effects become negligible.

**Increasingly wide tuning.** For increasingly wide tuning curves, when $\beta \to \infty$ (or $\beta^{-1} \to 0$ in plots), $P_s(\cdot)$ and $Q_s(\cdot)$ become $Q_s(\beta) \to 0$ and $P_s(\beta) \to 0$, such that $\hat{B}(\beta) \to 0$, and

$$\boxed{\lim_{\beta\to\infty}\hat{\mathcal{I}}_{H,B} = 0} \tag{135}$$

which is to say that tuning curves become maximally uninformative in this case. This would align with intuition: Over a bounded tuning curve space, when tuning curves are too wide, all neurons will fire simultaneously, and make stimulus discrimination impossible.

#### 5.2 Metabolically constrained populations

We use the same functions $\zeta_i$, $\lambda_i$, and $\omega(c_0, c_1)$. In the metabolically constrained case,

$$\langle \boldsymbol{\mathcal{I}}_{n,1,1}(\boldsymbol{s};\Omega_n)\rangle_{p(\boldsymbol{\mu})} = \frac{a\xi_2}{2\sigma_1^3(s_{\max}-s_{\min})^2}\left(\sqrt{2\pi}\lambda_1+\pi\sigma_1\xi_1\right), \tag{136}$$

$$\langle \boldsymbol{\mathcal{I}}_{n,2,2}(\boldsymbol{s};\Omega_n)\rangle_{p(\boldsymbol{\mu})} = \frac{a\xi_1}{2\sigma_2^3(s_{\max}-s_{\min})^2}\left(\sqrt{2\pi}\lambda_2+\pi\sigma_2\xi_2\right), \tag{137}$$

$$\begin{aligned}\langle \boldsymbol{\mathcal{I}}_{n,1,2}(\boldsymbol{s};\Omega_n)\rangle_{p(\boldsymbol{\mu})} &= \langle \boldsymbol{\mathcal{I}}_{n,2,1}(\boldsymbol{s};\Omega_n)\rangle_{p(\boldsymbol{\mu})} \\ &= \frac{a}{\sigma_1\sigma_2(s_{\max}-s_{\min})^2}\left(\omega(c_{\max},c_{\max})-\omega(c_{\max},c_{\min})-\omega(c_{\min},c_{\max})+\omega(c_{\min},c_{\min})\right),\end{aligned} \tag{138}$$

and, computing the average over $p(\boldsymbol{s}|a,\sigma_1,\sigma_2)$ (using $P(\cdot)$, $Q(\cdot)$ and $S(\cdot)$ from above):

$$\boldsymbol{\mathcal{I}}_{n,1,1}(a,\sigma_1,\sigma_2) = \frac{a}{2\sigma_1^2(s_{\max}-s_{\min})^4}\left[4\sigma_1 Q(\sigma_1)+\sqrt{2\pi}P(\sigma_1)\right]\left[\sigma_2 Q(\sigma_2)+\sqrt{\frac{\pi}{2}}P(\sigma_2)\right], \tag{139}$$

$$\boldsymbol{\mathcal{I}}_{n,2,2}(a,\sigma_1,\sigma_2) = \frac{a}{2\sigma_2^2(s_{\max}-s_{\min})^4}\left[4\sigma_2 Q(\sigma_2)+\sqrt{2\pi}P(\sigma_2)\right]\left[\sigma_1 Q(\sigma_1)+\sqrt{\frac{\pi}{2}}P(\sigma_1)\right], \tag{140}$$

$$\boldsymbol{\mathcal{I}}_{n,1,2}(a,\sigma_1,\sigma_2) = \boldsymbol{\mathcal{I}}_{n,2,1}(a,\sigma_1,\sigma_2) = -\frac{1}{2}\pi(s_{\max}-s_{\min})^{-4}\left(S(\sigma_1)S(\sigma_2)\right). \tag{141}$$

**Recovering the unbounded case.** As before, we make sure to recover the expressions for the unbounded case in the $(c_{\min}, c_{\max}) \to (-\infty, \infty)$ limit. In this limit we find $Q(\sigma) \to 0$, $P(\sigma) \to 2(s_{\min} - s_{\max})$, and $S(\sigma) \to 0$, such that

$$\lim_{(c_{\min},c_{\max})\to(-\infty,\infty)} \mathcal{I}_{n,1,1}(a, \sigma_1, \sigma_2) = \frac{2\pi a}{(s_{\max} - s_{\min})^2} \frac{1}{\sigma_1^2}, \tag{142}$$

$$\lim_{(c_{\min},c_{\max})\to(-\infty,\infty)} \mathcal{I}_{n,2,2}(a, \sigma_1, \sigma_2) = \frac{2\pi a}{(s_{\max} - s_{\min})^2} \frac{1}{\sigma_2^2}, \tag{143}$$

$$\lim_{(c_{\min},c_{\max})\to(-\infty,\infty)} \mathcal{I}_{n,1,2}(a, \sigma_1, \sigma_2) = \mathcal{I}_{n,2,1}(a, \sigma_1, \sigma_2) = 0. \tag{144}$$

Averaging over $\sigma_1, \sigma_2$ and $a$ again recovers the expressions for the unbounded, metabolically constrained case above.

**A simplifying assumption:** $(c_{\min}, c_{\max}) = (s_{\min}, s_{\max})$

Using the same simplifying assumption as above and $P_s(\cdot)$, $Q_s(\cdot)$ and $S_s(\cdot)$ from the previous section, the FI terms become:

$$\mathcal{I}_{n,1,1}(a, \sigma_1, \sigma_2) = \frac{a}{2\sigma_1^2(s_{\max} - s_{\min})^4} \left[4\sigma_1 Q_s(\sigma_1) + \sqrt{2\pi} P_s(\sigma_1)\right] \left[\sigma_2 Q_s(\sigma_2) + \sqrt{\frac{\pi}{2}} P_s(\sigma_2)\right], \tag{145}$$

$$\mathcal{I}_{n,2,2}(a, \sigma_1, \sigma_2) = \frac{a}{2\sigma_2^2(s_{\max} - s_{\min})^4} \left[4\sigma_2 Q_s(\sigma_2) + \sqrt{2\pi} P_s(\sigma_2)\right] \left[\sigma_1 Q_s(\sigma_1) + \sqrt{\frac{\pi}{2}} P_s(\sigma_1)\right], \tag{146}$$

$$\mathcal{I}_{n,1,2}(a, \sigma_1, \sigma_2) = \mathcal{I}_{n,2,1}(a, \sigma_1, \sigma_2) = 0. \tag{147}$$

The same approximate marginalization over $\sigma_1$, $\sigma_2$ as before then leads to

$$\mathcal{I}_{n,1,1}(a) \approx \frac{a}{2\beta^2(s_{\max} - s_{\min})^4} \left[4\beta Q_s(\beta) + \sqrt{2\pi} P_s(\beta)\right] \left[\beta Q_s(\beta) + \sqrt{\frac{\pi}{2}} P_s(\beta)\right], \tag{148}$$

$$\mathcal{I}_{n,2,2}(a) \approx \frac{a}{2\beta^2(s_{\max} - s_{\min})^4} \left[4\beta Q_s(\beta) + \sqrt{2\pi} P_s(\beta)\right] \left[\beta Q_s(\beta) + \sqrt{\frac{\pi}{2}} P_s(\beta)\right], \tag{149}$$

$$\mathcal{I}_{n,1,2}(a) = \mathcal{I}_{n,2,1}(a) = 0. \tag{150}$$

Finally, averaging with respect to the tuning curve gains results in the final form for the bounded metabolically constrained tuning curve center average FI per neuron:

$$\boxed{\hat{\mathcal{I}}_{H,B,E} \approx K_{H,B,E}\beta^{-2}\hat{B}(\beta), \quad K_{H,B,E} = \frac{g_{\text{mean}}}{2(s_{\max} - s_{\min})^4}.} \tag{151}$$

**Limiting cases of $\beta$.** To again better understand the impact of boundary effects, let us again consider several limiting cases of $\beta$.

**Increasingly narrow tuning.** For $\beta \to 0$, we use the above limits for $P_s(\cdot)$, $Q_s(\cdot)$ and $\hat{B}(\beta)$, leading to

$$\boxed{\lim_{\beta\to 0} \hat{\mathcal{I}}_{H,B,E} \to \infty,} \tag{152}$$

which is equivalent to the metabolically constrained, unbounded case above.

**Increasingly wide tuning.** For $\beta \to \infty$, we again use the above limits for $P_s(\cdot)$, $Q_s(\cdot)$ and $\hat{B}(\beta)$, to find

$$\boxed{\lim_{\beta\to\infty} \hat{\mathcal{I}}_{H,B} = 0,} \tag{153}$$

which is to say that tuning curves become maximally uninformative in this case, similar to the metabolically unconstrained case.

### 6 How our theory relates to Zhang et al. (2023)

Zhang and colleagues have recently shown that the information encoded in neural populations encoding two-dimensional stimuli with unimodal tuning curves is increased when receptive fields exhibit shape heterogeneity [1]. They did so through simulations rather than through an analytical argument. In addition, they focused on a type of heterogeneity that is a special case of our theory. Given that some of their results appear to differ from ours, we devote this section to discussing these differences, and how they can be reconciled with our theory. First, Zhang et al. use a slightly different measure to quantify information. As we show below, this measure differs from ours in that it additionally depends on information variance. However, we also show that this variance-dependent difference vanishes for even moderate population sizes. Second, in contrast to our theory, Zhang et al.'s information appears to grow with population size, even when population-size normalized, and to peak at intermediate average field width sizes. We show that this is a consequence of a combination of assuming bounded tuning centers, and of numerical instabilities in their simulations that caused drops in measured information.

#### 6.1 An alternative measure to quantify information

So far, we have assessed encoded information using the square root determinant of the average population FI matrix, Eq. (1), which, by Eq. (23), equals the single-neuron FI matrix averaged across stochastic tuning curve parameter draws. Here, we ask how this measure relates to an alternative measure that takes the average across tuning curve parameters *after* taking the square root determinant of the FI matrix, as done in [1], rather than *before*, as we have done above. More generally, let $f(\cdot)$ be some nonlinear function, and let us ask how $f(\langle \boldsymbol{\mathcal{I}} \rangle)$ relates to $\langle f(\boldsymbol{\mathcal{I}}) \rangle$, where we here use $\boldsymbol{\mathcal{I}}$ as a shorthand for the encoded information $\boldsymbol{\mathcal{I}}(\boldsymbol{s};\boldsymbol{\Omega})$? We can use a Taylor expansion around $\langle \boldsymbol{\mathcal{I}} \rangle$ up to second order to understand the second quantity:

$$\begin{aligned} \langle f(\boldsymbol{\mathcal{I}}) \rangle &\approx f\left(\langle \boldsymbol{\mathcal{I}} \rangle\right) + \frac{1}{2} \sum_{ij} \sum_{kl} \left\langle \left(\boldsymbol{\mathcal{I}}_{ij} - \langle \boldsymbol{\mathcal{I}}_{ij} \rangle\right) \frac{\partial^2 f(\langle \boldsymbol{\mathcal{I}} \rangle)}{\partial \boldsymbol{\mathcal{I}}_{ij} \partial \boldsymbol{\mathcal{I}}_{kl}} \left(\boldsymbol{\mathcal{I}}_{kl} - \langle \boldsymbol{\mathcal{I}}_{kl} \rangle\right) \right\rangle \\ &= f\left(\langle \boldsymbol{\mathcal{I}} \rangle\right) + \frac{1}{2} \sum_{ij} \sum_{kl} \frac{\partial^2 f(\langle \boldsymbol{\mathcal{I}} \rangle)}{\partial \boldsymbol{\mathcal{I}}_{ij} \partial \boldsymbol{\mathcal{I}}_{kl}} \mathrm{Cov}\left(\boldsymbol{\mathcal{I}}_{ij}, \boldsymbol{\mathcal{I}}_{kl}\right), \end{aligned} \tag{154}$$

where the first-order (gradient) term equals zero, such that only the second-order term playes a role in this expansion. For the case where $f(\cdot) = \sqrt[D]{|\cdot|}$, we find the gradient and Hessian to be given by

$$\frac{\partial f(\boldsymbol{\mathcal{I}})}{\partial \boldsymbol{\mathcal{I}}_{ij}} = \frac{1}{D} f(\boldsymbol{\mathcal{I}}) \mathcal{I}_{ji}^{-1}, \qquad \frac{\partial^2 f(\boldsymbol{\mathcal{I}})}{\partial \boldsymbol{\mathcal{I}}_{ij} \partial \boldsymbol{\mathcal{I}}_{kl}} = \frac{1}{D} f(\boldsymbol{\mathcal{I}}) \left( \frac{1}{D} \boldsymbol{\mathcal{I}}_{ji}^{-1} \boldsymbol{\mathcal{I}}_{lk}^{-1} - \boldsymbol{\mathcal{I}}_{li}^{-1} \boldsymbol{\mathcal{I}}_{jk}^{-1} \right), \tag{155}$$

where we have used $\partial |\boldsymbol{\mathcal{I}}| / \partial \boldsymbol{\mathcal{I}}_{ij} = |\boldsymbol{\mathcal{I}}| \boldsymbol{\mathcal{I}}_{ji}^{-1}$ and $\partial \boldsymbol{\mathcal{I}}_{ji}^{-1} / \partial \boldsymbol{\mathcal{I}}_{kl} = -\mathcal{I}_{jk}^{-1} \boldsymbol{\mathcal{I}}_{li}^{-1}$, and where we use the shorthand notation $\boldsymbol{\mathcal{I}}_{ij}^{-1}$ to denote the $ij$th element of $\boldsymbol{\mathcal{I}}^{-1}$. Substituting this Hessian, evaluated at $\langle \boldsymbol{\mathcal{I}} \rangle$, into Eq. (154) results in

$$\begin{aligned} \left\langle \sqrt[D]{|\boldsymbol{\mathcal{I}}|} \right\rangle &\approx \sqrt[D]{|\langle \boldsymbol{\mathcal{I}} \rangle|} + \frac{\sqrt[D]{|\langle \boldsymbol{\mathcal{I}} \rangle|}}{2D} \sum_{ij} \sum_{kl} \left( \frac{1}{D} \langle \mathcal{I} \rangle_{ji}^{-1} \langle \mathcal{I} \rangle_{lk}^{-1} - \langle \mathcal{I} \rangle_{li}^{-1} \langle \mathcal{I} \rangle_{jk}^{-1} \right) \mathrm{Cov}\left(\boldsymbol{\mathcal{I}}_{ij}, \boldsymbol{\mathcal{I}}_{kl}\right) \\ &= \sqrt[D]{|\langle \boldsymbol{\mathcal{I}} \rangle|} - \frac{\sqrt[D]{|\langle \boldsymbol{\mathcal{I}} \rangle|}}{2D^2} \sum_{ij} \langle \mathcal{I} \rangle_{ii}^{-1} \langle \mathcal{I} \rangle_{jj}^{-1} \left( D \mathrm{Cov}\left(\boldsymbol{\mathcal{I}}_{ij}, \boldsymbol{\mathcal{I}}_{ji}\right) - \mathrm{Cov}\left(\boldsymbol{\mathcal{I}}_{ii}, \boldsymbol{\mathcal{I}}_{jj}\right) \right) \\ &= \sqrt[D]{|\langle \boldsymbol{\mathcal{I}} \rangle|} - \frac{1}{2D^2 \sqrt[D]{|\langle \boldsymbol{\mathcal{I}} \rangle|}} \sum_{ij} \left( D \mathrm{Var}\left(\boldsymbol{\mathcal{I}}_{ij}\right) - \mathrm{Cov}\left(\boldsymbol{\mathcal{I}}_{ii}, \boldsymbol{\mathcal{I}}_{jj}\right) \right). \end{aligned} \tag{156}$$

Here, the second equality is based on observing that $\langle \boldsymbol{\mathcal{I}} \rangle$ is diagonal, such that we can use $\langle \boldsymbol{\mathcal{I}}_{ij} \rangle = \delta_{ij} \boldsymbol{\mathcal{I}}_{ij}$, $\langle \mathcal{I} \rangle_{ij}^{-1} = \delta_{ij} \langle \mathcal{I} \rangle_{ij}^{-1}$, and $\langle \boldsymbol{\mathcal{I}}_{ii} \rangle^{-1} = 1/\langle \boldsymbol{\mathcal{I}}_{ii} \rangle$. The third equality furthermore assumes symmetry across the different stimulus dimensions, which leads to $\langle \mathcal{I} \rangle_{ii}^{-1} = \langle \mathcal{I} \rangle_{11}^{-1}$ for all $i$, such that $\sqrt[D]{|\langle \boldsymbol{\mathcal{I}} \rangle|} = \langle \mathcal{I} \rangle_{11}$, and $\boldsymbol{\mathcal{I}}_{ij} = \boldsymbol{\mathcal{I}}_{ji}$, leading to the final expression.

We can relate these expressions back to our discrimination measure and that used in [1] by noting that, by Eqs. (1) and $(23)$, we have $\hat{\mathcal{I}} = \sqrt[D]{|\langle \boldsymbol{\mathcal{I}} \rangle|}/N$, and that [1] uses $\hat{\mathcal{I}}_Z = \left\langle \sqrt[D]{|\boldsymbol{\mathcal{I}}|} \right\rangle / N$, which we here denote by $\hat{\mathcal{I}}_Z$. Then,

$$\hat{\mathcal{I}}_Z \approx \hat{\mathcal{I}} - \frac{1}{2D^2 \hat{\mathcal{I}}} \sum_{ij} \left( D \text{Var}\left( \frac{\boldsymbol{\mathcal{I}}_{ij}}{N} \right) - \text{Cov}\left( \frac{\boldsymbol{\mathcal{I}}_{ii}}{N}, \frac{\boldsymbol{\mathcal{I}}_{jj}}{N} \right) \right) \tag{157}$$

For $D = 2$, this expression simplifies to

$$\hat{\mathcal{I}}_Z \approx \hat{\mathcal{I}} - \frac{1}{8\hat{\mathcal{I}}} \left( \mathrm{Var}\left( \frac{\boldsymbol{\mathcal{I}}_{11} - \boldsymbol{\mathcal{I}}_{22}}{N} \right) + 4 \mathrm{Var}\left( \frac{\boldsymbol{\mathcal{I}}_{12}}{N} \right) \right), \tag{158}$$

showing that the encoded information measure used by [1] (up to second order) equals our encoded information measure minus some terms that depend on the variance of components of the Fisher information matrix $\boldsymbol{\mathcal{I}}(\boldsymbol{s};\boldsymbol{\Omega})$ of the whole population.

To better understand how these variances scale with population size, we use the law of total covariance to decompose the population FI covariance into

$$\text{Cov}\left(\mathcal{I}_{ij}\left(s;\mathbf{\Omega}\right),\mathcal{I}_{kl}\left(s;\mathbf{\Omega}\right)\right)_{p(s;\mathbf{\Omega})}=\left\langle\text{Cov}\left(\mathcal{I}_{ij}\left(s;\mathbf{\Omega}\right),\mathcal{I}_{kl}\left(s;\mathbf{\Omega}\right)|s\right)_{p(\mathbf{\Omega})}\right\rangle_{p(s)}+\text{Cov}\left(\left\langle\mathcal{I}_{ij}\left(s;\mathbf{\Omega}\right)|s\right\rangle_{p(\mathbf{\Omega})},\left\langle\mathcal{I}_{kl}\left(s;\mathbf{\Omega}\right)|s\right\rangle_{p(\mathbf{\Omega})}\right)_{p(s)}. \tag{159}$$

Due to the tuning curve parameters being independently drawn across neurons from the same distribution, the covariance in the first expectation simplifies to

$$\text{Cov}\left(\mathcal{I}_{ij}\left(s;\mathbf{\Omega}\right),\mathcal{I}_{kl}\left(s;\mathbf{\Omega}\right)|s\right)_{p(\mathbf{\Omega})}=\sum_n\text{Cov}\left(\mathcal{I}_{n,ij}\left(s;\Omega_n\right),\mathcal{I}_{n,kl}\left(s;\Omega_n\right)|s\right)_{p(\Omega_n)}=N\text{Cov}\left(\mathcal{I}_{n,ij}\left(s;\Omega_n\right),\mathcal{I}_{n,kl}\left(s;\Omega_n\right)|s\right)_{p(\Omega)}, \tag{160}$$

where $\mathcal{I}_{n,kl}\left(s;\Omega_n\right)$ denotes the $ij$th element of the FI matrix $\mathcal{I}_n\left(s;\Omega_n\right)$ for neuron $n$, and the second equality follows from all $\Omega_n$ being drawn from the same $p(\Omega)$, such that the covariance is the same across all $n$. This also implies that $\mathcal{I}(s)=\left\langle\mathcal{I}_n\left(s;\Omega_n\right)\right\rangle$ is independent of $n$, such that the second covariance in the decomposition simplifies to

$$\text{Cov}\left(\left\langle\mathcal{I}_{ij}\left(s;\mathbf{\Omega}\right)|s\right\rangle_{p(\mathbf{\Omega})},\left\langle\mathcal{I}_{kl}\left(s;\mathbf{\Omega}\right)|s\right\rangle_{p(\mathbf{\Omega})}\right)_{p(s)}=\text{Cov}\left(N\mathcal{I}_{ij}(s),N\mathcal{I}_{kl}(s)\right)_{p(s)}=N^2\text{Cov}\left(\mathcal{I}_{ij}(s),\mathcal{I}_{kl}(s)\right)_{p(s)}. \tag{161}$$

Overall, when including the $1/N$ normalization, this leads to

$$\text{Cov}\left(\frac{\mathcal{I}_{ij}\left(s;\mathbf{\Omega}\right)}{N},\frac{\mathcal{I}_{kl}\left(s;\mathbf{\Omega}\right)}{N}\right)_{p(s;\mathbf{\Omega})}=\frac{1}{N}\left\langle\text{Cov}\left(\mathcal{I}_{n,ij}\left(s;\Omega_n\right),\mathcal{I}_{n,kl}\left(s;\Omega_n\right)|s\right)_{p(\Omega)}\right\rangle_{p(s)}+\text{Cov}\left(\mathcal{I}_{ij}(s),\mathcal{I}_{kl}(s)\right)_{p(s)}. \tag{162}$$

Substituting this expression back into Eq. (157) leads to

$$\begin{aligned}\hat{\mathcal{I}}_Z\approx\hat{\mathcal{I}}-\frac{1}{2D^2\hat{\mathcal{I}}}\sum_{ij}\Bigg(&\frac{1}{N}\left(D\left\langle\text{Var}\left(\mathcal{I}_{n,ij}\left(s;\Omega_n\right)|s\right)_{p(\Omega)}\right\rangle_{p(s)}-\left\langle\text{Cov}\left(\mathcal{I}_{n,ii}\left(s;\Omega_n\right),\mathcal{I}_{n,jj}\left(s;\Omega_n\right)|s\right)_{p(\Omega)}\right\rangle_{p(s)}\right)\\&+D\text{Var}\left(\mathcal{I}_{ij}(s)\right)_{p(s)}-\text{Cov}\left(\mathcal{I}_{ii}(s),\mathcal{I}_{jj}(s)\right)_{p(s)}\Bigg).\end{aligned} \tag{163}$$

Let us now consider the two cases of unbounded and bounded tuning curve centers separately. For the unbounded population case, the per-neuron FI matrix, $\mathcal{I}(s)$, becomes independent of the value of the stimulus $s$. This implies that both $\text{Var}\left(\mathcal{I}_{ij}(s)\right)_{p(s)}=0$ and $\text{Cov}\left(\mathcal{I}_{ii}(s),\mathcal{I}_{jj}(s)\right)_{p(s)}=0$ for all $i$ and $j$. As a result,

$$\begin{aligned}\hat{\mathcal{I}}_Z&\approx\hat{\mathcal{I}}-\frac{1}{2D^2\hat{\mathcal{I}}}\frac{1}{N}\sum_{ij}\left(D\left\langle\text{Var}\left(\mathcal{I}_{n,ij}\left(s;\Omega_n\right)|s\right)_{p(\Omega)}\right\rangle_{p(s)}-\left\langle\text{Cov}\left(\mathcal{I}_{n,ii}\left(s;\Omega_n\right),\mathcal{I}_{n,jj}\left(s;\Omega_n\right)|s\right)_{p(\Omega)}\right\rangle_{p(s)}\right)\\&=\hat{\mathcal{I}}-\frac{1}{2\hat{\mathcal{I}}}\mathcal{O}\left(\frac{1}{N}\right),\end{aligned} \tag{164}$$

where the second line follows from observing that both variance and covariance in brackets are $\mathcal{O}(1)$. Its $1/N$ pre-factor makes its impact vanish for larger population sizes, such that $\hat{\mathcal{I}}_Z\to\hat{\mathcal{I}}$ once $N$ becomes sufficiently large. As our simulations in Extended Data Fig. 8 show, this is already the case below the $N\approx10^5$ regime of the rat hippocampus.

In the bounded case, the boundary effects cause the per-neuron FI matrix, $\mathcal{I}(s)$, to vary with $s$, such that, unlike for the unbounded case, the last two (co)variance terms in Eq. (163) do not vanish. That is, they contribute $\mathcal{O}(1)$ variances even for large population sizes, such that the relation between $\hat{\mathcal{I}}_Z$ and $\hat{\mathcal{I}}$ becomes

$$\hat{\mathcal{I}}_Z=\hat{\mathcal{I}}-\frac{1}{2\hat{\mathcal{I}}}\left(\mathcal{O}\left(\frac{1}{N}\right)+\mathcal{O}(1)\right). \tag{165}$$

In Extended Data Fig. 8 we show in simulations how the $\mathcal{O}(1/N)$ and $\mathcal{O}(1)$ contribute to $\hat{\mathcal{I}}_Z$. In that figure we furthermore demonstrate that the $\frac{1}{2\hat{\mathcal{I}}}$ pre-factor causes the $\mathcal{O}(1)$ term to only have an insignificant contribution to $\hat{\mathcal{I}}_Z$, such that we observe $\hat{\mathcal{I}}_Z\to\hat{\mathcal{I}}$ with increasing $N$ even in the bounded case.

To get a further intuition for why the measure used in [1] approaches ours for large $N$, consider that the measure $\hat{\mathcal{I}}_Z=\left\langle\sqrt[D]{|\mathcal{I}(s;\mathbf{\Omega})/N|}\right\rangle$ is a function of the normalized population FI matrix, $\mathcal{I}(s;\mathbf{\Omega})/N$, for a given set of population parameters $\mathbf{\Omega}$. Using Eq. (23), this normalized population FI matrix can be re-expressed as

$$\frac{\mathcal{I}(s;\mathbf{\Omega})}{N}=\frac{1}{N}\sum_{n=1}^{N}\mathcal{I}_n(s;\Omega_n), \tag{166}$$

where each neuron's tuning parameters $\Omega_n$ are independently drawn from the same distribution $\Omega_n \sim p(\Omega)$. Thus, the sum over neurons can be interpreted as a Monte Carlo approximation of $\mathcal{I}(s) = \langle \mathcal{I}_n(s;\Omega_n)\rangle_{p(\Omega_n)}$. This approximation becomes precise in the large-$N$ limit, that is

$$\frac{\mathcal{I}(s;\Omega)}{N} \underset{N\to\infty}{\to} \langle \mathcal{I}_n(s;\Omega_n)\rangle_{p(\Omega_n)} = \mathcal{I}(s). \tag{167}$$

This limit reflects the fact that in large populations we are likely to encounter a large diversity of neural tunings that on average well-capture the diversity required to take expectation over tuning curve parameters. This implies that, for fixed $s$, $\mathcal{I}(s;\Omega)/N$ ceases to be a function of $\Omega$ and so ceases to be a random variable. Overall, this implies for large $N$ that

$$\hat{\mathcal{I}}_Z = \left\langle \sqrt[D]{\left|\frac{\mathcal{I}(s;\Omega)}{N}\right|} \right\rangle_{p(s,\Omega)} \approx \left\langle \sqrt[D]{|\mathcal{I}(s)|} \right\rangle_{p(s)} \approx \hat{\mathcal{I}}, \tag{168}$$

where, for the last approximation we have assumed negligible boundary effects, such that $\mathcal{I}(s)$ becomes largely independent of $s$ (note that in the bounded case, this is certainly true in the limit of $\beta^{-1} \to \infty$, and in the unbounded case, this is exactly true). This explains why, in Eq. (158), the variance terms vanish once $N$ becomes sufficiently large.

#### 6.2 How this alternative measure relates to stimulus decoding error

A natural question that arises after comparing our measures is how the measure used by [1] relates to decoding error. We showed in Sec. 2.1.1 that our measure acts as a lower bound on both the mean-squared discrimination error and generalized discrimination error variance. Is the same true for Zhang et al.'s measure? Returning to Eq. (3),

$$\left\langle \sqrt[D]{|\mathrm{Cov}(\epsilon(s;\Omega)|} \right\rangle_{p(s,\Omega)} \geq \left\langle \sqrt[D]{|\mathcal{I}(s;\Omega)^{-1}|} \right\rangle_{p(s,\Omega)} = \left\langle \frac{1}{\sqrt[D]{|\mathcal{I}(s;\Omega)|}} \right\rangle_{p(s,\Omega)} \geq \frac{1}{\left\langle \sqrt[D]{|\mathcal{I}(s;\Omega)|} \right\rangle_{p(s,\Omega)}} \tag{169}$$

where the final inequality follows from Jensen's inequality on the convexity of $f(x) = \frac{1}{x}$. This implies that Zhang et al.'s measure is also a lower bound for the generalized variance, and (by Appendix A.3) thus also a lower bound on the mean-squared discrimination error. However, the relative magnitude of our respective measures imposes consequences for understanding whether or not a positional code is "optimal" with respect to minimizing decoding error.

We can show that, in general, our encoded information measure upper-bounds that used in Zhang et al. It is known that the mapping $X \mapsto \sqrt[D]{|X|}$ is concave over SPD. Then by Jensen's inequality,

$$\sqrt[D]{|\langle \mathcal{I}(s;\Omega)\rangle_{p(s,\Omega)}|} \geq \left\langle \sqrt[D]{|\mathcal{I}(s;\Omega)|} \right\rangle_{p(s,\Omega)}. \tag{170}$$

This implies for its inverses that

$$\frac{1}{\sqrt[D]{|\langle \mathcal{I}(s;\Omega)\rangle_{p(s,\Omega)}|}} \leq \frac{1}{\left\langle \sqrt[D]{|\mathcal{I}(s;\Omega)|} \right\rangle_{p(s,\Omega)}}. \tag{171}$$

In other words, our measure forms a lower bound of the inverse of the measure used by Zhang et al. Interestingly, this implies that the measure used in [1] could in principle be construed as a tighter lower bound on the decoding error than ours. However, as we have show above, the difference vanishes for sufficiently large population sizes (which is approximately achieved in the number of neurons in rat hippocampus), such that in realistic circumstances both of our measurements achieve the same tightness in lower bounding decoding error.

#### 6.3 Why the encoded information in Zhang et al. depends non-monotonically on average receptive field size

In their work, Zhang et al. observed that encoded information depends non-monotonically on the average receptive field size [1, Fig. 4] ($\beta$, in our model). In particular, they found that when decreasing $\beta$ and thus reducing the average receptive field size, encoded information first increases, peaks, and then decreases. Furthermore, they found that the average receptive field size at which information peaks depends on population size, and that peak information grows with the population size, even when normalized by this population size (reproduced in Extended Data Fig. 8b). This seems to conflict with out theory, which, for the maximal shape heterogeneity and $D = 2$ used in [1], predicts by Eq. (42) that encoded information scales as $\beta^{D-2}$, and so becomes independent of both average receptive field size and population size for $D = 2$.

To explain this discrepancy, let us first focus on the initial rise of encoded information when decreasing average field size, $\beta$. This rises is due to Zhang and colleagues assuming that tuning centers lie exclusively within the range of encoded stimuli, whereas our Eq. (42) assumed them to extend significantly beyond this range. Indeed, once we restricted these tuning centers to only lie within the range of encoded stimuli, we found that the encoded information follows Eq. (133) and so becomes dependent on $\beta$, Extended Data Fig. 7. Our theory then predicts that encoded information is larger for smaller average receptive field sizes, thus explaining the initial rise observed by Zhang and colleagues.

We attribute the peak and subsequent drop of information with decreasing $\beta$ to a combination of two additional factors. The first is that Zhang and colleagues used a slightly different measure to quantify information, which we have shown decreases if the information varies more strongly across across differently tuned neurons (Eq. (158), above). However, above we argued that the variance-dependent term vanishes already for moderately sized population. Indeed, this happens asymptotically for large populations, but the rate at which the variance term vanishes is itself dependent on the magnitude of the information variance. This is where Zhang et al.'s specific choice of receptive field heterogeneity becomes important. In particular, the authors assumed field widths to be distributed according to an exponential distribution, which corresponds to a specific case of our theory for which $v \to 1$. This distribution puts higher mass on smaller field widths, and has a mode at field width equal to zero. As a consequence, our theory predicts that encoded information diverges towards infinity when receptive field widths are exponentially distributed. However, Zhang and colleagues did not have analytical expressions for encoded information, and instead computed it through numerical simulations. This implies that they performed Monte Carlo approximations of integrals that do not converge, leading to numerical instabilities. These instabilities in turn significantly increased the variance in information across neurons — a variance that becomes larger for smaller average receptive field sizes $\beta$ (Extended Data Fig. 8e,j). The authors consequently saw that encoded information drops once receptive field sizes decrease further. As we demonstrate in Extended Data Fig. 8, this drop disappears when receptive field width distributions avoid such instabilities.

### A Appendix

#### A.1 Fisher information is additive across independent neurons

Fisher information is additive across neurons for any population of neurons whose stochastic activity is (conditionally on $\boldsymbol{s}$) independent across neurons in the population, that is, where

$$p(\boldsymbol{r}|\boldsymbol{s},\boldsymbol{\Omega}) = \prod_{n=1}^{N} p(r_n|\boldsymbol{s},\Omega_n) \tag{172}$$

holds. To show this, let us use the following, less common, definition of the Fisher information:

$$\boldsymbol{\mathcal{I}}_{i,j}(\boldsymbol{s}) = -\left\langle \left\langle \partial_{s_i}\partial_{s_j} \log p(\boldsymbol{r}|\boldsymbol{s},\boldsymbol{\Omega}) \right\rangle_{p(\boldsymbol{r}|\boldsymbol{s},\boldsymbol{\Omega})} \right\rangle_{p(\boldsymbol{\Omega})}. \tag{173}$$

Then,

$$\begin{aligned} \boldsymbol{\mathcal{I}}_{i,j}(\boldsymbol{s}) &= -\left\langle \left\langle \sum_{n=1}^{N} \partial_{s_i}\partial_{s_j} \log p(r_n|\boldsymbol{s},\Omega_n) \right\rangle_{p(r_n|\boldsymbol{s},\Omega_n)} \right\rangle_{p(\Omega_n)} \\ &= \sum_{n=1}^{N} -\left\langle \left\langle \partial_{s_i}\partial_{s_j} \log p(r_n|\boldsymbol{s},\Omega_n) \right\rangle_{p(r_n|\boldsymbol{s},\Omega_n)} \right\rangle_{p(\Omega_n)} = \sum_{n=1}^{N} \boldsymbol{\mathcal{I}}_{n,i,j}(\boldsymbol{s}), \end{aligned} \tag{174}$$

establishing the required fact. Note that the used definition of Fisher information relies on certain regularity conditions, namely, the existence of the second order derivative of $p(\boldsymbol{r}|\boldsymbol{s},\boldsymbol{\Omega})$. A slightly longer argument (not provided here) can establish the same fact using the more common square-of-gradient Fisher information definition, Eq. (24), without requiring this regularity condition.

#### A.2 Expectations of $\sigma^{-k}$ where $\sigma \sim \Gamma(\cdot)$

Our derivation requires finding $\langle \sigma^{-k} \rangle$ where $\sigma \sim \Gamma(\upsilon, \beta/\upsilon)$ so that $\langle \sigma \rangle = \beta$. We find this expectation using

$$\begin{aligned} \langle \sigma^{-k} \rangle &= \int_0^\infty \sigma^{-k} \left( \frac{1}{\Gamma(\upsilon)(\beta/\upsilon)^{\upsilon}} \sigma^{\upsilon-1} \exp\left(-\frac{\sigma\upsilon}{\beta}\right)\right) d\sigma \\ &= \int_0^\infty \left( \frac{1}{\Gamma(\upsilon)(\beta/\upsilon)^{\upsilon}} \sigma^{\upsilon-k-1} \exp\left(-\frac{\sigma\upsilon}{\beta}\right)\right) d\sigma \\ &= \frac{\Gamma(\upsilon-k)(\beta/\upsilon)^{\upsilon-k}}{\Gamma(\upsilon-k)(\beta/\upsilon)^{\upsilon-k}} \int_0^\infty \left( \frac{1}{\Gamma(\upsilon)(\beta/\upsilon)^{\upsilon}} \sigma^{\upsilon-k-1} \exp\left(-\frac{\sigma\upsilon}{\beta}\right)\right) d\sigma \\ &= \frac{\Gamma(\upsilon-k)(\beta/\upsilon)^{\upsilon-k}}{\Gamma(\upsilon)(\beta/\upsilon)^{\upsilon}} \int_0^\infty \left( \frac{1}{\Gamma(\upsilon-k)(\beta/\upsilon)^{\upsilon-k}} \sigma^{\upsilon-k-1} \exp\left(-\frac{\sigma\upsilon}{\beta}\right)\right) d\sigma \\ &= \frac{\Gamma(\upsilon-k)(\beta/\upsilon)^{\upsilon-k}}{\Gamma(\upsilon)(\beta/\upsilon)^{\upsilon}} = \frac{\upsilon^k \Gamma(\upsilon-k)}{\Gamma(\upsilon)} \beta^{-k} \end{aligned} \tag{175}$$

where the last line follows because

$$\int_0^\infty \left( \frac{1}{\Gamma(\upsilon-k)(\beta/\upsilon)^{\upsilon-k}} \sigma^{\upsilon-k-1} \exp\left(-\frac{\sigma\upsilon}{\beta}\right)\right) d\sigma = \int_0^\infty \Gamma(\upsilon-k, \beta/\upsilon) d\sigma = 1 \tag{176}$$

Note that this expression is undefined unless $\upsilon - k > 0$, and so $\upsilon$ must be chosen accordingly to guarantee that this expectation exists.

#### A.3 The generalized decoding error lower bounds the mean-squared decoding error

In section 2.1.1, we introduce two measures of stimulus decoding error, one based on the trace of the error covariance matrix (the mean-squared decoding error), and one based on the determinant of the error covariance matrix. A natural question is how these two error measures relate to each other.

The error covariance matrix is a symmetric positive semi-definite (SPSD) matrix. As such, one can invoke the arithmetic mean-geometric mean (AM-GM) inequality to relate our two decoding error measurements. We know that for an SPSD matrix $\boldsymbol{X} \in \mathbb{R}^{D\times D}$ that

$$\text{Tr}(\boldsymbol{X}) = \sum_{i=1}^{D} \lambda_i \qquad |\boldsymbol{X}| = \prod_{i=1}^{D} \lambda_i \tag{177}$$

where $\{\lambda_i\}_{i=1}^D$ are the (real, non-negative) eigenvalues of $\boldsymbol{X}$. Then by the AM-GM inequality,

$$\frac{1}{D}\sum_{i=1}^{D}\lambda_i \geq \sqrt[D]{\prod_{i=1}^{D}\lambda_i} \implies \frac{1}{D}\text{Tr}(\boldsymbol{X}) \geq \sqrt[D]{|\boldsymbol{X}|}. \tag{178}$$

Returning to our error covariance matrix, this implies that the generalized decoding error lower bounds the mean-squared decoding error:

$$\frac{1}{D}\text{Tr}(\text{Cov}(\boldsymbol{\epsilon}(\boldsymbol{s};\boldsymbol{\Omega}))) \geq \sqrt[D]{\text{Cov}(\boldsymbol{\epsilon}(\boldsymbol{s};\boldsymbol{\Omega}))} \implies \left\langle \frac{1}{D}\text{Tr}(\text{Cov}(\boldsymbol{\epsilon}(\boldsymbol{s};\boldsymbol{\Omega}))) \right\rangle_{p(\boldsymbol{s},\boldsymbol{\Omega})} \geq \left\langle \sqrt[D]{\text{Cov}(\boldsymbol{\epsilon}(\boldsymbol{s};\boldsymbol{\Omega}))} \right\rangle_{p(\boldsymbol{s},\boldsymbol{\Omega})} \tag{179}$$

#### A.4 The map $X \to |X|^{-1/D}$ is convex for symmetric positive definite matrices

**Lemma 1.** *The map $\boldsymbol{X} \mapsto |\boldsymbol{X}|^{-1/D}$ is convex over the set of symmetric positive definite (*`SPD`*) matrices.*

*Proof.* For our proof, we will first restrict this function to a line (this will simplify our analysis), and show that this restricted function of a single variable is convex, thereby showing that the original function is also convex. Consider the function $h(t) = |\boldsymbol{X} + t\boldsymbol{Y}|^{-1/D}$. It suffices to show that $\forall \boldsymbol{X}, \boldsymbol{Y} \in \texttt{SPD}, t \in [0,1]$ that $h''_{\boldsymbol{X},\boldsymbol{Y}}(t) \geq 0$. We can rewrite $h_{\boldsymbol{X},\boldsymbol{Y}}(t) = f(g_{\boldsymbol{X},\boldsymbol{Y}}(t))$ where $f(a) = a^{-1}$ and $g_{\boldsymbol{X},\boldsymbol{Y}}(t) = |\boldsymbol{X} + t\boldsymbol{Y}|^{1/D}$. For conciseness, denote $\boldsymbol{M}(t) := \boldsymbol{X} + t\boldsymbol{Y}$. The proof follows naturally from the well-known theorem that $g$ is concave over `SPD`:

$$h''_{\boldsymbol{X},\boldsymbol{Y}}(t) = f''(g_{\boldsymbol{X},\boldsymbol{Y}}(t))\left[g'_{\boldsymbol{X},\boldsymbol{Y}}(t)\right]^2 + f'(g_{\boldsymbol{X},\boldsymbol{Y}}(t))g''_{\boldsymbol{X},\boldsymbol{Y}}(t) \tag{180}$$

$$= \frac{2}{D}|\boldsymbol{M}(t)|^{-2/D}\left(\text{Tr}\left(\boldsymbol{M}(t)^{-1}\boldsymbol{Y}\right) - g''_{\boldsymbol{X},\boldsymbol{Y}}(t)\right) \tag{181}$$

Because $g$ is concave, $g''_{\boldsymbol{X},\boldsymbol{Y}}(t)$ is negative $\forall \boldsymbol{X}, \boldsymbol{Y} \in \texttt{SPD}, t \in [0,1]$. Then to conclude the proof, we just need to show that the terms $|\boldsymbol{M}(t)|^{-2/D}$ and $\text{Tr}(\boldsymbol{M}(t)^{-1}\boldsymbol{Y})$ are positive. The first quantity is clearly positive, as $|\boldsymbol{M}(t)|^{-2/D} = \left(|\boldsymbol{M}(t)|^{-1/D}\right)^2$. Then $\text{Tr}(\boldsymbol{M}(t)^{-1}\boldsymbol{Y})$ is positive, because `SPD` is a convex set, so $\boldsymbol{M}(t)$ is `SPD`, and thus $\boldsymbol{M}(t)^{-1}$ is `SPD`, and consequently, $\boldsymbol{M}(t)^{-1}\boldsymbol{Y}$ is `SPD`. Then, the trace, or the sum of its eigenvalues, is positive. □
